# The Neuromusculoskeletal Modeling Pipeline: MATLAB-based Model Personalization and Treatment Optimization Functionality for OpenSim

**DOI:** 10.1101/2024.10.30.620965

**Authors:** C.V. Hammond, S.T. Williams, M.M. Vega, D. Ao, G. Li, R.M. Salati, K.M. Pariser, M.S. Shourijeh, A.W. Habib, C. Patten, B.J. Fregly

**Affiliations:** Rice University; Stanford University; University of California, Davis

## Abstract

Neuromusculoskeletal injuries including osteoarthritis, stroke, spinal cord injury, and traumatic brain injury affect roughly 19% of the U.S. adult population. Standardized interventions have produced suboptimal functional outcomes due to the unique treatment needs of each patient. Strides have been made to utilize computational models to develop personalized treatments, but researchers and clinicians have yet to cross the “valley of death” between fundamental research and clinical usefulness. This article introduces the Neuromusculoskeletal Modeling (NMSM) Pipeline, two MATLAB-based toolsets that add Model Personalization and Treatment Optimization functionality to OpenSim. The two toolsets facilitate computational design of individualized treatments for neuromusculoskeletal impairments through the use of personalized neuromusculoskeletal models and predictive simulation. The Model Personalization toolset contains four tools for personalizing 1) joint structure models, 2) muscle-tendon models, 3) neural control models, and 4) foot-ground contact models. The Treatment Optimization toolset contains three tools for predicting and optimizing a patient’s functional outcome for different treatment options using a patient’s personalized neuromusculoskeletal model with direct collocation optimal control methods. Support for user-defined cost functions and model modification functions facilitate simulation of a vast number of possible treatments.

An NMSM Pipeline use case is presented for an individual post-stroke with impaired walking function, where the goal was to predict how the subject’s neural control could be changed to improve walking speed without increasing metabolic cost. First the Model Personalization toolset was used to develop a personalized neuromusculoskeletal model of the subject starting from a generic OpenSim full-body model and experimental walking data (video motion capture, ground reaction, and electromyography) collected from the subject at his self-selected speed. Next the Treatment Optimization toolset was used with the personalized model to predict how the subject could recruit existing muscle synergies more effectively to reduce muscle activation disparities between the paretic and non-paretic legs. The software predicted that the subject could increase his walking speed by 60% without increasing his metabolic cost per unit time by modifying existing muscle synergy recruitment. This hypothetical treatment demonstrates how NMSM Pipeline tools could allow researchers working collaboratively with clinicians to develop personalized neuromusculoskeletal models of individual patients and to perform predictive simulations for the purpose of designing personalized treatments that maximize a patient’s post-treatment functional outcome.

## 1. Introduction

Approximately 19% of the U.S. adult population is afflicted by a movement impairment caused by stroke, osteoarthritis, traumatic brain injury, limb amputation, cerebral palsy, Parkinson’s disease, or spinal cord injury.^1–3^ In addition to the physical challenges caused by a movement impairment, these individuals are more likely to experience increased health care costs, lower work productivity, a higher risk of developing heart disease or diabetes, a reduction or loss of independence, and a decreased quality of life.^1,4^ At present, treatment of these life-altering impairments typically involves selection of a standardized treatment based on subjective clinical assessment rather than design of a customized treatment based on objective evidence-based methods. Consequently, existing treatments for movement impairments do not address the unique clinical situation and needs of the individual patient to the extent that they could, and should.

Not surprisingly, the standardized treatment paradigm is often unable to achieve the level of post- treatment functional recovery that patients desire and expect. The percentage of patients who are dissatisfied with their functional outcome following a standardized treatment supports this viewpoint.

For those who experience stroke, satisfaction with functional outcome is closely tied to ability to return to a pre-stroke level of movement function.^5^ However, stroke patient satisfaction with mobility improvements generally declines after hospital discharge,^5^ with only 65% of individuals post-stroke recovering the ability to walk.^6^ Even then, they tend to walk at a slow self-selected speed with an asymmetric gait pattern that is less efficient metabolically than for healthy individuals.^6–8^ Furthermore, 20% of stroke survivors remain dependent on others to perform activities of daily living (ADLs).^9^ For those with physician-diagnosed arthritis, roughly 44% have functional limitations that affect their ability to perform ADLs.^10^ For those who receive a total knee replacement (TKR), the best predictor of post- surgery patient satisfaction is functional outcome.^11^ However, 22% of TKR patients are dissatisfied with their functional outcome one year after surgery,^12^ while 16 to 30% are dissatisfied with their ability to perform specific ADLs.^13^ For those who receive a total hip replacement, 7% are dissatisfied with their functional outcome,^14^ with expectations for improved walking and stair climbing ability not being met in over 30% of patients.^15^ For those who receive opening wedge high tibial osteotomy surgery, 22% are dissatisfied with their outcome, primarily due to unmet expectations for their post-surgery ability to perform functional activities.^16^ For those with cerebral palsy (CP), 33% do not improve walking speed as a result of multi-level orthopedic surgery,^17^ while 21% of parents of children with CP do not feel that their child’s orthopedic surgery was worth the time, effort, pain, and cost.^18^ More than 50% of children with CP who undo multi-level surgeries do not see improvements in their gait patterns^19^. For those with lower limb amputation, between 12 and 32% cannot perform desired ADLs with their prosthesis, while only 30% to 52% find the same ADLs very easy to perform.^20^ Those with upper limb amputation use their prosthesis primarily for cosmetic rather than functional reasons, with the percentage of individuals who rate their upper extremity prosthesis as beneficial for performing various ADLs ranging from 5% to only 25%.^21^ These statistics across a myriad of clinical conditions that impair movement highlight the need for a better treatment design paradigm to help a larger percentage of patients recover their desired amount of post-treatment function.

One possible way to improve treatment design and thus functional outcomes for individuals with movement impairments is to employ engineering design optimization methods that utilize physics-based computational models. Physics-based computational models have an advantage over “black box” machine learning models since they extrapolate better to situations outside the boundaries of the training data, which is where optimal solutions often lie. Engineering design optimization methods have dramatically improved function and reliability of commercial products in numerous industries including automotive, aerospace, farming, construction, and structural engineering, among others.^22–27^ Using these virtual prototyping methods, engineers can evaluate countless designs – including highly novel ones – in a time- and cost-efficient manner to determine an optimal design for a given problem, thereby reducing or eliminating trial-and-error design iterations performed on physical prototypes. Researchers have already demonstrated the benefits of combining engineering design optimization with personalized models to develop effective personalized interventions for a small number of clinical problems.^28–33^

While a similar computational approach holds promise for designing personalized treatments for movement impairments, at least three significant challenges must first be overcome for such an approach to become a reality. First, researchers must be able to personalize the neuromusculoskeletal computer models used in the computational treatment design process.^34^ Published musculoskeletal modeling studies have typically used scaled generic models rather than models whose relevant characteristics have been personalized to a subject’s movement data.^34^ This decision is understandable given the challenges of collecting the experimental movement (and possibly imaging) data needed for the model personalization process coupled with the challenges of performing the personalization process itself for any aspect of a model (e.g., joint functional axes). However, differences between patients with the same clinical condition are precisely what make the clinical treatment design process so challenging. Second, researchers must be able to perform the treatment design optimization process using a patient’s personalized neuromusculoskeletal model. Such a process requires performing repeated predictive simulations of patient function following different implementations of a planned intervention. To date, the majority of such simulations have been performed using “home brew” computational software,^35–47^ making it difficult for the broader research community to engage in the process, to repeat studies performed by others, and to transfer knowledge about the process to others. Third, researchers must have access to computational software that makes the model personalization and treatment optimization processes easy, repeatable, and transferable. While recent software advances such as OpenSim Moco^48^ have made it possible to perform predictive simulations more easily, neither OpenSim^49,50^ nor Moco^48^ provide means to personalize relevant aspects of a patient’s neuromusculoskeletal computer model or to use a personalized model to perform the treatment design optimization process.

Several existing musculoskeletal modeling software tools have sought to address these challenges to various degrees.^51–58^ These tools primarily focus on personalizing the musculoskeletal model creation process, including personalizing bone geometry from medical imaging data, personalizing muscle attachment points on bones, and generating scaled generic musculoskeletal models using automated methods. The SimCP simulation platform, designed to predict post-surgery walking function for individuals with cerebral palsy, goes beyond other tools in several important ways.^51^ It supports personalization of Hill-type muscle-tendon model properties, personalization of neural control properties using a muscle synergy structure, and prediction of the joint moment “capability gap” that would remain following implementation of a proposed surgical plan. Nonetheless, none of these tools can personalize multiple aspects of a patient’s neuromusculoskeletal computer model and then use the personalized model within a predictive simulation framework to design clinical interventions.

This article introduces the Neuromusculoskeletal Modeling (NMSM) Pipeline, a MATLAB-based software package that provides researchers and clinicians working together with state-of-the-art computational tools for designing effective personalized treatments for movement disorders. The software package is built on the foundation of the OpenSim musculoskeletal modeling software and adds two toolsets to OpenSim^49,50^. The Model Personalization toolset uses patient movement data and gradient-based optimization to personalize four subcomponents of a patient’s neuromusculoskeletal computer model: 1) joint structure models, 2) muscle-tendon models, 3) neural control models, and 4) foot-ground contact models. The Treatment Optimization toolset uses a patient’s personalized model with direct collocation optimal control methods^35,41,48,59–62^ to predict how the patient’s neural control and/or anatomy should be changed, or how an external device (e.g., an exoskeleton) or implant should be designed and/or controlled, to maximize improvement in the patient’s post-treatment movement function. By emphasizing functionality, ease of use, flexibility, and computational speed, the NMSM Pipeline will hopefully accelerate involvement of the neuromusculoskeletal modeling research community in the computational design of clinically implementable personalized neurorehabilitation and surgical interventions.

The remainder of this article is outlined as follows. First, a design overview of the NMSM Pipeline is presented, including an overview of the philosophy and goals of the software and an introduction to the two included toolsets. Second, implementation details, design decisions, and important information for each tool are presented. Third, an example treatment design use case is provided including specifics about the use of each tool and final results. Fourth, a discussion section is included to provide information about future work, limitations, and access to the software.

## 2. Design Overview

### 2.1 Philosophy and Goals

The goal of the NMSM Pipeline software is to help neuromusculoskeletal modeling researchers engage in clinical research - and cross the valley of death,^63^ thereby moving neuromusculoskeletal modeling a significant step closer to becoming a clinically useful tool. To achieve this goal, the NMSM Pipeline uses a personalized medicine approach, where physics- and physiology-based personalized neuromusculoskeletal computer models constructed from patient movement data are used to design personalized treatments for movement impairments.^64^ Such impairments arise from clinical conditions like osteoarthritis, stroke, cerebral palsy, spinal cord injury, Parkinson’s disease, limb amputation, and even cancer. Because the open-source OpenSim^49,50^ and commercial Anybody^65^ software commonly used for musculoskeletal modeling research lack the ability to personalize neuromusculoskeletal computer models to patient movement data, these software tools cannot currently be used to optimize treatments for individual patients. To overcome this critical limitation, the open-source NMSM Pipeline software uses MATLAB (The MathWorks, Natick MA) to build on the OpenSim foundation and provides two new toolsets - a Model Personalization toolset that facilitates the creation of personalized neuromusculoskeletal computer models, and a Treatment Optimization toolset that uses the personalized models to design personalized treatments that maximize a patient’s post-treatment movement function.

To develop these two new MATLAB-based toolsets, we defined a high-level software design philosophy. First, the NMSM Pipeline was designed to be both accessible for basic users and extensible for advanced users so that neuromusculoskeletal modeling researchers of all levels can benefit from it. Second, modeling and simulation methods were based on laws of physics and principles of physiology and neuroscience rather than on machine learning, since no training data exist for post-treatment movement conditions, and machine learning methods do not, in general, extrapolate well outside the boundaries of the training data. Third, existing OpenSim functionality and model entities were utilized to the fullest extent possible to avoid duplication. Fourth, NMSM Pipeline functionality was designed to have the “look and feel” of OpenSim so that those with experience using OpenSim can quickly become proficient using the Pipeline. Fifth, the NMSM Pipeline was designed to work with any OpenSim model, providing complete flexibility in the musculoskeletal anatomy and movement situations that can be modeled and simulated.

To achieve the goal of accessibility for basic users and extensibility for advanced users, we designed NMSM Pipeline tools to work using Extensible Markup Language (XML) settings files, similar to OpenSim. This approach allows users to run each NMSM Pipeline tool with only a single line of code. For basic users, we developed two OpenSim GUI plugins - one for the Model Personalization toolset with its four tools, and one for the Treatment Optimization toolset with its three tools, where each toolset appears under the OpenSim “Tools” menu. The GUI plugin window for each tool allows users to set up tool runs by configuring the most common settings for each tool (Figure 1), with default values being used automatically for advanced tool settings. Tool settings selected in the GUI plugin, along with default advanced tool settings, can be exported by the user to an XML settings file that can be read back into the GUI and modified as desired, just as with OpenSim tools. For advanced users, advanced tool settings are exposed in each tool’s XML settings file, again similar to OpenSim. Advanced users can create an initial XML settings file using the GUI plugin, or they can start from a pre-existing XML settings file and edit the values of basic and advanced tool settings using any text editor. For both basic and advanced users, each tool is run from the MATLAB command line using a tool-specific function that takes a tool-specific XML settings file as an input. Thus, no “Run” button is provided in any of the GUI plugin tool windows. Tool outputs can be plotted in MATLAB using included utility or custom functions.

**Figure 1:**
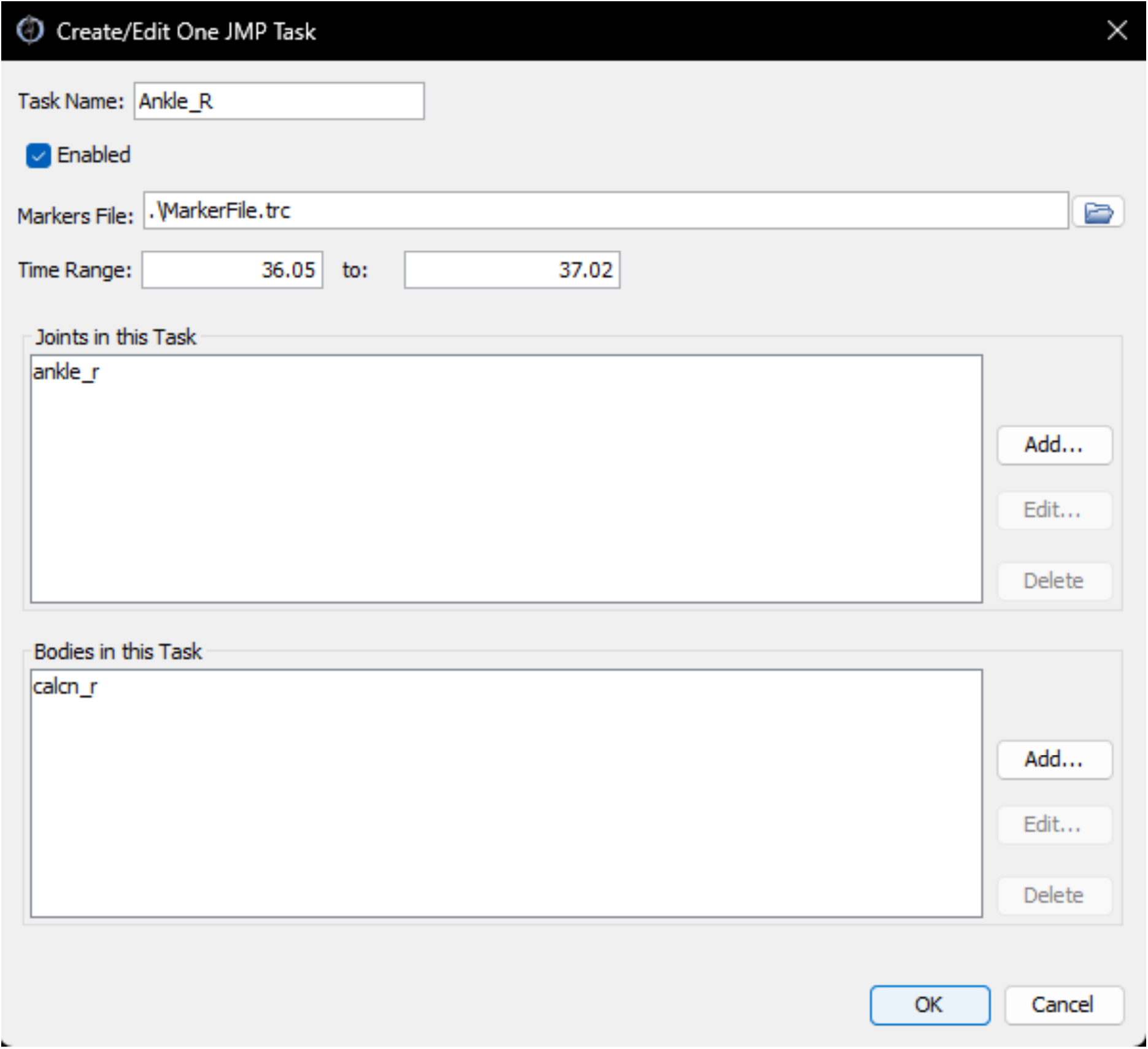
Screenshot of the JMP GUI tool sub-window for adding a JMP task. The GUI tools allow for modifying common settings and creating XML settings files. Advanced settings can be modified directly in the XML file after creation.

Software engineering principles were used for NMSM Pipeline development to facilitate extensibility. Namely, atomic functions - functions that complete only a single operation - were used extensively. All code was written using clear, English-language variable and function names and organized into logical groupings. GitHub’s source control, automated testing framework, and continuous integration tools were used to maintain this extensive project and its documentation throughout the development process. These choices make NMSM Pipeline code easy to understand and easy to modify.

Because NMSM Pipeline tools are based on laws of physics and principles of physiology and neuroscience, we selected computational methods that best facilitate the intended personalized medicine approach. Gradient-based optimization was used for all tools within the NMSM Pipeline primarily for computational speed reasons but also for the ability of gradient-based optimizers to find better solutions than the initial guess, even if the final solution is not the global optimum. For the Model Personalization toolset, built-in MATLAB optimization algorithms (primarily lsqnonlin and fmincon) were used exclusively, while for the Treatment Optimization toolset, the open-source IPOPT optimizer^66^ was used within the commercial GPOPS-II direct collocation optimal control software.^60^ This approach eliminates GPOPS-II and direct collocation optimal control from the Model Personalization process, making model personalization a more reliable and less complex process than it would be if direct collocation optimal control were required.^41,67,68^ The Model Personalization toolset automatically uses MATLAB’s built-in parallel processing capabilities for calculating finite difference derivatives needed by MATLAB optimization algorithms, while the Treatment Optimization toolset automatically uses parallel processing only for repeated OpenSim analyses performed through MATLAB via C++ Mex files.

Through the utilization of OpenSim’s native models and entities, we included a number of convenient facilities in the NMSM Pipeline. OpenSim native file types were used for all data interactions including OpenSim model files (.osim), marker data files (.trc), kinematic motion and ground reaction data (.mot), time series data (.sto), and XML settings files (.xml). Only one new file type was developed-- the NMSM Pipeline model file type (.osimx) to store personalized model properties that are not included in native OpenSim models. Similarly, easy-to-use OpenSim GUI tools such as the Scale tool, Inverse Kinematics (IK) tool, Inverse Dynamics (ID) tool, and Muscle Analysis tool were leveraged as part of the typical pipeline run-through using experimental movement data. For some NMSM Pipeline tools, groups of muscles must be identified, and the muscle groups functionality available in OpenSim models was used for this purpose to avoid the need to store this group information elsewhere.

### 2.2 NMSM Pipeline Toolsets

The NMSM Pipeline is composed of two toolsets-- a Model Personalization toolset and a Treatment Optimization toolset. The Model Personalization toolset personalizes four aspects of a scaled generic OpenSim model using experimental movement data and gradient-based optimization (Figure 2). Those four aspects are 1) joint models, 2) muscle-tendon models, 3) neural control models, and 4) ground contact models. The personalization process is important since personalized rather than generic models are needed to support model-based treatment design for clinical conditions where patients exhibit significant heterogeneity (e.g., stroke).^69,70^ Furthermore, without a personalization process, neuromusculoskeletal models do not reliably predict internal muscle and joint contact forces, body motion, or metabolic cost.^41,71–78^ Each stage in the NMSM Pipeline Model Personalization process has been shown to play an important role in predicting these quantities reliably.^41,71–73,78,79^

**Figure 2:**
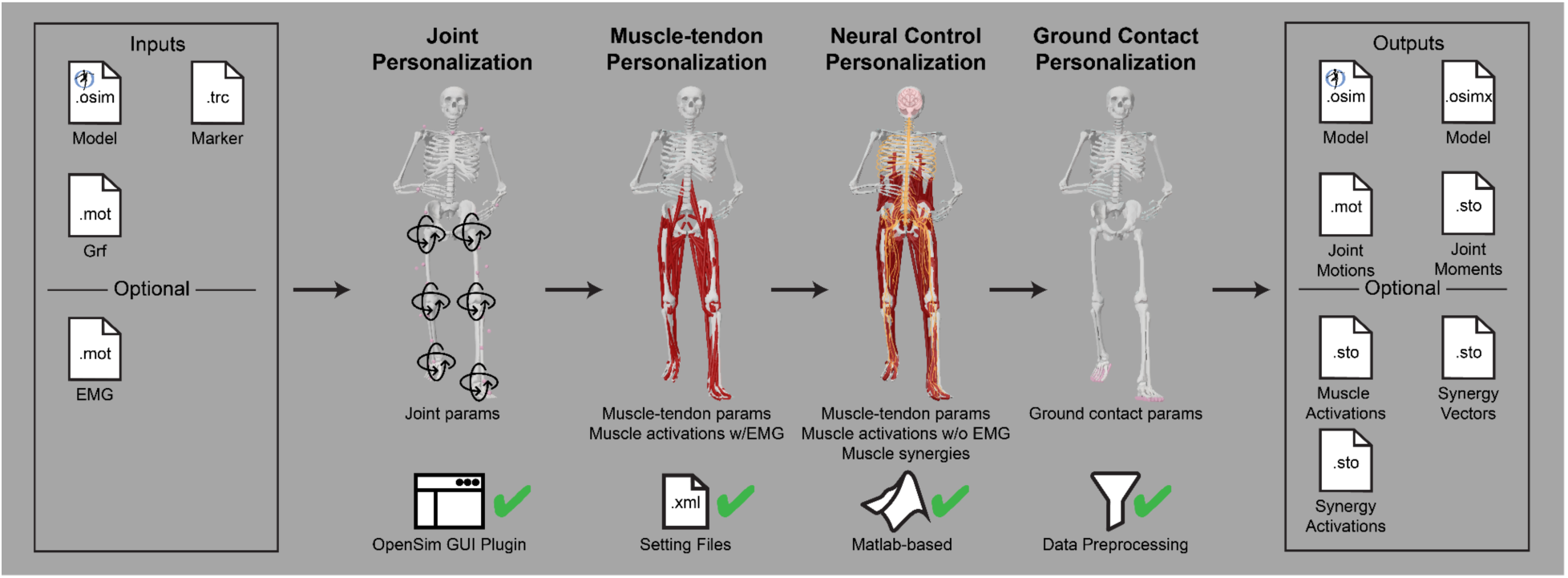
Overview of the Model Personalization toolset. This toolset creates neuromusculoskeletal models with personalized joint, muscle-tendon, neural control, and ground contact parameters through the use of gradient-based optimization.

The Treatment Optimization toolset uses the subject’s personalized neuromusculoskeletal model to predict their post-treatment movement function using direct collocation optimal control (Figure 3). Over the past 10 years, musculoskeletal modeling researchers have converged on direct collocation optimal control as the best method for generating predictive simulations of human movement.^35–37,37,40,41,48,59,67,80–88^ The Treatment Optimization Toolset uses the MATLAB-based GPOPS-II direct collocation optimal control software^60^ to perform predictive simulations of movement for novel conditions where no experimental data exist. Similar to MATLAB, GPOPS-II is inexpensive for academic use. The optimal control solver can adjust neural and/or torque control signals along with neural control, anatomical, internal implant, and/or external device parameters in a patient’s personalized neuromusculoskeletal model to achieve a specified treatment goal (e.g., achieve a desired reduction in metabolic cost). For any patient, this approach can potentially provide the best treatment prescription, the best dosage, and an estimate of the patient’s capacity for improvement.

**Figure 3:**
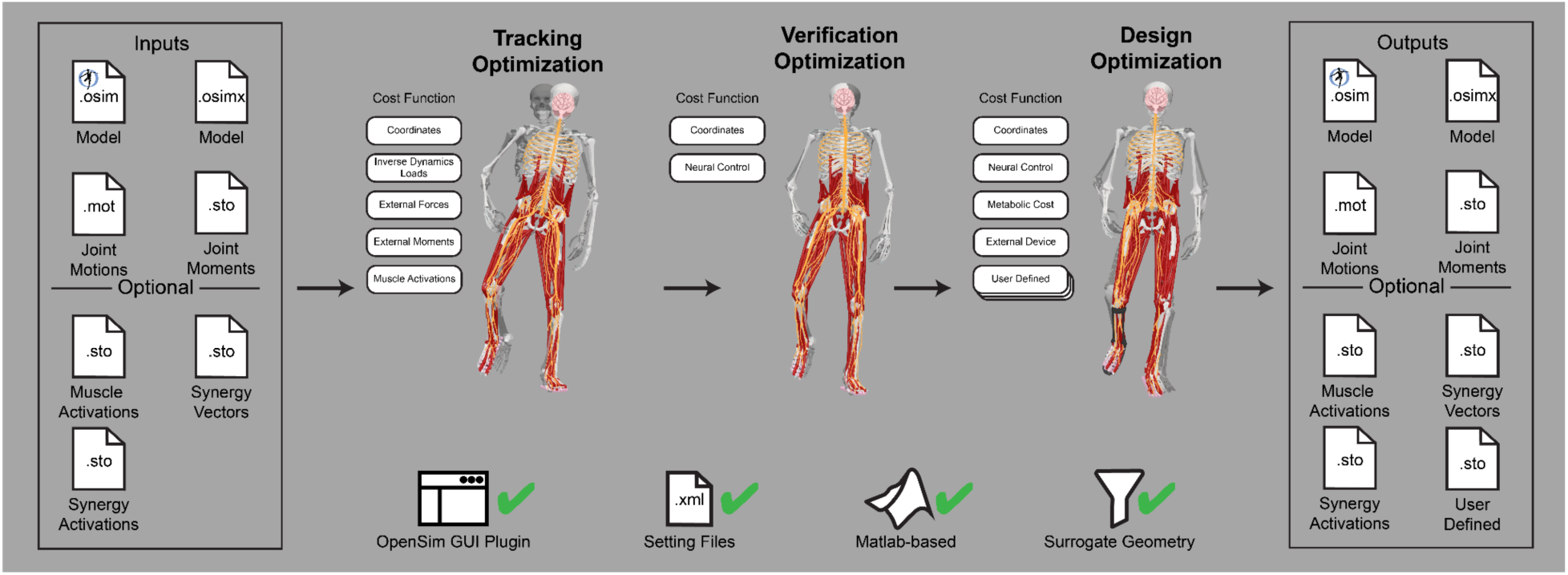
Overview of the Treatment Optimization toolset. This toolset generates a tracking simulation and two subsequent predictive simulations through the use of direct collocation optimal control.

Optimization cost function terms for both toolsets were designed to minimize squared errors and/or cost term values and utilize a common formulation with physically meaningful cost function parameters. Specifically, cost function terms utilize a maximum allowable error and, in some cases, a user-provided error center (Figure 4). For example, if a cost function term seeks to minimize errors between model and experimental joint moments, and matching to within 2 Nm is acceptable, then the user would set their cost term error center to 0 and the associated maximum allowable error to 2. Each cost function term has equal weight with all other cost function terms included in the optimal control problem formulation, where continuous cost function terms involving time series data (e.g., joint angles, joint moments) are normalized by final time so that they do not contribute to the total cost more heavily than do terminal cost function terms involving only a single quantity (e.g., metabolic cost). This method for formulating all cost function terms allows for contextualized decision-making when determining weights for different cost function terms.

**Figure 4:**
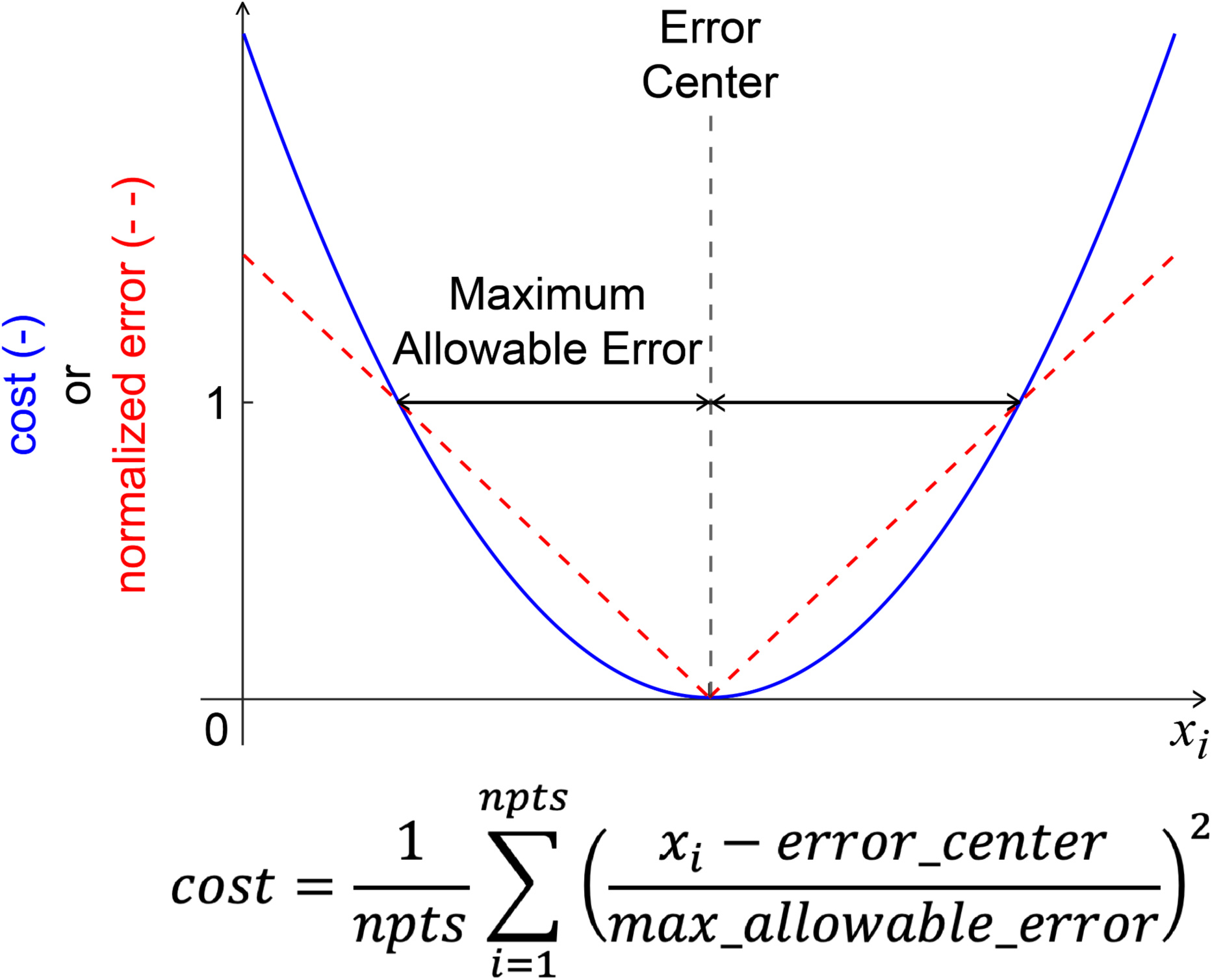
The generic cost function term formulation used throughout the Model Personalization and Treatment Optimization toolsets. The formulation utilizes a maximum allowable error and error center to make the calculated cost physically meaningful. If the difference between the current value and the error center is greater than the maximum allowable error, then the quantity in parentheses will be greater than 1, and squaring this quantity will amplify the cost (e.g., 1.1^2 = 1.21). In contrast, if the difference between the current value and the error center is less than the maximum allowable errors, then the quantity in parentheses will be less than 1, and squaring this quantity will attenuate the cost (e.g., 0.9^2 = 0.81). Some cost function terms track experimental time-varying data, these have an implicit error center of the experimental data value. Some cost function terms minimize errors relative to a static reference value. The type of the cost function term is included in the name for Treatment Optimization cost function terms.

Neural control models for both toolsets are described using muscle synergies. Muscle synergies provide a physiologically-relevant low dimensional representation of a large number muscle activations and can be personalized to represent a specific patient’s neural control strategy.^89–97^ Each muscle synergy is composed of a time-varying synergy activation and an associated time-invariant synergy vector. Each synergy vector contains a weight for each muscle in the model that defines how the associated synergy activation contributes to the total activation of the muscle. Muscle synergies are commonly calculated using non-negative matrix factorization,^89,90^ and typically between three and six muscle synergies are required to account for over 95% of the signal variability for common tasks such as walking, running, and reaching.^98^

## 3. Implementation Details

### 3.1 Model Personalization

The Model Personalization toolset calibrates parameter values in an OpenSim neuromusculoskeletal model so that the model closely reproduces a wide range of experimental movement data. The toolset consists of four tools and was designed so that the tools can be used largely independently from one another, though a preferred tool workflow exists. Personalization is likely to be necessary if the musculoskeletal model is to be used for clinical treatment design purposes.^34^ It also improves models for use in predictive simulations by maximizing the self-consistency of collected experimental data and easing the task of finding a realistic, dynamically consistent motion.

The four tools of the Model Personalization toolset personalize different anatomical, physiological, and neurological aspects of a patient’s OpenSim model. Joint Model Personalization (JMP) calibrates the properties of OpenSim constraint-based joint models, Muscle-tendon Model Personalization (MTP) calibrates the properties of Hill-type muscle-tendon models, Neural Control Model Personalization (NCP) calibrates the properties of muscle synergy control models, and Ground Contact Model Personalization (GCP) calibrates the properties of elastic foundation foot-ground contact models. OpenSim models personalized using the Model Personalization toolset can be used in both the NMSM Pipeline’s Treatment Optimization Toolset and, in the near future, OpenSim Moco. For use with OpenSim Moco, an NMSM Pipeline .osimx model file must be converted to an OpenSim .osim model file using an provided MATLAB model conversion utility program.

The Model Personalization toolset requires input data generated using standard OpenSim tools.

First, a generic musculoskeletal model is scaled using the OpenSim Scale Model tool. To facilitate modeling and simulation of full-body walking motions, we recommend starting with the RCNL2023 OpenSim model (distributed with the NMSM Pipeline software), which is a modified version of the Rajagopal full-body OpenSim model^99^ where the ankle joint kinematic model has been changed to match van den Bogert et al. (1994)^100^ and the knee joint kinematic model changed to eliminate all rotational offsets while producing the same motion. A scaled generic model is one of the required inputs for the JMP tool. After the joints of the model are personalized to the individual’s functional joint axes using the JMP tool, the user must calculate IK motions and ID loads using the OpenSim Inverse Kinematics and Inverse Dynamics tools, respectively. The user must then run the Muscle Analysis tool to calculate muscle-tendon lengths and velocities as well as muscle moment arms if the NMSM Pipeline MTP and NCP tools are to be used. For the JMP tool, marker files can be manually concatenated to personalize joint parameters, values that characterise the transformation of the parent and child bodies of a joint, using multiple motion trials (e.g., marker data from isolated ankle, knee, and hip motion trials can be concatenated with marker data from a walking motion trial to calibrate the functional axes and joint centers of all lower body joints simultaneously). The MTP and NCP tools accept data from one or more motion trials, and the GCP tool accepts data from only a single motion trial, ideally the same trial to be used for Treatment Optimization. If multiple motion trials are used with either the MTP or NCP tools, IK motions, ID loads, muscle-tendon lengths and velocities, and muscle moment arms need to be calculated for each motion trial.

#### 3.1.1 Joint Model Personalization

The Joint Model Personalization tool optimizes joint parameters, body scaling, and marker placement to minimize IK marker distance errors.^100–106^. Reducing inverse kinematics marker distance errors reduces downstream errors in calculated inverse dynamic joint moments^72^, muscle-tendon lengths and velocities, muscle moment arms, and ultimately muscle activations and forces. These quantities are used by subsequent Model Personalization tools.

JMP tool inputs are a scaled generic OpenSim model and marker data from one or more motion trials, where the tool settings file allows the user to specify one or more tasks (Figure 5). The design variables available in a JMP settings file include joint locations and orientations in parent and child bodies, body scale factors applied uniformly, and marker location offsets for each direction independently. The only cost function term is the sum of the squared marker distance errors normalized by the motion trial duration and number of markers. No constraints are included in this optimization problem. The output of JMP is a new OpenSim model with optimized joint parameters, body scale factors, and/or marker placements.

**Figure 5:**
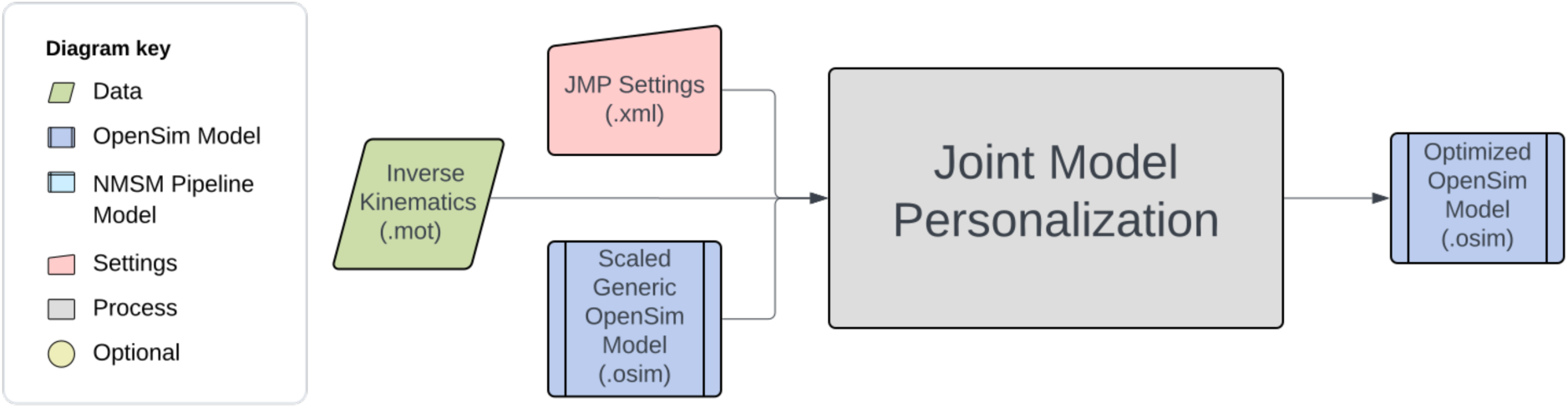
Flowchart showing the inputs to and outputs from the JMP tool. The diagram key also applies to Figs. 6 through 9.

Several important considerations should be kept in mind when using the JMP tool. A JMP run can be split into multiple tasks, each with its own set of design variables and an associated marker file. Careful JMP setup is required to avoid obtaining non-anatomical results. Such a result could be found by personalizing a joint parameter for a coordinate that does not move through sufficient range of motion (> ∼25 degrees) during the motion trial^107^. This issue can be rectified by personalizing the associated joint parameters with separate tasks using isolated joint motion trials, or by concatenating data from an isolated joint trial with data from the desired functional trial (e.g., walking). Another cause of non-anatomic results is selection of joint parameters that produce relative bone motions that are inconsistent with joint function. For example, the parent frame of the hip joint is often defined to be at the center of the acetabulum, while the child frame is often defined to be at the center of the femoral head. Allowing joint parameters to modify the location of the hip joint in either the parent or the child frame could move the center of the femoral head out of the center of the acetabulum, breaking that anatomical relationship. In addition, muscle paths and/or relative body sizes may also become non-anatomical due to inclusion of inappropriate design variables. JMP tool results should be visualized in the OpenSim GUI and problem formulations modified to achieve an anatomically realistic post-JMP model.

#### 3.1.2 Muscle-tendon Model Personalization

The Muscle-tendon Model Personalization tool finds an optimal set of subject-specific muscle-tendon properties and muscle activations from EMG, joint kinematic, and joint moment data by balancing optimization cost function terms related to muscle properties, similarity of properties among grouped muscles, and matching of EMG-driven and experimental inverse dynamics joint moments.^75,76,108,109^ Muscle activation and force predictions are sensitive to optimal muscle fiber length and tendon slack length.^71,110–112^ Therefore, reliable personalization of these parameters is essential for generating reliable predictions of muscle activations and forces during predictive simulations of movement.

The inputs to the MTP tool are a post-JMP OpenSim model as well as IK motion, ID load, muscle-tendon length and velocity, and muscle moment arm data from one or more motion trials of interest (Figure 6). The design variables that can be optimized include activation dynamics and muscle-tendon length parameters for each muscle.^41,75,113^ Cost function terms used by the MTP tool minimize muscle-tendon model parameter deviations from initial values, ID joint moment tracking errors, passive muscle forces, inconsistencies among grouped normalized muscle fiber lengths, inconsistencies among grouped muscle excitation scale factors, and inconsistencies among grouped electromechanical delays. No constraints are applied in this optimization problem. The outputs of the MTP tool are a .osimx model file containing optimized muscle-tendon properties and estimated muscle activations, both for use by other tools in the NMSM Pipeline.

**Figure 6:**
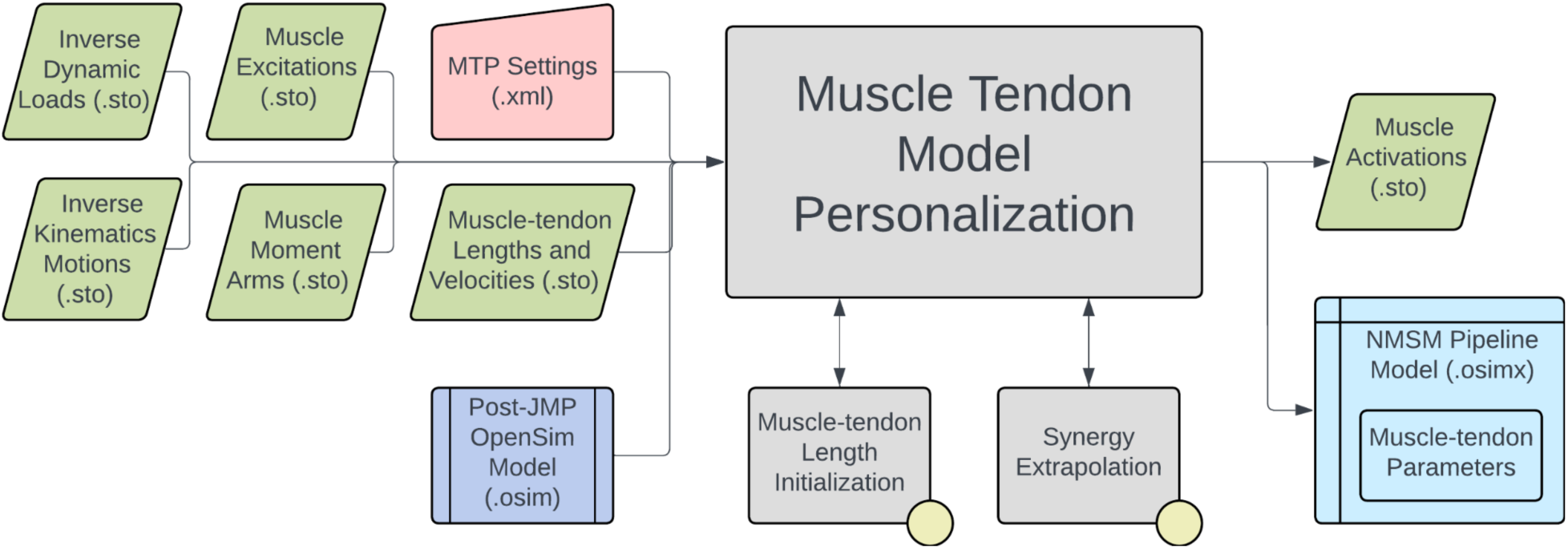
Flowchart showing the inputs to and outputs from the MTP tool. The diagram key is included in Fig. 5.

Several important considerations should be kept in mind when using the MTP tool. Muscle Tendon Length Initialization is included as an option within the MTP tool and is typically used to find initial values for muscle-tendon length parameters that act as reasonable reference values for cost function deviation terms involving muscle-tendon length parameters. Muscle Tendon Length Initialization also estimates the maximum isometric force of each muscle based on published regression relationships.^114^ Since scaling maximum isometric force and muscle excitation at the same time produces a non-unique muscle force solution, and since muscle excitations are scaled using EMG scale factor design variables, maximum isometric force is held constant during an MTP tool run.^71^ Synergy Extrapolation (SynX)^115,116^ is also included in the MTP tool to estimate the activations of muscles without experimental EMG data. SynX uses muscle synergy concepts to estimate missing activations and is more accurate than traditionally used static optimization at predicting missing EMG signals^117^. A typical use case for SynX is estimating the activations of deep muscles from which EMG data cannot be easily collected. SynX can use either principal component analysis or non-negative matrix factorization to calculate the muscle synergies needed to predict muscle activations associated with missing EMG data. The MTP tool uses a Hill-type muscle-tendon model with a rigid tendon and assumes that muscles function primarily on the ascending region of their normalized active force-length curves.^118^ Users can modify cost terms to specify the region of the normalized force-length curve over which muscles are assumed to operate.

Some MTP cost function terms, such as inconsistencies among grouped normalized muscle fiber lengths, calculate the difference between the grouped values relative to their mean. These cost terms are intended to promote similarity in muscle properties within a group under the assumption that similar muscles, such as the three uniarticular vastus muscles, should have similar properties. These muscle groups must be defined within the OpenSim model. Similar cost function terms are also present in NCP.

#### 3.1.3 Neural Control Model Personalization

The Neural Control Model Personalization tool finds muscle synergies that are as consistent as possible with ID joint moments and, when available, MTP-estimated muscle activations. The NCP tool fits muscle synergies at the level of muscle activations (i.e., after electromechanical delay and activation dynamics) for regions of the body where either all muscle activations are available from the MTP tool (e.g., the right lower extremity) or no muscle activations are available.^119,120^ The tool can also be used to find muscle synergies in multiple regions of the body simultaneously - some with and some without previously calculated muscle activations. During the NCP optimization process, grouped muscle properties are maintained and activation-related cost function terms are minimized. Muscle synergies are typically calculated from muscle activations alone using non-negative matrix factorization, and thus the muscle activations reconstructed from the resulting synergies will not closely reproduce ID joint moments when input into a subject’s musculoskeletal model. NCP rectifies this issue by optimizing synergy activations and synergy vectors to be consistent with both ID joint moments and available muscle activations simultaneously.

The inputs to the NCP tool are a post-JMP OpenSim model as well as data for IK motions, ID loads, muscle-tendon lengths and velocities, muscle moment arms, and, optionally, MTP-calculated muscle activations from one or more motion trials of interest (Figure 7). The OpenSim model must contain muscle groups that define lists of muscles for which synergy sets are to be fitted (e.g., all right leg muscles). The design variables for the NCP tool are a set of time-varying synergy activations and an associated set of time-invariant synergy vectors. The cost function terms for the NCP tool minimize ID joint moment matching error, MTP muscle activation matching errors, muscle activations, inconsistencies among grouped normalized muscle fiber lengths, and inconsistencies among grouped muscle activations. The NCP tool includes a constraint that the sum of the muscle weights in each synergy vector must equal a constant times the number of muscles described by that synergy. Upon output, the synergy vectors and synergy activations are normalized such that the largest weight in each synergy vector is one. The NCP tool produces a set of synergy groups inside a new or pre-existing NMSM Pipeline model file (.osimx) as well as data files (.sto) containing synergy activations and their associated synergy vectors.

**Figure 7:**
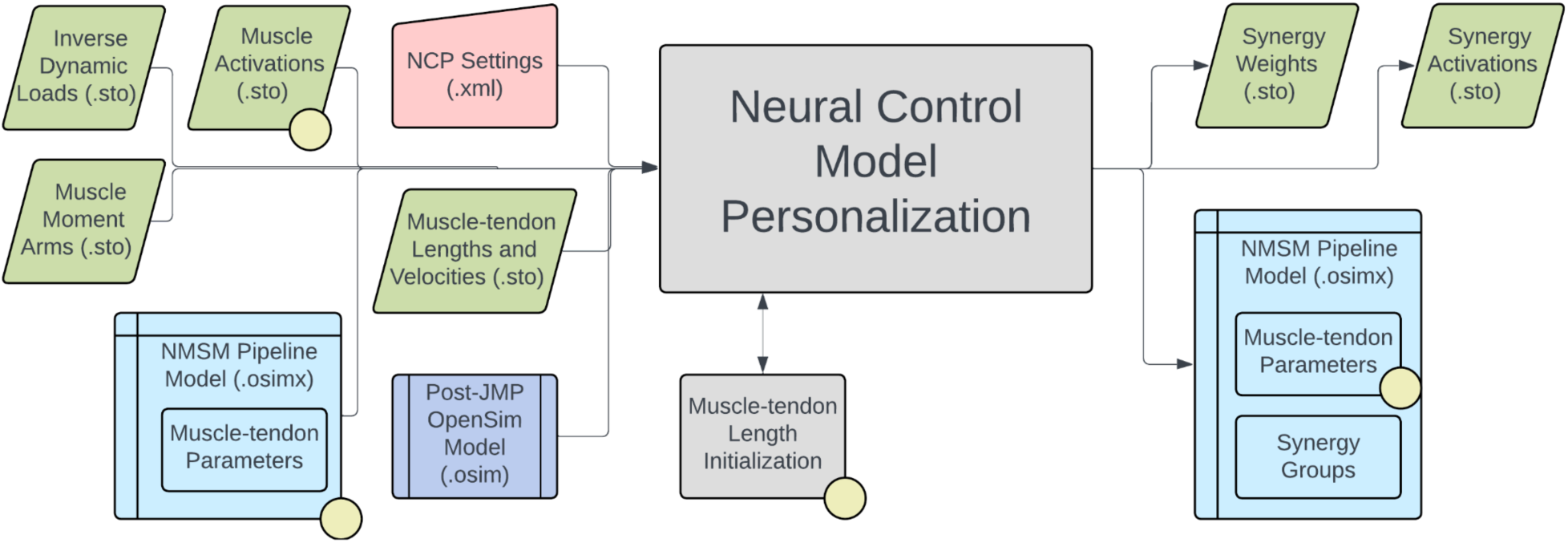
Flowchart showing the inputs to and outputs from the NCP tool. The diagram key is included in Fig. 5.

Several important considerations should be kept in mind when using the NCP tool. Synergy-driven Tracking Optimization (TO), part of the Treatment Optimization toolset, requires that NCP tool results be used as the initial guess. NCP results that distribute errors equally between MTP muscle activation matching and ID joint moment matching will facilitate the process of finding good synergy-driven TO results. In addition, to facilitate obtaining good NCP results, users should perform their own synergy analysis on the muscle activations produced by MTP to determine the optimal number of synergies to use. The number of synergies should be selected to achieve the desired variability accounted for (VAF), which is often set at 90 or 95%.^41,93,121,122^ The NCP tool can be run without having MTP tool results as inputs through the use of the activation minimization cost function term. Since running the NCP tool without MTP tool results produces a non-unique solution, the maximum allowable error of the activation minimization cost function term can be set to achieve a unique solution that contains activations of expected magnitude. Shared synergy vectors between synergy groups can also be enforced by the NCP tool. This feature allows for two muscle synergy groups, such as those for right and left leg muscles, to have nearly identical synergy vectors.

#### 3.1.4 Ground Contact Model Personalization

The Ground Contact Model Personalization tool finds physical properties for elastic foundation foot-ground contact models that closely reproduce experimental ground reaction forces and moments while allowing slight changes in foot kinematics. The elastic foundation is composed of a uniform grid of linear springs with nonlinear damping and friction placed across the bottom of the foot. The tool can personalize ground contact models for each foot separately, where physical properties will differ between the two feet, or both feet together, where physical properties will be the same for the two feet. Currently, the tool works with only a two-segment foot model composed of a hindfoot segment (including calcaneus, cuboid, navicular, cuneiform, and metatarsal bones) and a toes segment (including phalange bones). While generic foot-ground contact models can reproduce experimental ground contact forces accurately, they reproduce experimental ground reaction moments less accurately, introducing errors into the lower body joint moments generated by predictive simulations of walking.^79^ Personalized foot-ground contact models can overcome this issue by reproducing experimental ground reaction moments as well.

The inputs to the GCP tool are a post-JMP lower-body or full-body OpenSim model along with IK motion and associated ground reaction data for a walking trial, where the IK and ground reaction data must possess the same time increments (Figure 8). The ground reaction data must contain three forces, three moments, and a point about which the moments are calculated, which is typically either a time-invariant point fixed in the laboratory or a time-varying center of pressure. The tool automatically strips off both feet from the lower-body or full-body OpenSim model and creates a 6 degrees of freedom (DOF) custom joint connecting each specified hindfoot body to ground. The design variables for the GCP tool are stiffness coefficients unique to each spring with a nonlinear viscous damping coefficient, a dynamic friction coefficient, a viscous friction coefficient, and a spring resting length common to all springs. Additional design variables are B-spline nodal points defining small deviations from experimental foot kinematics with respect to ground. The cost function terms for the GCP tool include minimization of foot marker position errors, foot marker velocity errors, hindfoot and toes segment rotational coordinate errors, hindfoot segment translational coordinate errors, vertical ground reaction force errors, horizontal ground reaction force errors, ground reaction moment errors, and spring coefficient deviations from neighboring spring coefficient values. No constraints are present in this optimization problem. The output of the GCP tool is a new NMSM Pipeline model file (.osimx) containing foot-ground contact model parameter values.

**Figure 8:**
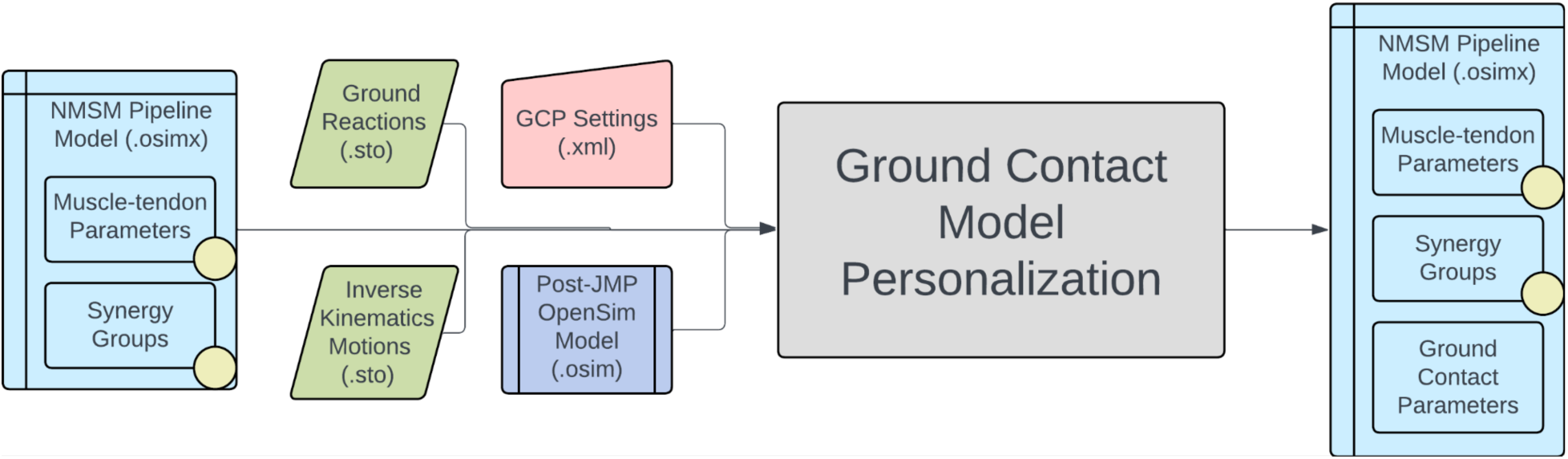
Flowchart showing the inputs to and outputs from the GCP tool. The diagram key is included in Fig. 5.

Several important considerations should be kept in mind when using the GCP tool. The tool can reproduce all six components of ground reactions with good fidelity. To facilitate reproducing ground reaction moments, we use the time-varying projection of a midfoot superior marker onto the ground as the point about which ground reaction moments are calculated. This automatic recalculation of the ground reaction moments helps keep the moments well-scaled during a GCP run, since the moment arm of the ground reaction force vector never becomes excessively large. The center of pressure is not used for joint moment matching since it becomes inaccurate at the transitions into and out of contact due to division by small vertical ground reaction force values. At this time, the GCP tool requires the use of a two-segment foot model with the distal toes segment connected to the hindfoot segment via an oblique pin joint. Before personalization, the GCP tool automatically extracts two-segment OpenSim foot models from the post-JMP lower-body or full-body OpenSim model and automatically calculates hindfoot kinematics with respect to ground that are consistent with the input lower-body or full-body kinematics. Since the foot-ground contact model is sensitive to foot kinematic changes below the measurement accuracy of a markered motion capture system, the GCP tool allows for small changes to the foot motion trajectory to facilitate ground reaction matching. Each foot model possesses seven coordinates: six ground-to-hindfoot coordinates via a 6 DOF custom joint and one toes coordinate via a pin joint.

Coordinate deviations are parameterized with B-spline nodes to reduce the number of design variables. The GCP tool automatically estimates the number of B-spline nodes needed based on a specified lowpass filter cutoff frequency. Another important feature of the GCP tool is its ability to use a different time window for calibrating each foot. This feature facilitates the use of the GCP tool with ground reaction data obtained from labs possessing only two in-ground force plates. The GCP tool also allows for symmetric physical properties between contact surfaces on the two feet, and users can modify the number of springs along each dimension of the grid.

### 3.2 Treatment Optimization

The Treatment Optimization toolset generates predictive simulations of a patient’s post-treatment movement function by optimizing specified treatment design parameters. The toolset consists of three tools designed using a “theme and variation” approach, where the tools are intended to be used in a specific order, with each tool serving a distinct purpose. Each tool uses the GPOPS-II direct collocation optimal control software for MATLAB^60^ and maintains a consistent structure for data inputs, problem design, cost function terms (see Table S1), constraint terms (see Table S2), and outputs, with variations (Figure 9). The Tracking Optimization tool (TO) generates “starting point” predictive simulations of patient movement that are dynamically consistent and closely reproduce all relevant experimental movement data available from the patient (e.g., joint motion, joint moment, ground reaction, and EMG data). The Verification Optimization tool (VO) starts from the Tracking Optimization solution and verifies that the controls found produce the correct motion without tracking the motion directly, which ensures that the subsequent treatment design problem is well formulated before simulated treatments are explored. Finally, the Design Optimization tool (DO) starts from the Verification Optimization solution, applies and (if desired) optimizes simulated treatments, and predicts the patient’s resulting post- treatment movement along with associated controls and loads. All three tools work with torque controlled, muscle synergy controlled, and combined torque and muscle synergy controlled problem formulations.

**Figure 9:**
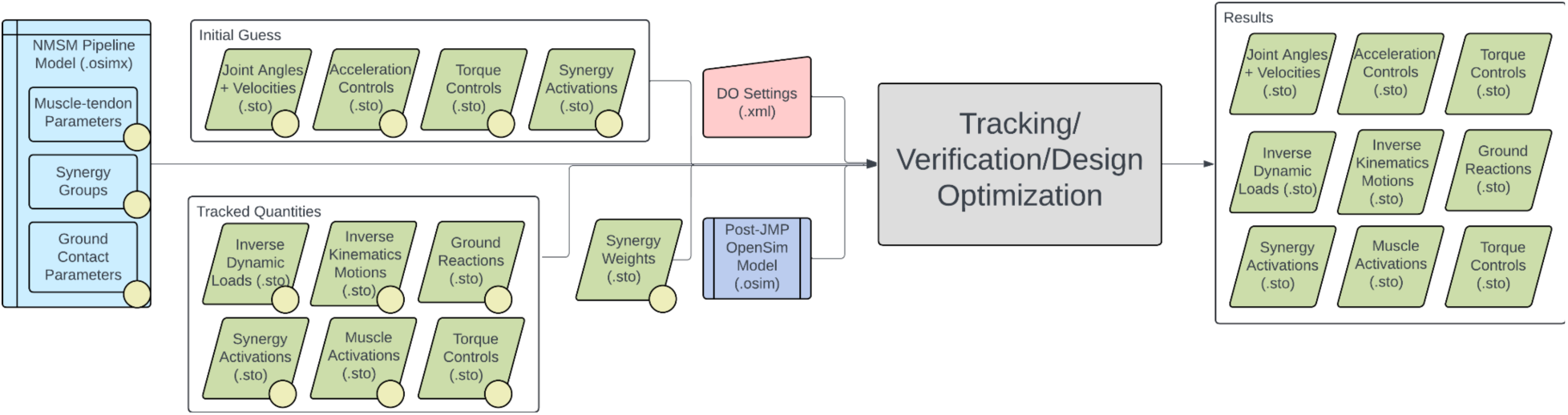
Flowchart showing the inputs to and outputs from the tools included in the Treatment Optimization toolset. Since these tools follow a theme and variation approach, various data combinations can be included or excluded based on the goals of the user. The diagram key is included in Fig. 5.

Several important considerations should be kept in mind when working with the Treatment Optimization toolset. Direct collocation optimal control problems are solved by modifying states and controls simultaneously to minimize an objective function and fulfill constraints. For Treatment Optimization, the states consist of joint positions and velocities for selected OpenSim model coordinates and the controls consist of corresponding joint accelerations (due to the use of an implicit dynamics formulation^59^), torque controls, synergy controls, or both. All Treatment Optimization tools accept both a “tracked quantities” and an “initial guess” directory. The tracked quantities directory contains reference data (e.g., experimental joint positions, joint moments, muscle activation, and ground reactions) to be tracked by specified cost function terms while the initial guess directory contains data specifying the initial guess for the problem’s states and controls. Treatment Optimization XML settings files allow for joint position, velocity, and acceleration range scale factors as well as muscle synergy and torque controller range scale factors. These range scale factors describe the search space of the optimal control problem by confining the states and controls to be within a certain range relative to their initial range (see Figure 10). After a completed run, all Treatment Optimization tools output the same files to a user-specified results directory. Specifically, the solution’s states and controls, along with IK motions, ID loads, and ground reactions calculated from the solution’s states and controls, are output to the specified directory. Saved states and controls can then be passed to the next Treatment Optimization tool as an initial guess to pick up where the previous tool left off.

**Figure 10:**
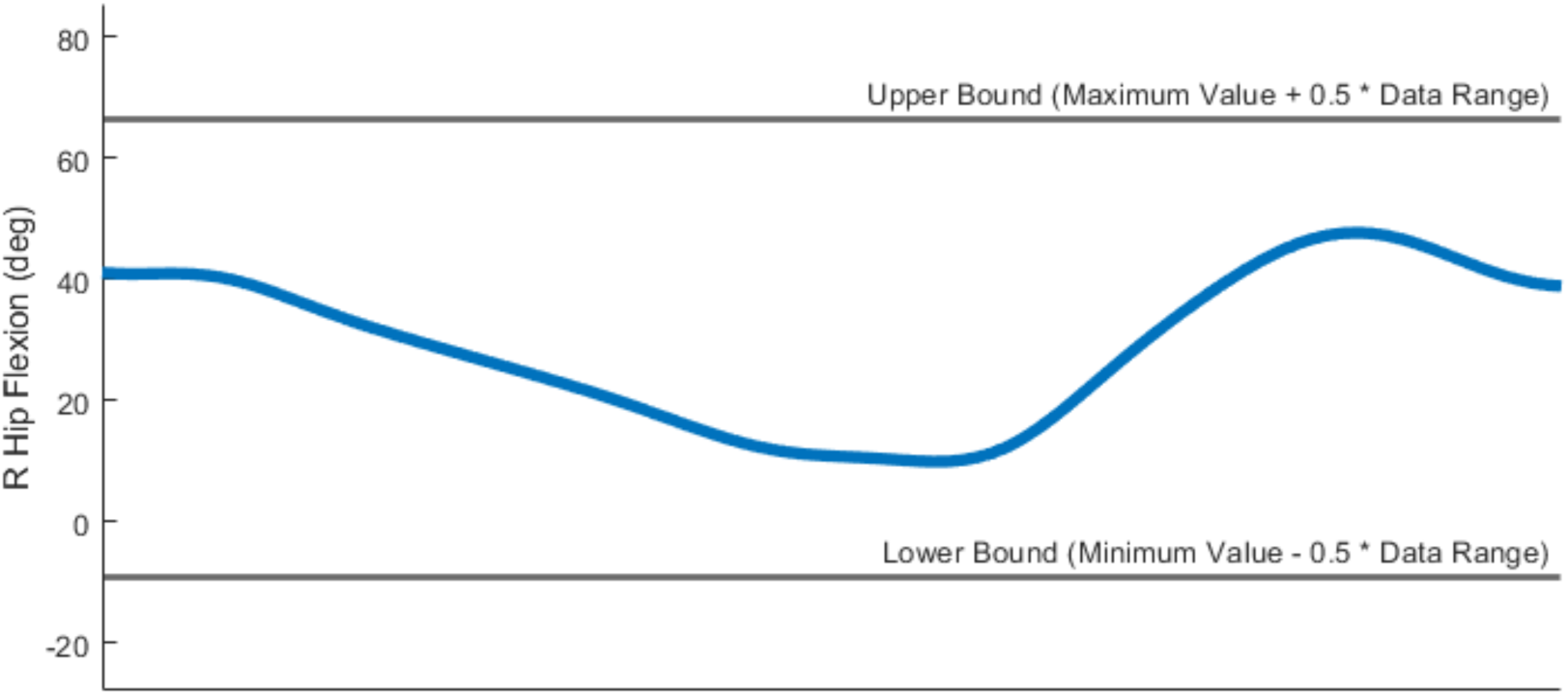
Illustration showing how range scale factors are used to set bounds on the states and controls within GPOPS-II. The range scale factor value is used to set the maximum and minimum searchable spaces for the states and controls as offsets beyond the initial range. In this figure, a range scale factor of 0.5 was used.

Because Treatment Optimization tools use implicit dynamics,^41,59^ several constraint terms must be included for successful use of the toolset. One constraint is that the joint acceleration controls must be the first time derivatives of the joint velocity states, and the joint velocity states must be the first time derivatives of the joint position states. These constraints are handled internally by GPOPS-II. Another constraint is that the net joint loads produced by the torque and synergy controls must be consistent with the model’s inverse dynamic loads. This constraint must be handled by the user by including kinetic consistency constraint terms for all controlled coordinates included in the problem formulation. In addition, if users want their Treatment Optimization solutions to be dynamically consistent, then the root segment residual load constraint must be included to ensure that the residual loads acting on the model’s root segment (i.e., the segment connecting the model to ground) are close to zero. In addition, if the motion of interest is expected to be periodic, such as for walking, running, cycling, or stair-climbing, joint position and velocity and ground reaction force and moment periodicity constraints can be used to ensure that a near-periodic motion is found.

#### 3.2.1 Tracking Optimization

The Tracking Optimization tool uses a personalized model to produce a dynamically consistent movement simulation that closely reproduces all available experimental motion data, including joint motions, joint moments, ground reaction forces and moments, and muscle activations. To achieve a dynamically consistent motion, the tool spreads out matching errors between the different experimental quantities based on user-specified maximum allowable errors. The tool accepts a post-JMP OpenSim model (.osim file) and personalized NMSM Pipeline model (.osimx file) along with experimental IK motions, ID loads, ground reactions, muscle-tendon lengths and velocities, muscle moment arms, and, if using synergy controls, NCP results for the trial of interest. The following cost function terms are commonly included to achieve a successful Tracking Optimization run: generalized coordinate tracking, generalized speed tracking, inverse dynamic moment tracking, muscle activation tracking, external force tracking, and external moment tracking. Common constraint terms include kinetic consistency, state position periodicity, state velocity periodicity, and root segment residual load bounding.

Several important considerations should be kept in mind when using the Tracking Optimization tool. Finding a good result with the Tracking Optimization tool is the biggest challenge in the NMSM Pipeline. The goal is to trade off errors between tracked experimental data sources by modifying the maximum allowable error in each cost function term based on the importance of and confidence in the respective experimental data. Higher quality Model Personalization results and highly consistent tracked experimental quantities (e.g., tracked muscle activations should closely reproduce tracked ID moments) make finding a good Tracking Optimization solution easier.

#### 3.2.2 Verification Optimization

The Verification Optimization tool is used to perform a “dry run” Design Optimization without including any “design elements”, the goal being to ensure a good initial guess and to verify the appropriateness of the optimal control problem formulation. The tool accepts the same inputs as the TO tool, but in general, the results directory of a TO run is used for both the initial guess and tracked quantities. The cost function terms for the VO tool include controller tracking and, for uncontrolled joints, generalized coordinate tracking. Similar to the TO tool, the VO tool includes constraint terms for kinetic consistency, state position periodicity, state velocity periodicity, and root segment residual load bounding. If a VO problem is well formulated, it should converge quickly since the initial guess is already at the optimal solution.

Several important considerations should be kept in mind when using the Verification Optimization tool. Users must decide whether each coordinate should be controlled or tracked. If controlled, the coordinate should be assigned the same controller (synergy, torque, or both) as in the TO tool, and the controls should be tracked by the VO tool. Conversely, if the coordinate does not have a controller or has a non-essential controller, then the coordinate should be tracked by the VO tool. An example of a non-essential controller is a torque controller on an upper body joint for a full body gait model with only lower body muscles. The results of a VO run should closely match the outputs of the TO run used to define the initial guess and tracked quantities. If the results do not match closely, then a problem formulation issue is likely present, and users should review their TO and VO settings. Examples of potential problem formulation issues are having a generalized coordinate that is neither controlled nor tracked, having a generalized coordinate that is both controlled and tracked, or having a generalized coordinate that is tracked with a different maximum allowable error than was used by TO. Each of these situations can produce a VO solution that differs from the TO initial guess. After an adequate VO solution is found, users can move on to the DO tool by adding a “treatment” to their VO problem formulation.

#### 3.2.3 Design Optimization

The Design Optimization tool predicts or optimizes how a planned treatment will affect a patient’s post-treatment movement function. The tool accepts the same inputs as the other Treatment Optimization tools, but typically VO results are used as the initial guess. Similar to VO, DO allows for controller tracking and generalized coordinate tracking alongside other DO-specific cost function terms. Similar to TO and VO, DO can use kinetic consistency, state position periodicity, state velocity periodicity, and root segment residual load constraint terms as well as other constraints to facilitate the design of a personalized intervention.

Several important considerations should be kept in mind when using the Design Optimization tool. Unlike the other Treatment Optimization tools, the DO tool allows for both fixed and free final time problem formulations. For free final time problems, the time vector can be constrained to a user defined range and tracked control quantities are stretched or compressed in time to match the current time vector. A number of built-in cost function terms have been included in the DO tool. These terms include minimizing braking and propulsive impulse errors from target values, metabolic cost per unit of time or distance^123^ errors from a relative target value, whole-body angular momentum error from a target value, and joint mechanical energy generation or absorption errors from a target value. Two key features of the DO tool are support for model modification functions and user-defined cost function terms. By employing model modification functions alongside GPOPS-II’s static parameters feature, users can change the OpenSim musculoskeletal model, modify the neural control model, or adjust parameter values in an assistive device. Possible model modifications include changing bone geometry (e.g., osteotomy parameters), changing muscle-tendon properties (e.g., strengthening muscles or adjusting optimal muscle fiber lengths), changing neural control properties (e.g., altering synergy vector weights or synergy activation profiles), modifying ground contact model parameters, optimizing muscle attachment locations or muscle wrapping surface parameters, redefining joint parameters, and modifying external device design and/or control parameters. User-defined cost function terms allow users to write a custom MATLAB function, with a specific function signature, that is added as an additional cost function term given the iteration’s states, controls, and static parameters.

## 4. Example Application

### 4.1 Treatment Design Rationale

To demonstrate the capabilities of the NMSM Pipeline, we employed it to investigate a hypothetical treatment to improve walking speed and function for an individual post-stroke. Published studies have hypothesized that time-varying synergy activations originate in the brain while time-invariant synergy vectors reside in the spinal cord.^124,125^ If this hypothesis is true, then one might expect a stroke to affect a patient’s synergy activations but not synergy vectors.^125^ This expectation in turn suggests that an individual post-stroke may have similar synergy vectors between the two legs but lack the ability to recruit one or more of them effectively on the paretic side. Furthermore, published studies have reported that a stroke results in altered neural coordination even on the non-paretic side.^126–128^ Taken together, these observations lead to the hypothesis that a stroke may impair recruitment of one or more synergy activations in the paretic leg, resulting in altered recruitment of one or more synergy activations in the non-paretic leg, and that restoration of these impaired and altered synergy activations could result in improved self-selected walking speed and metabolic cost.^128–130^

To investigate this hypothesis, we used the NMSM Pipeline to design a hypothetical treatment that would modify paretic and non-paretic leg synergy activations for an individual post-stroke with the goal of increasing his self-selected walking speed without increasing his metabolic cost. Previously published experimental walking data collected from an individual post-stroke with a self-selected walking speed of 0.5 m/s were used for the investigation.^41^ Research has shown that both healthy individuals and individuals post-stroke tend to choose a self-selected walking speed to achieve a desired metabolic cost per unit time.^131^ Furthermore, to achieve successful community ambulation (as defined by the walking speed required to cross a typical street before the light changes^132^), individuals should be able to walk at a self-selected speed of 0.8 m/s.^132,133^ Thus, the specific goal of the hypothetical treatment was to increase the subject’s self-selected walking speed to 0.8 m/s while maintaining the same metabolic cost per unit time estimated for the subject’s original self-selected walking speed of 0.5 m/s.

### 4.2 Experimental Data Collection

As described in Meyer et al. (2016),^41^ the experimental data used for the present study were collected from a high-functioning hemiparetic subject with chronic stroke-related walking dysfunction (sex male, age 79 years, LE Fugl-Meyer Motor Assessment 32/34 pts, right-sided hemiparesis, height 1.7 m, mass 80.5 kg). Experimental data collection was approved by the University of Florida Health Science Center Institutional Review Board (IRB-01) and the Malcom Randall VA Medical Center Research and Development Committee, and the subject gave written informed consent prior to participation. Motion capture (Vicon Corp., Oxford, UK), ground reaction (Bertec Corp., Columbus, OH, USA), and electromyography (EMG) data (Motion Lab Systems, Baton Rouge, LA, USA) were collected simultaneously while the subject walked on a split-belt instrumented treadmill with belts tied. The subject walked at his self-selected speed of 0.5 m/s and fastest comfortable speed of 0.8 m/s. To facilitate ground contact model personalization, the subject wore Adidas Samba Classic sneakers, which possess a flat sole with neutral midsole. Motion capture data utilized a full-body marker set with one marker at each toe tip, three markers on each hindfoot, shank, and thigh, three markers on the pelvis, and one marker on each acromion, elbow, and wrist (Reinbolt et al., 2005; Fregly et al., 2007). Ground reaction data from each treadmill force plate included three force and moment components along with the location of the fixed point used for moment calculations. EMG data included a combination of surface and fine-wire electrodes collected from 16 muscles on each leg, including deep muscles adductor longus, iliopsoas, tibialis posterior, extensor digitorum longus, and flexor digitorum longus.

Processing of marker motion, ground reaction, and EMG data followed a previously published protocol.^41^ Marker motion and ground reaction data were filtered at a variable cut-off frequency of 7/tf Hz, where tf is the period of the gait cycle being analyzed, using a fourth-order zero phase lag Butterworth filter.^134^ EMG data were high-pass filtered at 40 Hz,^75^ demeaned, rectified, and low-pass filtered at a variable cut-off frequency 3.5/tf Hz using a fourth-order zero phase lag Butterworth filter. Each processed EMG signal was normalized by the maximum value over all walking cycles and then offset so that the minimum value was zero. For both walking speeds, the single most periodic walking cycle (heel strike to heel strike for the paretic right leg) was selected for analysis, and all data for this one cycle were resampled to 101 time points. After resampling, EMG data were padded with 200 milliseconds (18 time points) before the start of the cycle to account for electromechanical delay.

### 4.3 Model Personalization Process

We utilized all four tools in the Model Personalization toolset to personalize a full-body neuromusculoskeletal model of the subject being studied. The starting point for the model personalization process was a scaled RCNL2023 full-body OpenSim model, where scaling was performed using the OpenSim Model Scaling tool with experimental marker data collected from the subject during a static standing trial. All subsequent NMSM Pipeline tool runs were performed on a Dell PC workstation possessing 48 cores.

First, the JMP tool was used to personalize lower body joint parameters and lower body scaling by minimizing IK marker distance errors. The selected 0.8 m/s gait cycle marker data were used for all JMP tool tasks. To avoid non-anatomical muscle paths around the hip complex, the left and right hip joint parameters were not personalized. The left and right knee and ankle joints were personalized, but the subtalar joint was not personalized because the joint did not move through a sufficient range of motion to personalize the joint’s parameter values accurately. The JMP tool reduced marker distance errors for the thigh, shank, and foot markers by 48.4% for the gait cycle of interest. Additional JMP results are included in Table 1 and in the supplementary material. Convergence was achieved after approximately 90 minutes of CPU time.

**Table 1:** RMS and maximum marker distance errors (mm) for the right and left knee and ankle (including subtalar) joints. Markers included in each joint’s JMP marker set included thigh and shank markers for each knee joint and shank and foot markers for each ankle joint. These markers were used to find the pre- and post-JMP RMS and maximum marker distance errors.

|  | RMS (mm) |  | Max (mm) |  |
| --- | --- | --- | --- | --- |
|  | Pre-JMP | Post-JMP | Pre-JMP | Post-JMP |
| **Right Knee** | 7.5 | 4.2 | 18.5 | 11.8 |
| **Right Ankle** | 6.1 | 3.3 | 16.9 | 9.1 |
| **Left Knee** | 9.5 | 4.1 | 26.3 | 12.8 |
| **Left Ankle** | 6.5 | 3.2 | 16.4 | 10.1 |

Second, the MTP tool was used to personalize the properties of all lower body muscles by minimizing errors in matching inverse dynamic joint moments and changes in initial muscle-tendon parameter values while also predicting missing EMG signals for deep or “small” muscles (e.g., piriformis). Before the MTP tool was run, the OpenSim Inverse Kinematics, Inverse Dynamics, and Muscle Analysis tools were run from the OpenSim GUI using the selected 0.8 m/s walking cycle data. Using the outputs of these GUI tools, an NMSM Pipeline utility script processed EMG data, calculated muscle-tendon velocity from muscle-tendon length data, and automatically organized the trial data into the directory structure required by the MTP tool. Muscle groups were added to the post-JMP OpenSim model to group muscles based on normalized fiber length similarity and activation similarity. Muscle-tendon Length Initialization was used to find initial values for optimal muscle fiber length and tendon slack length. At that point, muscle-tendon model and activation dynamics parameter values were personalized. This process was repeated for each leg separately via separate runs of the MTP tool. After the MTP tool was run for both legs, the left leg was found to match lower body joint moments with an average root mean square (RMS) error of 3.5 Nm and the right leg with an average RMS error of 2.1 Nm. Joint-specific moment matching results are included in Table 2 with additional plots available in the supplementary material. A custom script was used to combine the left and right leg results for use by the NCP tool. Convergence was achieved after approximately 30 minutes of CPU time.

**Table 2:** RMS errors between net joint moments calculated by the OpenSim ID tool and those calculated by the NMSM Pipeline MTP tool following muscle-tendon model personalization.

| Joint<br>Coordinate | Hip<br>Flexion | Hip<br>Adduction | Hip<br>Rotation | Knee<br>Angle | Ankle<br>Angle | Subtalar<br>Angle |
| --- | --- | --- | --- | --- | --- | --- |
| Right (Nm) | 2.9 | 2.7 | 3.1 | 1.7 | 1.0 | 1.5 |
| Left (Nm) | 5.8 | 4.3 | 4.8 | 2.2 | 2.4 | 1.5 |

Third, the NCP tool was used to calculate lower body muscle synergies (i.e., synergy vectors and activations) by minimizing errors in matching inverse dynamic joint moments and muscle activations produced by the MTP tool. Given our treatment design rationale above, we assumed that the subject’s synergy vectors were the same while his synergy activations were different for the two legs. As a result, the NCP tool was used with the bilateral synergy vector symmetry option enabled. Synergy sets were defined as muscle groups denoting the right and left legs. A previous synergy analysis performed on the subject’s processed EMG data indicated that six synergies were sufficient to account for the signal variability even with the added synergy vector symmetry constraint. Muscle Tendon Length Initialization was not performed as part of this NCP tool run because all muscles had MTP-derived muscle activations and thus the activation minimization term was not used. The muscle synergies found by the NCP tool reproduced all lower body joint moments with an average RMSE of 2.2 Nm and all MTP-derived muscle activations with an average RMSE of 0.034. Additional NCP results are included in Table 3 and in the supplementary material. Convergence was achieved after approximately 3 hours of CPU time.

**Table 3:** RMS errors between net joint moments calculated by the OpenSim ID tool and those calculated by the NMSM Pipeline NCP tool following neural control model personalization using muscle synergies.

| Joint Coordinate | Hip Flexion | Hip Adduction | Hip Rotation | Knee Angle | Ankle Angle | Subtalar Angle |
| --- | --- | --- | --- | --- | --- | --- |
| Right (Nm) | 1.2 | 2.2 | 2.9 | 2.7 | 1.0 | 1.5 |
| Left (Nm) | 1.6 | 3.0 | 3.4 | 2.8 | 2.4 | 1.7 |

Fourth, the GCP tool was used to personalize foot-ground contact model parameter values by minimizing errors in matching ground reaction forces, ground reaction moments, and foot kinematics. A contact element grid size of 5 by 11 was selected to match previous successful investigations.^79^ The cost functions for the three tasks performed by the tool were dominated by terms that emphasized matching the vertical GRF, vertical and horizontal GRF, and vertical and horizontal GRF as well as ground reaction moments, respectively, with each task starting from the solution of the previous task to track additional quantities dependent on the previous quantities. To facilitate smooth ground reaction forces during Treatment Optimization, we used the viscous friction coefficient as a design variable and turned off dynamic friction. The viscous friction model penalizes excessive foot velocity with respect to the ground during ground contact, which could lead to excessive horizontal GRFs in predictive simulations. The GCP tool produced a personalized non-uniform distribution of spring stiffness coefficients and matched ground reaction forces, ground reaction moments, foot positions, and foot orientations with RMSEs of 4.6 N, 2.1 Nm, 6.4 mm, and 1.7 degrees, respectively.^135,136^ Additional GCP results are included in Table 4 and in the supplementary material. Convergence was achieved after approximately 4 hours of CPU time.

**Table 4:**
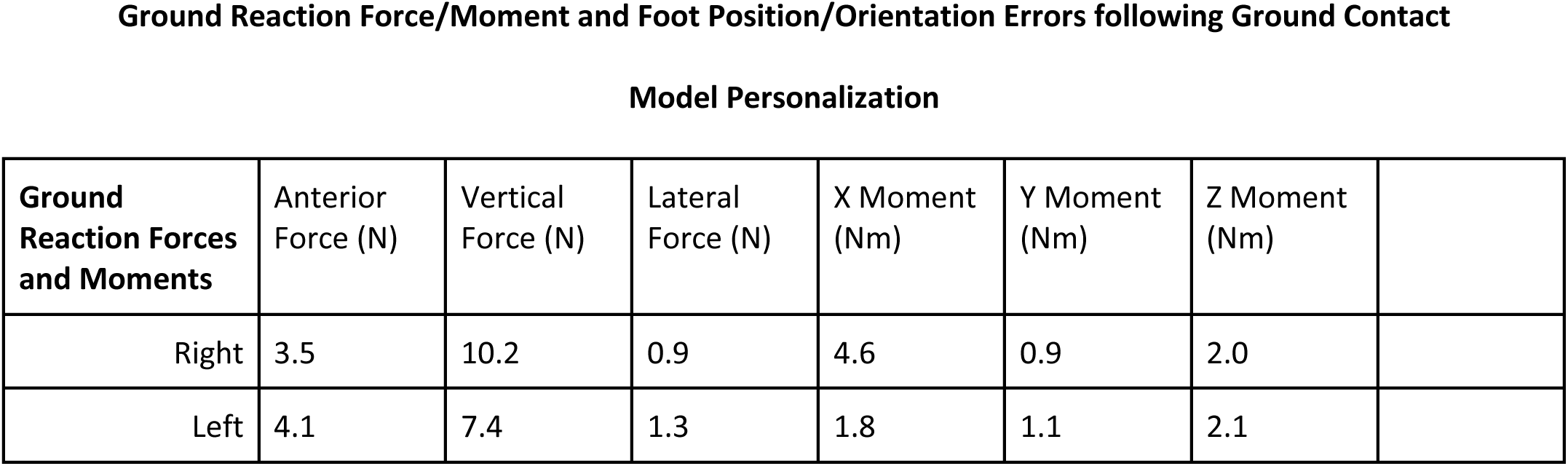
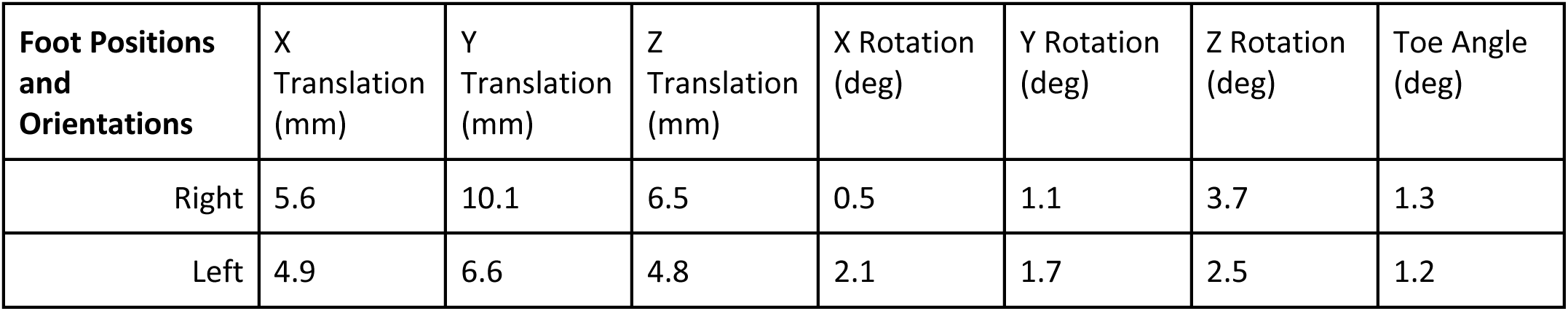
RMS errors between ground reaction forces and moments measured experimentally, and foot positions and orientations measured experimentally and those calculated by the NMSM Pipeline GCP tool following ground contact model personalization.

Model Personalization
| Ground Reaction Forces and Moments | Anterior Force (N) | Vertical Force (N) | Lateral Force (N) | X Moment (Nm) | Y Moment (Nm) | Z Moment (Nm) |
| --- | --- | --- | --- | --- | --- | --- |
| Right | 3.5 | 10.2 | 0.9 | 4.6 | 0.9 | 2.0 |
| Left | 4.1 | 7.4 | 1.3 | 1.8 | 1.1 | 2.1 |

| Foot Positions and Orientations | X Translation (mm) | Y Translation (mm) | Z Translation (mm) | X Rotation (deg) | Y Rotation (deg) | Z Rotation (deg) | Toe Angle (deg) |
| --- | --- | --- | --- | --- | --- | --- | --- |
| Right | 5.6 | 10.1 | 6.5 | 0.5 | 1.1 | 3.7 | 1.3 |
| Left | 4.9 | 6.6 | 4.8 | 2.1 | 1.7 | 2.5 | 1.2 |

### 4.4 Treatment Optimization Process

After personalizing the subject’s neuromusculoskeletal model using all four Model Personalization tools, we ran all three Treatment Optimization tools to predict how underutilized paretic and non-paretic leg synergy activations should be upregulated to achieve a 0.8 m/s self-selected walking speed without increasing metabolic cost per unit time above the subject’s 0.5 m/s level.

First, the TO tool was used to generate a dynamically consistent full body walking motion at 0.8 m/s that closely reproduced the subject’s joint motion, joint moment, ground reaction, and muscle activation data simultaneously using muscle synergy controls for the lower body and joint torque controls for the back and upper body. Since synergy-driven TO requires an initial guess for synergy vectors and activations, the TO tool XML settings file was configured such that the initial guess directly was defined to be the NCP results directory while the tracked quantities directory was defined to be the preprocessed data directory created after running the JMP tool. Other information, including an NMSM Pipeline model file (.osimx), the post-JMP RCNL2023 OpenSim model, and the specified output directory, were also included in the TO tool XML settings file. All lower body joints were controlled by synergy controllers except for the toes coordinate, which was left uncontrolled, and all other joints were controlled by torque controllers. Cost function terms tracked joint positions and velocities found by a post-JMP IK analysis, joint moments found by a post-JMP ID analysis (excluding pelvis residual loads), muscle activations found by MTP, and external loads consisting of processed experimental ground reaction data. Constraint terms included bounded kinetic consistency errors (0.01 Nm) for all controlled joints, root segment residual loads for the pelvis (1 N for forces and 0.1 Nm for moments), and joint position periodicity errors (0.05 rad for joints below the pelvis and 0.1 rad for joints above it). The solution produced by the TO tool matched tracked joint positions with an average RMSE of 2.1 degrees, joint moments with an average RMSE of 5.1 Nm, ground reaction forces with an average RMSE of 25.3 N, ground reaction moments with an average RMSE of 4.0 Nm, and muscle activations with an average RMSE of 0.038. Additional TO results are included in Table 5 and in the supplementary material.

**Table 5:**
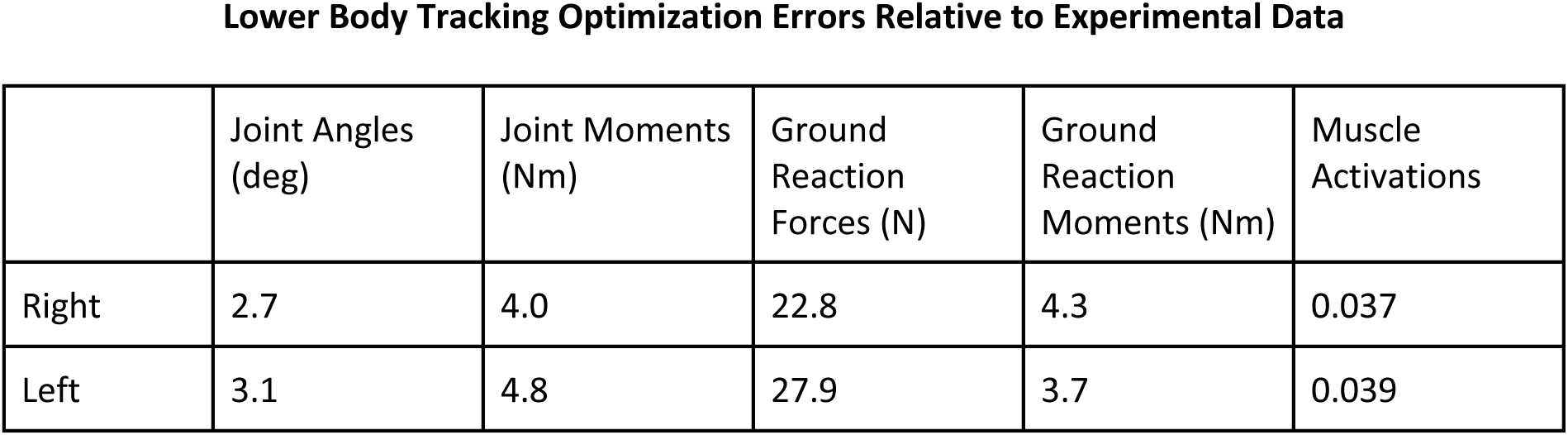
RMS errors between joint angles, joint moments, ground reaction forces, ground reaction moments, and muscle activations for the right and left legs obtained from experimental data and corresponding quantities predicted by the NMSM Pipeline TO tool.

|  | Joint Angles (deg) | Joint Moments (Nm) | Ground Reaction Forces (N) | Ground Reaction Moments (Nm) | Muscle Activations |
| --- | --- | --- | --- | --- | --- |
| Right | 2.7 | 4.0 | 22.8 | 4.3 | 0.037 |
| Left | 3.1 | 4.8 | 27.9 | 3.7 | 0.039 |

Convergence was achieved after 305 IPOPT iterations, which required approximately 770 minutes of CPU time.

Second, the VO tool was used to verify that tracking the muscle synergy controls and upper body joint kinematics found by the TO tool reproduced the joint motions, joint moments, and ground reactions predicted by the TO tool without tracking those quantities explicitly. For the VO run, the results of the previous TO run were used for both the initial guess and tracked quantities data. Cost function terms tracked TO-found synergy controls and TO-found torque-controlled joint positions. Constraint terms included bounded kinetic consistency errors (0.01 Nm) for all controlled joints, root segment residual loads for the pelvis (1 N for forces and 0.1 Nm for moments), and joint position periodicity errors (0.05 rad for joints below the pelvis and 0.1 rad for joints above it). The solution produced by the VO tool matched tracked joint positions with an average RMSE of 0.1 deg, joint moments with an average RMSE of 0.1 Nm, ground reaction forces with an average RMSE of 0.7 N, ground reaction moments with an average RMSE of 0.1 Nm, and muscle activations with an average RMSE of 0.01. Additional MTP results are included in Table 6 and in the supplementary material.

**Table 6:** RMS errors between joint angles, joint moments, ground reaction forces, ground reaction moments, and muscle activations for the right and left legs obtained from the NMSM Pipeline TO tool and corresponding quantities predicted by the NMSM Pipeline VO tool.

|  | Joint Angles (deg) | Joint Moments (Nm) | Ground Reaction Forces (N) | Ground Reaction Moments (Nm) | Muscle Activations |
| --- | --- | --- | --- | --- | --- |
| Right | 0.09 | 0.06 | 0.44 | 0.06 | 0.01 |
| Left | 0.08 | 0.09 | 0.97 | 0.09 | 0.01 |

Convergence was achieved after 101 IPOPT iterations, which required approximately 170 minutes of CPU time. Since the VO results closely matched the TO results, we were confident that the controller and joint position tracking cost function terms and associated constraint terms selected for the verification optimization could be used as the basis for the design optimization.

Finally, the DO tool was used to explore whether upregulating underutilized synergy activations in the paretic leg and downregulating overutilized synergy activations in the non-paretic legs could produce a faster self-selected walking speed of 0.8 m/s while maintaining the same metabolic cost per unit time calculated for the subject’s original self-selected walking speed of 0.5 m/s. The hypothesis in this problem formulation is that large differences in synergy activation amplitudes between the two legs are indicative of impaired paretic leg synergy activations that are smaller than desired and compensatory non-paretic leg synergy activations that are larger than desired. When we evaluated the synergy activations found by VO for the two legs, we identified four pairs that varied significantly in amplitude between the left and right sides. Based on this observation, we formulated a DO problem that sought to bring these four synergy activations with disparate amplitudes in the two legs closer to each other. Each pair of asymmetric synergy activations contained a “weak” and “strong” side. First, scaled-up “weak” synergy activations were tracked for each pair to increase their amplitudes. Second, a custom user-defined cost function was written to reward synergy amplitude similarities between the “weak” and “strong” synergy activations. A user-defined static parameter was added for each of the four synergy pairs representing the amplitude scale factor to be applied to the “strong“ synergy activation using a user-defined model modification function. These two terms increased weak synergy activations and encouraged over-active synergy activations to reduce to more symmetric magnitudes while not overconstraining the control space. In addition, a cost function term was added to find a solution with a metabolic cost per unit time that matched the value calculated for the subject’s original 0.5 m/s self-selected walking speed. This value was 90% of the original value calculated for the subject’s 0.8 m/s self-selected walking speed, making a 10% reduction to the target value. Cost function terms also tracked VO-found synergy controls for the two symmetric control pairs and VO-found torque controls.

Constraint terms included bounded kinetic consistency errors (0.01 Nm) for all controlled joints, root segment residual loads for the pelvis (1 N for forces and 0.1 Nm for moments), and joint position periodicity errors (0.05 rad for joints below the pelvis and 0.1 rad for joints above it). The VO results were used in both the initial guess and tracked quantities data directories, and the final time was fixed at the value determined experimentally for the selected 0.8 m/s walking trial.

The DO predictive simulation produced modified kinematics, synergy activations, and ground reactions that reduced the subject’s metabolic cost per unit time by 9.9% for the selected gait cycle and achieved closely matching maximum amplitudes for the four modified synergy activations. Final results are plotted in Figure 11. Convergence was achieved after 489 IPOPT iterations, which required approximately 730 minutes of CPU time. These DO results suggest that upregulating weak paretic leg synergy activations while downregulating paired non-paretic leg synergy activations could potentially allow the modeled subject to increase his self-selected walking speed by 60% without increasing his metabolic cost per unit time.

**Figure 11:**
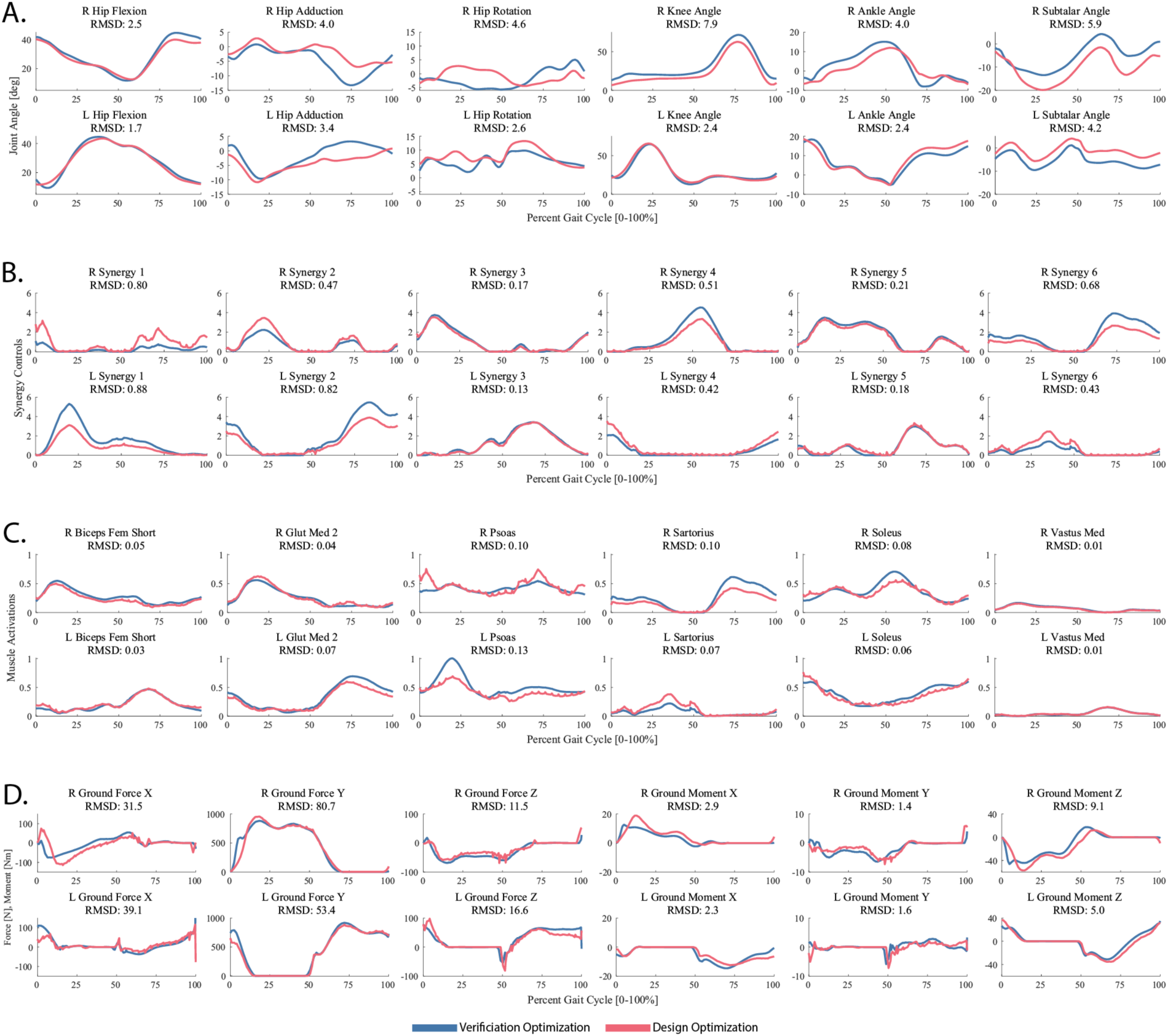
Plots showing the final results of the DO tool. Subfigures A) kinematics, B) synergy activations, C) muscle activations, and D) ground reaction forces and moments after VO (blue) and DO (red), respectively.

## Discussion

The primary objective of this work was to develop a novel set of easy-to-use computational tools that facilitate the creation of personalized neuromusculoskeletal models and the subsequent design of personalized clinical treatments by researchers and clinicians with little to no programming experience. These tools dramatically reduce the barrier to entry for researchers and clinicians working together who would like to explore using personalized computational neuromusculoskeletal models to design personalized treatment approaches. As shown in the example application, the NMSM Pipeline has the potential to change the way treatments are investigated and applied throughout the neuromusculoskeletal injury space.

Going forward, two key questions need to be addressed if the NMSM Pipeline is to be used to design actual clinical treatments for actual patients. The first question is, how should the patient’s personalized model and neural control strategy be changed in response to the simulated treatment? The second question is, how can one validate a patient’s predicted post-treatment movement function? The answers to these two questions will depend on the specific clinical application and which aspects of the patient’s neuromusculoskeletal system need to be modeled and personalized to investigate that application. The answers will also involve developing an understanding of how the planned intervention will affect a patient’s model parameter values and neural control strategy. Collection of post-intervention movement data will be required so that the entire Model Personalization and Treatment Optimization process can be evaluated, refined, and ultimately validated for one clinical application at a time.

To conceptualize how these two important questions could be addressed, one can consider two NSM Pipeline clinical applications being explored by the authors - one involving knee osteoarthritis rehabilitation design, and one involving pelvic cancer surgical planning. Past computational research involving a subject with bilateral medial compartment knee osteoarthritis investigated whether personalized gait modifications could be designed to reduce the subject’s peak adduction moment in both knees to an extent comparable to invasive high tibial osteotomy surgery [write reference in (author, year) form]. For this treatment optimization problem, no muscles or neural control needed to be modeled, which simplified the treatment design process. The computational treatment design process involved: a) creating a personalized full-body dynamic skeletal model of the subject using pre-rehabilitation gait data, b) selecting a post-treatment control approach to be used when predicting post-treatment walking function (in this case, minimize changes in lower body joint moments relative to pre-rehabilitation gait data), c) performing a treatment optimization process with the personalized model and selected control approach to identify gait modifications that would offload the medial compartment of both knees, d) training the subject to perform and consolidate the modified walking motion, and finally e) re-testing the subject in a gait lab to evaluate how well the predicted gait modifications could reduce both peak adduction moment peaks. After informal training, the subject was able to perform the predicted gait modifications and achieve bilateral reductions in the peak knee adduction moment that were comparable to the treatment optimization predictions. A similar computational treatment design process is currently being repeated using the NMSM Pipeline except with greater flexibility in the range of potential gait modifications that can be predicted.

Ongoing computational research involving subjects with pelvic cancer is investigating whether surgical decisions can be planned to achieve a post-surgery gait motion and self-selected walking speed that is as close to normal as possible for each patient^68,137^. For this treatment optimization problem, both muscles and neural control need to be modeled, which complicates the treatment design process.

The computational treatment design process being explored involves: a) creating a personalized full-body dynamic neuromusculoskeletal model of the patient using pre-surgery gait, EMG, and imaging data (to personalize pelvis bony geometry), b) selecting a post-surgery control approach to be used when predicting post-surgery walking function (in this case, minimize changes in muscle synergy quantities relative to pre-surgery EMG data)^137^, c) performing a treatment optimization process with the personalized model and selected control approach to identify surgical decisions that achieve clear margins while also maximizing recovery of normal walking function^68^, d) having the orthopedic oncologist perform the patient’s surgery using the optimized surgical plan, and finally e) re-testing the patient in a gait lab following plateau in recovery to evaluate how well the predicted surgical plan allowed the patient to achieve the predicted post-surgery gait motion and speed. Since trial and error treatment is not possible with orthopedic surgeries, extensive testing, evaluation, and refinement of this treatment optimization process must be performed before actual surgical plans can be designed for actual pelvic cancer patients. To this end, the authors are collecting extensive pre- and post-surgery gait, EMG, and imaging data sets from this patient population, along with the implemented surgical decisions. Only after the NMSM Pipeline can predict post-surgery walking function reliably for multiple patients when post-surgery walking function and the implemented surgical decisions are known can the NMSM Pipeline be considered for actual use in this clinical application. A similar process would need to be followed for any other clinical problem to which the NMSM Pipeline would be applied.

Although the decision to use MATLAB for the NMSM Pipeline facilitates development and ease of access for users, the decision does create several limitations. C++ and other similar compiled programming languages generally run significantly faster than interpreted programming languages like MATLAB. However, they also introduce challenges since few engineering researchers are strong C++ programmers, and system-specific build procedures can often be complicated.^138,139^ Due to these drawbacks, most biomechanics software written in compiled languages can be modified only by reporting a bug or requesting a feature and waiting for experienced software developers to release a new version of the software. The NMSM Pipeline actively encourages modifications, bug fixes, and improvements through open-source easy-to-modify MATLAB toolsets. Other interpreted programming languages like Python can struggle with package management, virtual environments, and/or dependency version mismatches that increase barriers to use. MATLAB’s native debugging tools, English-readable error messages, and robust, stable core functions allow for development with confidence, making the NMSM Pipeline robust to issues common to other interpreted languages.

Our hope is that the NMSM Pipeline’s open-source, easy-to-use framework will encourage investigations by the research community into “best practices” for each of the included tools. With the availability of seven new tools, each with many optional features, the research community is encouraged to explore novel approaches to achieving the best results from each tool. Investigations into changes in task order, data processing/organization, cost function error centers and maximum allowable errors, among others, will inform the entire research community on the best ways to leverage these unique tools for successful future research and clinical outcomes.

We are already planning future enhancements to improve the versatility and applicability of the NMSM Pipeline to a broader range of clinical and research problems. Potential enhancements would allow greater flexibility and customization during use of the Model Personalization toolset. Support for compliant tendon muscle models and muscle models that differentiate between fast and slow twitch fibers in the MTP tool could facilitate muscle-tendon model personalization and simulation for “fast” activities like running or jumping.^61,140^ Support for time-delayed sensory feedback models in the NCP tool could enable investigation of how spasticity, hyperreflexia, and hypertonia interact with incoordination and weakness to produce functional impairment in individuals with an upper motor neuron pathology. Support for multi-segment foot models in the GCP tool could facilitate higher fidelity simulations of foot biomechanics and better reproduction of ground reaction moments. Other potential enhancements could facilitate novel use of the Treatment Optimization toolset. User-defined controls, time-delayed sensory feedback controls, and individual muscle controls could facilitate computational design of personalized functional electrical stimulation protocols^67^ and prediction of how downregulating overactive reflexes could improve an individual’s movement function. Support for user-defined loads as a function of model states and controls and making it easy to add new states and controls to a DO problem could facilitate computational design of personalized passive and/or active exoskeletons, which would be added to a subject’s personalized model after TO and VO had been performed. An enhancement supporting multi-phase optimal control problems, where each phase simulates a different movement cycle with control information shared across phases, could facilitate simulating neural feedback mechanisms, neural adaptation, and cycle-to-cycle motion and control variability. Enhancements supporting the use of closed-chain kinematic models would facilitate simulation of shoulder models with improved kinematic fidelity for surgical, exoskeleton, and exercise equipment applications. Support for connecting NMSM Pipeline controls to any type of OpenSim actuator could allow exploration of alternative optimal control problem formulations that significantly improve IPOPT convergence rate, especially for TO. Finally, for both toolsets, development of easy-to-use MATLAB utility functions that automatically process experimental movement data and put it into a format that can be used directly by NMSM Pipeline tools, without requiring any manual data manipulation, would make the toolsets accessible to a wider user base.

With all of these potential future enhancements, several limitations and challenges in future NMSM Pipeline development exist. First, the quality of the personalization results found by the Model Personalization tools is limited by the quality and quantity of collected experimental data. At this time, it can be challenging to collect all of the experimental data needed to exercise all seven NMSM Pipeline tools with a synergy-driven neuromusculoskeletal model. Leveraging markerless motion capture or inertial measurement units could potentially increase the ability of researchers to collect the movement data required to perform a high-quality pipeline run-through. For Model Personalization, because gradient-based optimization techniques are used, solutions may become entrapped in a local minimum, though the final solution will still be better than the initial guess. The use of muscle activations rather than excitations for the NCP tool and the Treatment Optimization toolset is another limitation, though back calculation of muscle excitations from muscle activations could potentially be performed given estimates of activation dynamics parameter values. Using the OpenSim framework through the OpenSim MATLAB API prevents the NMSM Pipeline from using automatic differentiation to speed up optimizations performed during the Model Personalization or Treatment Optimization process.

Computations leveraging the OpenSim MATLAB API are inefficient, and speed limitations inherent to this framework cannot be overcome.

Several research teams have released novel software tools for the musculoskeletal modeling research community^51–58^, and these tools could be used synergistically with the NMSM Pipeline. As shown in Table 7, though none of these tools possess the broad range of model personalization and treatment optimization capabilities available in the NMSM Pipeline, all of these tools facilitate some aspect of creating scaled musculoskeletal models that incorporate personalized bone geometry and muscle attachment locations. Consequently, these tools can provide the scaled musculoskeletal model required as a starting point for the NMSM Pipeline Model Personalization process. For example, NMSBuilder, MAP Client, Torsion Tool, Bone Deformation Tool, or OpenSim Creator could be used to create personalized bone models from imaging data or physical measurements^52–55,57,141,142^. SimCP, NMSBuilder, MAP Client, or OpenSim Creator could then be used to personalize muscle attachment locations^51–53,57^. Next, SimCP, NMSBuilder, MAP Client, AddBiomechanics, or Scale Tool could be used to create a scaled OpenSim musculoskeletal model possessing the personalized bone geometry and muscle attachments^51–53,56^. In addition, AddBiomechanics could potentially be used to perform limited initial personalization of joint functional axes^56^, while SimCP could be used to perform initial personalization of optimal muscle fiber lengths, tendon slack lengths, and synergy control properties^51^.

**Table 7:**

Comparison of Functionality Available in the NMSM Pipeline and Other Musculoskeletal Modeling Software Tools
| Software Tool | Personalize Bone Geometry | Personalize Muscle Attachments | Scale Generic Models | JMP | MTP | NCP | GCP | TO | VO | DO |
| --- | --- | --- | --- | --- | --- | --- | --- | --- | --- | --- |
| NMSM Pipeline | No (1) | No | No | Yes | Yes | Yes | Yes | Yes | Yes | Yes |
| SimCP | No (1) | Yes | Yes | No | Yes | Yes | No | No | No | No |
| NMS Builder | Yes | Yes | Yes | No | No | No | No | No | No | No |
| MAP Client | Yes | Yes | Yes | No | No | No | No | No | No | No |
| Torsion Tool | Yes | No | No | No | No | No | No | No | No | No |
| Bone Deformation Tool | Yes | No | No | No | No | No | No | No | No | No |
| AddBiomechanics | No | No | Yes | Yes (2) | No | No | No | No | No | No |
| OpenSim Creator | Yes | Yes | No | No | No | No | No | No | No | No |
| Automatic Scale Tool | No | No | Yes | No | No | No | No | No | No | No |

(1) Personalized bone models produced using image processing and surface fitting software (e.g., Mimics, Materialize, Leuven, Belgium; Geomagic Wrap, Oqton, Ghent, Belgium) can be added.
(2) Some, but not all, of the features of JMP

As shown in Table 7 above, a current limitation of the NMSM Pipeline is the lack of functionality to personalize a patient’s bone morphology to clinical imaging data. The NMSM Pipeline was specifically designed to avoid the use of imaging data and require only experimental data typically collected in a human movement lab. However, personalized bone geometry generated using existing methods and software tools^143,53,137^ can easily be incorporated into scaled generic OpenSim models with personalized joint functional axes^137^ to produce more accurate muscle tendon lengths, moment arms, and joint and bone loading scenarios.

Treatment Optimization also does not currently include a muscle fatigue model. To date, few muscle fatigue models for neuromusculoskeletal modeling applications have been published in the literature^144,145^. Thus, selecting an appropriate existing muscle fatigue model would be challenging.

Availability of a muscle fatigue model could improve the utility of predictive simulations for athletic, other high-intensity, or long duration movements, making this research area an important one for future investigation.

The NMSM Pipeline’s Treatment Optimization toolset has key differences in design and implementation from the existing OpenSim Moco tool. While Moco is generally the faster software due to its C++-based implementation, Treatment Optimization leverages the flexibility of MATLAB and GPOPS-II to facilitate novel features. Current unique features include user-defined functions and the use of .osimx personalized model components, while future unique features could include support for multi- phase optimal control problems, as discussed above. Treatment Optimization was designed to be accessible to both novice and advanced users through intuitive XML settings files, whereas Moco users typically set up problems through MATLAB or Python scripts. Even so, Moco’s implementation facilitates potential use of C++ automatic differentiation libraries to improve computational speed for gradient calculations, resulting in more efficient optimizations. Lastly, Moco is free while both MATLAB and GPOPS-II are commercial products with significant academic discounts. By incorporating NMSM Pipeline model components into native OpenSim in the future, we plan to give users the option of using their NMSM Pipeline personalized models in either our Treatment Optimization toolset or OpenSim Moco, allowing the research community to explore the tradeoffs between these two approaches to developing predictive simulations.

In conclusion, the NMSM Pipeline represents a significant step forward in the ability to personalize neuromusculoskeletal computer models to patient movement data and then optimize parameters of potential treatments using the personalized models. The software assembles over two decades of biomechanics research into a well-designed and accessible framework that makes previously “high end” modeling, simulation, and optimization techniques accessible to the entire research community. Our hope is that by empowering researchers and clinicians working together to explore clinically relevant questions and to design novel personalized interventions, the NMSM Pipeline will bring neuromusculoskeletal computer modeling not only to the doorstep of clinical utility but also across it.

## Declarations

### Ethics Approval and Consent to Participate

The de-identified experimental walking data used in this study were collected and disseminated as part of a previously published study.^41^

### Consent for Publication

Not applicable.

### Availability of Data and Materials

The NMSM Pipeline software, along with all data, models, and settings files described in this article, are available from the “Neuromusculoskeletal Modeling (NMSM) Pipeline” project on SimTK.org (https://simtk.org/projects/nmsm). Users are welcome to share feedback and request new functionality through this project’s forum. Documentation including implementation details, examples, and tutorials is available at https://nmsm.rice.edu.

Further details on the NMSM Pipeline software are provided below: Project name: Neuromusculoskeletal Modeling Pipeline Project home page: https://nmsm.rice.edu

Archived version:

Operating systems: Windows, MacOS Programming language: MATLAB

Other requirements: Version 2023a or higher License: Apache 2.0

The software has no restrictions for use by non-academics.

### Authors’ Information

Claire V. Hammond -

Spencer T. Williams -

Marleny M. Vega -

Di Ao -

Geng Li -

Robert M. Salati -

Kayla M. Pariser -

Mohammad S. Shourijeh -

Ayman W. Habib -

Carolynn Patten -

Benjamin J. Fregly -

## Acknowledgments

The authors would like to thank Dr. Marleny Vega for the foundational work she did organizing approximately 10 years of MATLAB code written by previous Ph.D. students so that it could be used as the starting point for developing the NMSM Pipeline software. Without that extensive effort, the NMSM Pipeline software would not have been possible.

## Abbreviations

NMSM: Neuromusculoskeletal Modeling
JMP: Joint Model Personalization
MTP: Muscle-tendon Model Personalization
NCP: Neural Control Model Personalization
GCP: Ground Contact Model Personalization
TO: Tracking Optimization
VO: Verification Optimization
DO: Design Optimization
SynX: Synergy Extrapolation
IK: Inverse Kinematics
ID: Inverse Dynamics
EMG: Electromyography
DOF: Degrees of Freedom
RMS: Root Mean Squared
ADL: Activity of Daily Living
TKR: Total Knee Replacement
CP: Cerebral Palsy
GUI: Graphical User Interface
XML: Extensible Markup Language

## References

(1) Ma, V. Y.; Chan, L.; Carruthers, K. J. The Incidence, Prevalence, Costs and Impact on Disability of Common Conditions Requiring Rehabilitation in the US: Stroke, Spinal Cord Injury, Traumatic Brain Injury, Multiple Sclerosis, Osteoarthritis, Rheumatoid Arthritis, Limb Loss, and Back Pain. Arch. Phys. Med. Rehabil. 2014, 95 (5), 986–995.e1. 10.1016/j.apmr.2013.10.032.

(2) Ovbiagele, B.; Goldstein, L. B.; Higashida, R. T.; Howard, V. J.; Johnston, S. C.; Khavjou, O. A.; Lackland, D. T.; Lichtman, J. H.; Mohl, S.; Sacco, R. L.; Saver, J. L.; Trogdon, J. G.; American Heart Association Advocacy Coordinating Committee and Stroke Council. Forecasting the Future of Stroke in the United States: A Policy Statement from the American Heart Association and American Stroke Association. Stroke 2013, 44 (8), 2361–2375. 10.1161/STR.0b013e31829734f2.

(3) Theis, K. A.; Roblin, D. W.; Helmick, C. G.; Luo, R. Prevalence and Causes of Work Disability among Working-Age U.S. Adults, 2011-2013, NHIS. Disabil. Health J. 2018, *11* (1), 108–115. 10.1016/j.dhjo.2017.04.010.

(4) Praemer, A.; Furner, S.; Rice, D. P. *Musculoskeletal Conditions in the United States*; American Academy of Orthopaedic Surgeons: Rosemont, IL, 1999.

(5) Clark, M. S.; Smith, D. S. Factors Contributing to Patient Satisfaction with Rehabilitation Following Stroke. Int. J. Rehabil. Res. 1998, 21 (2), 143.

(6) Brandstater, M. E.; de Bruin, H.; Gowland, C.; Clark, B. M. Hemiplegic Gait: Analysis of Temporal Variables. Arch. Phys. Med. Rehabil. 1983, 64 (12), 583–587.

(7) Chen, G.; Patten, C.; Kothari, D. H.; Zajac, F. E. Gait Deviations Associated with Post-Stroke Hemiparesis: Improvement during Treadmill Walking Using Weight Support, Speed, Support Stiffness, and Handrail Hold. Gait Posture 2005, 22 (1), 57–62. 10.1016/j.gaitpost.2004.06.008.

(8) Olney, S. J.; Monga, T. N.; Costigan, P. A. Mechanical Energy of Walking of Stroke Patients. Arch. Phys. Med. Rehabil. 1986, 67 (2), 92–98. 10.1016/0003-9993(86)90109-7.

(9) Rejnö, Å.; Nasic, S.; Bjälkefur, K.; Bertholds, E.; Jood, K. Changes in Functional Outcome over Five Years after Stroke. Brain Behav. 2019, 9 (6), e01300. 10.1002/brb3.1300.

(10) Barbour, K. E. Vital Signs: Prevalence of Doctor-Diagnosed Arthritis and Arthritis-Attributable Activity Limitation — United States, 2013–2015. MMWR Morb. Mortal. Wkly. Rep. 2017, 66. 10.15585/mmwr.mm6609e1.

(11) Kahlenberg, C. A.; Nwachukwu, B. U.; McLawhorn, A. S.; Cross, M. B.; Cornell, C. N.; Padgett, D. E. Patient Satisfaction after Total Knee Replacement: A Systematic Review. HSS Journal® 2018, 14 (2), 192–201. 10.1007/s11420-018-9614-8.

(12) Muertizha, M.; Cai, X.; Ji, B.; Aimaiti, A.; Cao, L. Factors Contributing to 1-Year Dissatisfaction after Total Knee Arthroplasty: A Nomogram Prediction Model. *J*. Orthop. Surg. 2022, 17 (1), 367. 10.1186/s13018-022-03205-2.

(13) Bourne, R. B.; Chesworth, B. M.; Davis, A. M.; Mahomed, N. N.; Charron, K. D. J. Patient Satisfaction after Total Knee Arthroplasty: Who Is Satisfied and Who Is Not? Clin. Orthop. 2009, 468 (1), 57. 10.1007/s11999-009-1119-9.

(14) Anakwe, R. E.; Jenkins, P. J.; Moran, M. Predicting Dissatisfaction After Total Hip Arthroplasty: A Study of 850 Patients. J. Arthroplasty 2011, 26 (2), 209–213. 10.1016/j.arth.2010.03.013.

(15) Tilbury, C.; Haanstra, T. M.; Leichtenberg, C. S.; Verdegaal, S. H. M.; Ostelo, R. W.; Vet, H. C. W. de; Nelissen, R. G. H. H.; Vlieland, T. P. M. V. Unfulfilled Expectations After Total Hip and Knee Arthroplasty Surgery: There Is a Need for Better Preoperative Patient Information and Education. J. Arthroplasty 2016, 31 (10), 2139–2145. 10.1016/j.arth.2016.02.061.

(16) Kim, M. S.; Koh, I. J.; Kim, C. K.; Choi, K. Y.; Yang, J. S.; In, Y. Patient Expectations and Satisfaction After Medial Opening Wedge High Tibial Osteotomy. J. Arthroplasty 2020, 35 (12), 3467–3473. 10.1016/j.arth.2020.06.076.

(17) Stephan-Carlier, A.; Facione, J.; Speyer, E.; Rumilly, E.; Paysant, J. Quality of Life and Satisfaction after Multilevel Surgery in Cerebral Palsy: Confronting the Experience of Children and Their Parents. Ann. Phys. Rehabil. Med. 2014, 57 (9–10), 640–652. 10.1016/j.rehab.2014.09.012.

(18) Blumetti, F. C.; Wu, J. C. N.; Bau, K. V.; Martin, B.; Hobson, S. A.; Axt, M. W.; Selber, P. Orthopedic Surgery and Mobility Goals for Children with Cerebral Palsy GMFCS Level IV: What Are We Setting out to Achieve? J. Child. Orthop. 2012, 6 (6), 485–490. 10.1007/s11832-012-0454-7.

(19) Schwartz, M. H.; Ries, A. J.; Georgiadis, A. G.; Kainz, H. Demonstrating the Utility of Instrumented Gait Analysis in the Treatment of Children with Cerebral Palsy. PLOS ONE 2024, 19 (4), e0301230. 10.1371/journal.pone.0301230.

(20) DadeMatthews, O. O.; Roper, J. A.; Vazquez, A.; Shannon, D. M.; Sefton, J. M. Prosthetic Device and Service Satisfaction, Quality of Life, and Functional Performance in Lower Limb Prosthesis Clients. Prosthet. Orthot. Int. 2024, 48 (4), 422–430. 10.1097/PXR.0000000000000285.

(21) Datta, D.; Selvarajah, K.; Davey, N. Functional Outcome of Patients with Proximal Upper Limb Deficiency--Acquired and Congenital. Clin. Rehabil. 2004, 18 (2), 172–177. 10.1191/0269215504cr716oa.

(22) The designer changing the way aircraft are built. https://www.bbc.com/future/article/20181129-the-ai-transforming-the-way-aircraft-are-built (accessed 2024-09-13).

(23) SpaceX Is Using These Simulations to Design the Rocket That’ll Take Us to Mars. https://www.vice.com/en/article/spacex-is-using-these-simulations-to-design-the-rocket-thatll-take-us-to-mars/ (accessed 2024-09-13).

(24) A brief history of computing in Formula 1. https://www.mclaren.com/racing/team/a-brief-history-of-computing-in-f1-1052199/ (accessed 2024-09-13).

(25) The America’s Cup: nerves, skill, and computer design | PCWorld. https://www.pcworld.com/article/447831/the-americas-cup-nerves-skill-and-a-lot-of-computers.html (accessed 2024-09-13).

(26) Planning and Design of Engineering Systems | Graeme Dandy, David Walke. https://www.taylorfrancis.com/books/mono/10.1201/9781351228121/planning-design-engineering-systems-graeme-dandy-trevor-daniell-robert-warner-bernadette-foley-graeme-dandy-david-walker-trevor-daniell-robert-warner (accessed 2024-09-13).

(27) López Gualdrón, C.-I.; Bravo Ibarra, E.-R.; Murillo Bohórquez, A.-P.; Garnica Bohórquez, I. Present and Future for Technologies to Develop Patient-Specific Medical Devices: A Systematic Review Approach. Med. Devices Auckl. NZ 2019, 12, 253–273. 10.2147/MDER.S215947.

(28) U, V.; Mehrotra, D.; Howlader, D.; Singh, P. K.; Gupta, S. Patient Specific Three-Dimensional Implant for Reconstruction of Complex Mandibular Defect. J. Craniofac. Surg. 2019, 30 (4), e308– e311. 10.1097/SCS.0000000000005228.

(29) Sutradhar, A.; Park, J.; Carrau, D.; Miller, M. J. Experimental Validation of 3D Printed Patient-Specific Implants Using Digital Image Correlation and Finite Element Analysis. Comput. Biol. Med. 2014, 52, 8–17. 10.1016/j.compbiomed.2014.06.002.

(30) Pinheiro, M.; Alves, J. L. The Feasibility of a Custom-Made Endoprosthesis in Mandibular Reconstruction: Implant Design and Finite Element Analysis. J. Cranio-Maxillofac. Surg. 2015, 43 (10), 2116–2128. 10.1016/j.jcms.2015.10.004.

(31) Transforming the Diagnosis and Management of Coronary Artery Disease. https://www.heartflow.com/. https://www.heartflow.com/ (accessed 2024-09-13).

(32) Hlatky, M. A.; De Bruyne, B.; Pontone, G.; Patel, M. R.; Norgaard, B. L.; Byrne, R. A.; Curzen, N.; Purcell, I.; Gutberlet, M.; Rioufol, G.; Hink, U.; Schuchlenz, H. W.; Feuchtner, G.; Gilard, M.; Andreini, D.; Jensen, J. M.; Hadamitzky, M.; Wilk, A.; Wang, F.; Rogers, C.; Douglas, P. S.; PLATFORM Investigators. Quality-of-Life and Economic Outcomes of Assessing Fractional Flow Reserve With Computed Tomography Angiography: PLATFORM. J. Am. Coll. Cardiol. 2015, 66 (21), 2315–2323. 10.1016/j.jacc.2015.09.051.

(33) Douglas, P. S.; Pontone, G.; Hlatky, M. A.; Patel, M. R.; Norgaard, B. L.; Byrne, R. A.; Curzen, N.; Purcell, I.; Gutberlet, M.; Rioufol, G.; Hink, U.; Schuchlenz, H. W.; Feuchtner, G.; Gilard, M.; Andreini, D.; Jensen, J. M.; Hadamitzky, M.; Chiswell, K.; Cyr, D.; Wilk, A.; Wang, F.; Rogers, C.; De Bruyne, B. Clinical Outcomes of Fractional Flow Reserve by Computed Tomographic Angiography-Guided Diagnostic Strategies vs. Usual Care in Patients with Suspected Coronary Artery Disease: The Prospective Longitudinal Trial of FFRCT: Outcome and Resource Impacts Study. Eur. Heart J. 2015, 36 (47), 3359–3367. 10.1093/eurheartj/ehv444.

(34) Fregly, B. J. A Conceptual Blueprint for Making Neuromusculoskeletal Models Clinically Useful. Appl. Sci. 2021, 11 (5), 2037. 10.3390/app11052037.

(35) Umberger, B. R.; Miller, R. H. Optimal Control Modeling of Human Movement. In Handbook of Human Motion; Müller, B., Wolf, S. I., Brueggemann, G.-P., Deng, Z., McIntosh, A., Miller, F., Selbie, W. S., Eds.; Springer International Publishing: Cham, 2017; pp 1–22. 10.1007/978-3-319-30808-1_177-1.

(36) Lee, L.-F.; Umberger, B. R. Generating Optimal Control Simulations of Musculoskeletal Movement Using OpenSim and MATLAB. PeerJ 2016, 4, e1638. 10.7717/peerj.1638.

(37) Porsa, S.; Lin, Y.-C.; Pandy, M. G. Direct Methods for Predicting Movement Biomechanics Based Upon Optimal Control Theory with Implementation in OpenSim. Ann. Biomed. Eng. 2016, 44 (8), 2542–2557. 10.1007/s10439-015-1538-6.

(38) Pandy, M. G.; Zajac, F. E.; Sim, E.; Levine, W. S. An Optimal Control Model for Maximum-Height Human Jumping. J. Biomech. 1990, 23 (12), 1185–1198. 10.1016/0021-9290(90)90376-E.

(39) van den Bogert, A. J.; Hupperets, M.; Schlarb, H.; Krabbe, B. Predictive Musculoskeletal Simulation Using Optimal Control: Effects of Added Limb Mass on Energy Cost and Kinematics of Walking and Running. Proc. Inst. Mech. Eng. Part P J. Sports Eng. Technol. 2012, 226 (2), 123–133. 10.1177/1754337112440644.

(40) Rohani, F.; Richter, H.; Bogert, A. J. van den. Optimal Design and Control of an Electromechanical Transfemoral Prosthesis with Energy Regeneration. PLOS ONE 2017, 12 (11), e0188266. 10.1371/journal.pone.0188266.

(41) Meyer, A. J.; Eskinazi, I.; Jackson, J. N.; Rao, A. V.; Patten, C.; Fregly, B. J. Muscle Synergies Facilitate Computational Prediction of Subject-Specific Walking Motions. Front. Bioeng. Biotechnol. 2016, 4. 10.3389/fbioe.2016.00077.

(42) Dorn, T. W.; Wang, J. M.; Hicks, J. L.; Delp, S. L. Predictive Simulation Generates Human Adaptations during Loaded and Inclined Walking. PLOS ONE 2015, 10 (4), e0121407. 10.1371/journal.pone.0121407.

(43) Ong, C. F.; Geijtenbeek, T.; Hicks, J. L.; Delp, S. L. Predicting Gait Adaptations Due to Ankle Plantarflexor Muscle Weakness and Contracture Using Physics-Based Musculoskeletal Simulations. PLOS Comput. Biol. 2019, 15 (10), e1006993. 10.1371/journal.pcbi.1006993.

(44) Miller, R. H.; Esterson, A. Y.; Shim, J. K. Joint Contact Forces When Minimizing the External Knee Adduction Moment by Gait Modification: A Computer Simulation Study. The Knee 2015, 22 (6), 481–489. 10.1016/j.knee.2015.06.014.

(45) Miller, R. H. A Comparison of Muscle Energy Models for Simulating Human Walking in Three Dimensions. J. Biomech. 2014, 47 (6), 1373–1381. 10.1016/j.jbiomech.2014.01.049.

(46) McGowan, C. P.; Neptune, R. R.; Clark, D. J.; Kautz, S. A. Modular Control of Human Walking: Adaptations to Altered Mechanical Demands. J. Biomech. 2010, 43 (3), 412–419. 10.1016/j.jbiomech.2009.10.009.

(47) Allen, J. L.; Neptune, R. R. Three-Dimensional Modular Control of Human Walking. J. Biomech. 2012, 45 (12), 2157–2163. 10.1016/j.jbiomech.2012.05.037.

(48) Dembia, C. L.; Bianco, N. A.; Falisse, A.; Hicks, J. L.; Delp, S. L. OpenSim Moco: Musculoskeletal Optimal Control. PLOS Comput. Biol. 2020, 16 (12), e1008493. 10.1371/journal.pcbi.1008493.

(49) Delp, S. L.; Anderson, F. C.; Arnold, A. S.; Loan, P.; Habib, A.; John, C. T.; Guendelman, E.; Thelen, D. G. OpenSim: Open-Source Software to Create and Analyze Dynamic Simulations of Movement. IEEE Trans. Biomed. Eng. 2007, 54 (11), 1940–1950. 10.1109/TBME.2007.901024.

(50) Seth, A.; Hicks, J. L.; Uchida, T. K.; Habib, A.; Dembia, C. L.; Dunne, J. J.; Ong, C. F.; DeMers, M. S.; Rajagopal, A.; Millard, M.; Hamner, S. R.; Arnold, E. M.; Yong, J. R.; Lakshmikanth, S. K.; Sherman, M. A.; Ku, J. P.; Delp, S. L. OpenSim: Simulating Musculoskeletal Dynamics and Neuromuscular Control to Study Human and Animal Movement. PLOS Comput. Biol. 2018, 14 (7), e1006223. 10.1371/journal.pcbi.1006223.

(51) Pitto, L.; Kainz, H.; Falisse, A.; Wesseling, M.; Van Rossom, S.; Hoang, H.; Papageorgiou, E.; Hallemans, A.; Desloovere, K.; Molenaers, G.; Van Campenhout, A.; De Groote, F.; Jonkers, I. SimCP: A Simulation Platform to Predict Gait Performance Following Orthopedic Intervention in Children With Cerebral Palsy. Front. Neurorobotics 2019, 13. 10.3389/fnbot.2019.00054.

(52) Valente, G.; Crimi, G.; Vanella, N.; Schileo, E.; Taddei, F. nmsBuilder: Freeware to Create Subject-Specific Musculoskeletal Models for OpenSim. Comput. Methods Programs Biomed. 2017, 152, 85–92. 10.1016/j.cmpb.2017.09.012.

(53) Zhang, J.; Sorby, H.; Clement, J.; Thomas, C. D. L.; Hunter, P.; Nielsen, P.; Lloyd, D.; Taylor, M.; Besier, T. The MAP Client: User-Friendly Musculoskeletal Modelling Workflows. In Biomedical Simulation; Bello, F., Cotin, S., Eds.; Springer International Publishing: Cham, 2014; pp 182–192. 10.1007/978-3-319-12057-7_21.

(54) Veerkamp, K.; Kainz, H.; Killen, B. A.; Jónasdóttir, H.; van der Krogt, M. M. Torsion Tool: An Automated Tool for Personalising Femoral and Tibial Geometries in OpenSim Musculoskeletal Models. J. Biomech. 2021, 125, 110589. 10.1016/j.jbiomech.2021.110589.

(55) Modenese, L.; Barzan, M.; Carty, C. P. Dependency of Lower Limb Joint Reaction Forces on Femoral Version. Gait Posture 2021, 88, 318–321. 10.1016/j.gaitpost.2021.06.014.

(56) Werling, K.; Bianco, N. A.; Raitor, M.; Stingel, J.; Hicks, J. L.; Collins, S. H.; Delp, S. L.; Liu, C. K. AddBiomechanics: Automating Model Scaling, Inverse Kinematics, and Inverse Dynamics from Human Motion Data through Sequential Optimization. PLOS ONE 2023, 18 (11), e0295152. 10.1371/journal.pone.0295152.

(57) Kewley, A.; Beesel, J.; Seth, A. OpenSim Creator, 2025. 10.5281/zenodo.14755649.

(58) Di Pietro, A.; Bersani, A.; Curreli, C.; Di Puccio, F. AST: An OpenSim-Based Tool for the Automatic Scaling of Generic Musculoskeletal Models. Comput. Biol. Med. 2024, 175, 108524. 10.1016/j.compbiomed.2024.108524.

(59) van den Bogert, A. J.; Blana, D.; Heinrich, D. Implicit Methods for Efficient Musculoskeletal Simulation and Optimal Control. Procedia IUTAM 2011, 2 (2011), 297–316. 10.1016/j.piutam.2011.04.027.

(60) Patterson, M. A.; Rao, A. V. GPOPS-II: A MATLAB Software for Solving Multiple-Phase Optimal Control Problems Using Hp-Adaptive Gaussian Quadrature Collocation Methods and Sparse Nonlinear Programming. ACM Trans Math Softw 2014, 41 (1), 1:1-1:37. 10.1145/2558904.

(61) De Groote, F.; Kinney, A. L.; Rao, A. V.; Fregly, B. J. Evaluation of Direct Collocation Optimal Control Problem Formulations for Solving the Muscle Redundancy Problem. Ann. Biomed. Eng. 2016, 44 (10), 2922–2936. 10.1007/s10439-016-1591-9.

(62) Febrer-Nafría, M.; Pallarès-López, R.; Fregly, B. J.; Font-Llagunes, J. M. Comparison of Different Optimal Control Formulations for Generating Dynamically Consistent Crutch Walking Simulations Using a Torque-Driven Model. Mech. Mach. Theory 2020, 154, 104031. 10.1016/j.mechmachtheory.2020.104031.

(63) Butler, D. Translational Research: Crossing the Valley of Death. Nature 2008, 453 (7197), 840–842. 10.1038/453840a.

(64) Jørgensen, J. T. Twenty Years with Personalized Medicine: Past, Present, and Future of Individualized Pharmacotherapy. The Oncologist 2019, 24 (7), e432–e440. 10.1634/theoncologist.2019-0054.

(65) Rassmussen J; Vondrak V; Damsgaard M; de Zee M; Christensen ST. The AnyBody Project – Computer Analysis of the Human Body. Comput. Anal. Hum. Body 2002, No. Biomechanics of Man, 270–274.

(66) Wächter, A.; Biegler, L. T. On the Implementation of an Interior-Point Filter Line-Search Algorithm for Large-Scale Nonlinear Programming. Math. Program. 2006, 106 (1), 25–57. 10.1007/s10107-004-0559-y.

(67) Sauder, N. R.; Meyer, A. J.; Allen, J. L.; Ting, L. H.; Kesar, T. M.; Fregly, B. J. Computational Design of FastFES Treatment to Improve Propulsive Force Symmetry During Post-Stroke Gait: A Feasibility Study. Front. Neurorobotics 2019, 13, 80. 10.3389/fnbot.2019.00080.

(68) Vega, M. M.; Li, G.; Shourijeh, M. S.; Ao, D.; Weinschenk, R. C.; Patten, C.; Font-Llagunes, J. M.; Lewis, V. O.; Fregly, B. J. Computational Evaluation of Psoas Muscle Influence on Walking Function Following Internal Hemipelvectomy with Reconstruction. Front. Bioeng. Biotechnol. 2022, 10, 855870. 10.3389/fbioe.2022.855870.

(69) McMorland, A. J. C.; Runnalls, K. D.; Byblow, W. D. A Neuroanatomical Framework for Upper Limb Synergies after Stroke. Front. Hum. Neurosci. 2015, 9. 10.3389/fnhum.2015.00082.

(70) Akhras, M. A.; Bortoletto, R.; Madehkhaksar, F.; Tagliapietra, L. Neural and Musculoskeletal Modeling: Its Role in Neurorehabilitation. In Emerging Therapies in Neurorehabilitation II; Pons, J. L., Raya, R., González, J., Eds.; Springer International Publishing: Cham, 2016; pp 109–143. 10.1007/978-3-319-24901-8_5.

(71) Meyer, A. J.; Patten, C.; Fregly, B. J. Lower Extremity EMG-Driven Modeling of Walking with Automated Adjustment of Musculoskeletal Geometry. PLOS ONE 2017, 12 (7), e0179698. 10.1371/journal.pone.0179698.

(72) Reinbolt, J. A.; Haftka, R. T.; Chmielewski, T. L.; Fregly, B. J. Are Patient-Specific Joint and Inertial Parameters Necessary for Accurate Inverse Dynamics Analyses of Gait? IEEE Trans. Biomed. Eng. 2007, 54 (5), 782–793. 10.1109/TBME.2006.889187.

(73) Serrancolí, G.; Kinney, A. L.; Fregly, B. J.; Font-Llagunes, J. M. Neuromusculoskeletal Model Calibration Significantly Affects Predicted Knee Contact Forces for Walking. J. Biomech. Eng. 2016, 138 (8), 0810011–08100111. 10.1115/1.4033673.

(74) Żuk, M.; Syczewska, M.; Pezowicz, C. Influence of Uncertainty in Selected Musculoskeletal Model Parameters on Muscle Forces Estimated in Inverse Dynamics-Based Static Optimization and Hybrid Approach. J. Biomech. Eng. 2018, 140 (121001). 10.1115/1.4040943.

(75) Lloyd, D. G.; Besier, T. F. An EMG-Driven Musculoskeletal Model to Estimate Muscle Forces and Knee Joint Moments in Vivo. J. Biomech. 2003, 36 (6), 765–776. 10.1016/s0021-9290(03)00010-1.

(76) Manal, K.; Buchanan, T. S. Use of an EMG-Driven Biomechanical Model to Study Virtual Injuries. Med. Sci. Sports Exerc. 2005, 37 (11), 1917–1923. 10.1249/01.mss.0000176685.35442.6b.

(77) Moissenet, F.; Modenese, L.; Dumas, R. Alterations of Musculoskeletal Models for a More Accurate Estimation of Lower Limb Joint Contact Forces during Normal Gait: A Systematic Review. J. Biomech. 2017, 63, 8–20. 10.1016/j.jbiomech.2017.08.025.

(78) Arones, M. M.; Shourijeh, M. S.; Patten, C.; Fregly, B. J. Musculoskeletal Model Personalization Affects Metabolic Cost Estimates for Walking. Front. Bioeng. Biotechnol. 2020, 8, 588925. 10.3389/fbioe.2020.588925.

(79) Jackson, J. N.; Hass, C. J.; Fregly, B. J. Development of a Subject-Specific Foot-Ground Contact Model for Walking. J. Biomech. Eng. 2016, 138 (9), 0910021–09100212. 10.1115/1.4034060.

(80) Nguyen, V. Q.; Johnson, R. T.; Sup, F. C.; Umberger, B. R. Bilevel Optimization for Cost Function Determination in Dynamic Simulation of Human Gait. IEEE Trans. Neural Syst. Rehabil. Eng. Publ. IEEE Eng. Med. Biol. Soc. 2019, 27 (7), 1426–1435. 10.1109/TNSRE.2019.2922942.

(81) Kuska, E. C.; Steele, K. M. Does Crouch Alter the Effects of Neuromuscular Impairments on Gait? A Simulation Study. J. Biomech. 2024, 165, 112015. 10.1016/j.jbiomech.2024.112015.

(82) Pariser, K. M.; Higginson, J. S. Development and Validation of a Framework for Predictive Simulation of Treadmill Gait. J. Biomech. Eng. 2022, 144 (11), 114505. 10.1115/1.4054867.

(83) Falisse, A.; Serrancolí, G.; Dembia, C. L.; Gillis, J.; Jonkers, I.; De Groote, F. Rapid Predictive Simulations with Complex Musculoskeletal Models Suggest That Diverse Healthy and Pathological Human Gaits Can Emerge from Similar Control Strategies. J. R. Soc. Interface 2019, 16 (157), 20190402. 10.1098/rsif.2019.0402.

(84) Dorschky, E.; Nitschke, M.; Seifer, A.-K.; van den Bogert, A. J.; Eskofier, B. M. Estimation of Gait Kinematics and Kinetics from Inertial Sensor Data Using Optimal Control of Musculoskeletal Models. J. Biomech. 2019, 95, 109278. 10.1016/j.jbiomech.2019.07.022.

(85) Ackermann, M.; van den Bogert, A. J. Optimality Principles for Model-Based Prediction of Human Gait. J. Biomech. 2010, 43 (6), 1055–1060. 10.1016/j.jbiomech.2009.12.012.

(86) Johnson, R. T.; Bianco, N. A.; Finley, J. M. Patterns of Asymmetry and Energy Cost Generated from Predictive Simulations of Hemiparetic Gait. PLoS Comput. Biol. 2022, 18 (9), e1010466. 10.1371/journal.pcbi.1010466.

(87) Miller, R. H.; Russell Esposito, E. Transtibial Limb Loss Does Not Increase Metabolic Cost in Three-Dimensional Computer Simulations of Human Walking. PeerJ 2021, 9, e11960. 10.7717/peerj.11960.

(88) Weng, J.; Hashemi, E.; Arami, A. Adaptive Reference Inverse Optimal Control for Natural Walking With Musculoskeletal Models. IEEE Trans. Neural Syst. Rehabil. Eng. 2022, 30, 1567–1575. 10.1109/TNSRE.2022.3180690.

(89) Tresch, M. C.; Saltiel, P.; Bizzi, E. The Construction of Movement by the Spinal Cord. Nat. Neurosci. 1999, 2 (2), 162–167. 10.1038/5721.

(90) Lee, D.; Seung, H. Algorithms for Non-Negative Matrix Factorization. Adv Neural Inf. Process Syst 2001, 13.

(91) Ivanenko, Y. P.; Poppele, R. E.; Lacquaniti, F. Five Basic Muscle Activation Patterns Account for Muscle Activity during Human Locomotion. J. Physiol. 2004, 556 (Pt 1), 267–282. 10.1113/jphysiol.2003.057174.

(92) Cheung, V. C. K.; Piron, L.; Agostini, M.; Silvoni, S.; Turolla, A.; Bizzi, E. Stability of Muscle Synergies for Voluntary Actions after Cortical Stroke in Humans. Proc. Natl. Acad. Sci. U. S. A. 2009, 106 (46), 19563. 10.1073/pnas.0910114106.

(93) Clark, D. J.; Ting, L. H.; Zajac, F. E.; Neptune, R. R.; Kautz, S. A. Merging of Healthy Motor Modules Predicts Reduced Locomotor Performance and Muscle Coordination Complexity Post-Stroke. J. Neurophysiol. 2010, 103 (2), 844–857. 10.1152/jn.00825.2009.

(94) Gizzi, L.; Nielsen, J. F.; Felici, F.; Ivanenko, Y. P.; Farina, D. Impulses of Activation but Not Motor Modules Are Preserved in the Locomotion of Subacute Stroke Patients. J. Neurophysiol. 2011, 106 (1), 202–210. 10.1152/jn.00727.2010.

(95) Routson, R. L.; Clark, D. J.; Bowden, M. G.; Kautz, S. A.; Neptune, R. R. The Influence of Locomotor Rehabilitation on Module Quality and Post-Stroke Hemiparetic Walking Performance. Gait Posture 2013, 38 (3), 511–517. 10.1016/j.gaitpost.2013.01.020.

(96) Roh, J.; Rymer, W. Z.; Perreault, E. J.; Yoo, S. B.; Beer, R. F. Alterations in Upper Limb Muscle Synergy Structure in Chronic Stroke Survivors. J. Neurophysiol. 2013, 109 (3), 768–781. 10.1152/jn.00670.2012.

(97) Pitto, L.; Rossom, S. van; Desloovere, K.; Molenaers, G.; Huenaerts, C.; Groote, F. D.; Jonkers, I. Pre-Treatment EMG Can Be Used to Model Post-Treatment Muscle Coordination during Walking in Children with Cerebral Palsy. PLOS ONE 2020, 15 (2), e0228851. 10.1371/journal.pone.0228851.

(98) Rabbi, M. F.; Pizzolato, C.; Lloyd, D. G.; Carty, C. P.; Devaprakash, D.; Diamond, L. E. Non-Negative Matrix Factorisation Is the Most Appropriate Method for Extraction of Muscle Synergies in Walking and Running. Sci. Rep. 2020, 10 (1), 8266. 10.1038/s41598-020-65257-w.

(99) Rajagopal, A.; Dembia, C. L.; DeMers, M. S.; Delp, D. D.; Hicks, J. L.; Delp, S. L. Full-Body Musculoskeletal Model for Muscle-Driven Simulation of Human Gait. IEEE Trans. Biomed. Eng. 2016, 63 (10), 2068–2079. 10.1109/TBME.2016.2586891.

(100) van den Bogert, A. J.; Smith, G. D.; Nigg, B. M. In Vivo Determination of the Anatomical Axes of the Ankle Joint Complex: An Optimization Approach. J. Biomech. 1994, 27 (12), 1477–1488. 10.1016/0021-9290(94)90197-x.

(101) Reinbolt, J. A.; Schutte, J. F.; Fregly, B. J.; Koh, B. I.; Haftka, R. T.; George, A. D.; Mitchell, K. H. Determination of Patient-Specific Multi-Joint Kinematic Models through Two-Level Optimization. J. Biomech. 2005, 38 (3), 621–626. 10.1016/j.jbiomech.2004.03.031.

(102) Price, M. A.; LaPrè, A. K.; Johnson, R. T.; Umberger, B. R.; Sup IV, F. C. A Model-Based Motion Capture Marker Location Refinement Approach Using Inverse Kinematics from Dynamic Trials. Int. J. Numer. Methods Biomed. Eng. 2020, 36 (1), e3283. 10.1002/cnm.3283.

(103) Xu, D.; Carlton, L. G.; Rosengren, K. S. Anticipatory Postural Adjustments for Altering Direction during Walking. J. Mot. Behav. 2004, 36 (3), 316–326. 10.3200/JMBR.36.3.316-326.

(104) Bell, A. L.; Pedersen, D. R.; Brand, R. A. A Comparison of the Accuracy of Several Hip Center Location Prediction Methods. J. Biomech. 1990, 23 (6), 617–621. 10.1016/0021-9290(90)90054-7.

(105) Leardini, A.; Cappozzo, A.; Catani, F.; Toksvig-Larsen, S.; Petitto, A.; Sforza, V.; Cassanelli, G.; Giannini, S. Validation of a Functional Method for the Estimation of Hip Joint Centre Location. J. Biomech. 1999, 32 (1), 99–103. 10.1016/S0021-9290(98)00148-1.

(106) Charlton, I. W.; Tate, P.; Smyth, P.; Roren, L. Repeatability of an Optimised Lower Body Model. Gait Posture 2004, 20 (2), 213–221. 10.1016/j.gaitpost.2003.09.004.

(107) Chèze, L.; Fregly, B. J.; Dimnet, J. Determination of Joint Functional Axes from Noisy Marker Data Using the Finite Helical Axis. Hum. Mov. Sci. 1998, 17 (1), 1–15. 10.1016/S0167-9457(97)00018-3.

(108) Sartori, M.; Reggiani, M.; Farina, D.; Lloyd, D. G. EMG-Driven Forward-Dynamic Estimation of Muscle Force and Joint Moment about Multiple Degrees of Freedom in the Human Lower Extremity. PLOS ONE 2012, 7 (12), e52618. 10.1371/journal.pone.0052618.

(109) Pizzolato, C.; Lloyd, D. G.; Sartori, M.; Ceseracciu, E.; Besier, T. F.; Fregly, B. J.; Reggiani, M. CEINMS: A Toolbox to Investigate the Influence of Different Neural Control Solutions on the Prediction of Muscle Excitation and Joint Moments during Dynamic Motor Tasks. J. Biomech. 2015, 48 (14), 3929–3936. 10.1016/j.jbiomech.2015.09.021.

(110) De Groote, F.; Van Campen, A.; Jonkers, I.; De Schutter, J. Sensitivity of Dynamic Simulations of Gait and Dynamometer Experiments to Hill Muscle Model Parameters of Knee Flexors and Extensors. J. Biomech. 2010, 43 (10), 1876–1883. 10.1016/j.jbiomech.2010.03.022.

(111) Anderson, F. C.; Pandy, M. G. Static and Dynamic Optimization Solutions for Gait Are Practically Equivalent. J. Biomech. 2001, 34 (2), 153–161. 10.1016/S0021-9290(00)00155-X.

(112) Millard, M.; Uchida, T.; Seth, A.; Delp, S. L. Flexing Computational Muscle: Modeling and Simulation of Musculotendon Dynamics. J. Biomech. Eng. 2013, 135 (2), 0210051–02100511. 10.1115/1.4023390.

(113) Buchanan, T. S.; Lloyd, D. G.; Manal, K.; Besier, T. F. Estimation of Muscle Forces and Joint Moments Using a Forward-Inverse Dynamics Model. Med. Sci. Sports Exerc. 2005, 37 (11), 1911– 1916. 10.1249/01.mss.0000176684.24008.6f.

(114) Handsfield, G. G.; Meyer, C. H.; Hart, J. M.; Abel, M. F.; Blemker, S. S. Relationships of 35 Lower Limb Muscles to Height and Body Mass Quantified Using MRI. J. Biomech. 2014, 47 (3), 631–638. 10.1016/j.jbiomech.2013.12.002.

(115) Ao, D.; Shourijeh, M. S.; Patten, C.; Fregly, B. J. Evaluation of Synergy Extrapolation for Predicting Unmeasured Muscle Excitations from Measured Muscle Synergies. Front. Comput. Neurosci. 2020, 14. 10.3389/fncom.2020.588943.

(116) Ao, D.; Vega, M. M.; Shourijeh, M. S.; Patten, C.; Fregly, B. J. EMG-Driven Musculoskeletal Model Calibration with Estimation of Unmeasured Muscle Excitations via Synergy Extrapolation. Front. Bioeng. Biotechnol. 2022, 10. 10.3389/fbioe.2022.962959.

(117) Di, A.; Benjamin, J. F. Comparison of Synergy Extrapolation and Static Optimization for Estimating Multiple Unmeasured Muscle Activations during Walking. bioRxiv 2024, 2024.03.03.583228. 10.1101/2024.03.03.583228.

(118) Arnold, E. M.; Hamner, S. R.; Seth, A.; Millard, M.; Delp, S. L. How Muscle Fiber Lengths and Velocities Affect Muscle Force Generation as Humans Walk and Run at Different Speeds. J. Exp. Biol. 2013, 216 (11), 2150–2160. 10.1242/jeb.075697.

(119) Gopalakrishnan, A.; Modenese, L.; Phillips, A. T. M. A Novel Computational Framework for Deducing Muscle Synergies from Experimental Joint Moments. Front. Comput. Neurosci. 2014, 8. 10.3389/fncom.2014.00153.

(120) Shourijeh, M. S.; Fregly, B. J. Muscle Synergies Modify Optimization Estimates of Joint Stiffness During Walking. J. Biomech. Eng. 2020, 142 (1), 011011. 10.1115/1.4044310.

(121) Ivanenko, Y. P.; Cappellini, G.; Dominici, N.; Poppele, R. E.; Lacquaniti, F. Coordination of Locomotion with Voluntary Movements in Humans. J. Neurosci. 2005, 25 (31), 7238–7253. 10.1523/JNEUROSCI.1327-05.2005.

(122) Rodriguez, K. L.; Roemmich, R. T.; Cam, B.; Fregly, B. J.; Hass, C. J. Persons with Parkinson’s Disease Exhibit Decreased Neuromuscular Complexity during Gait. Clin. Neurophysiol. 2013, 124 (7), 1390–1397. 10.1016/j.clinph.2013.02.006.

(123) Bhargava, L. J.; Pandy, M. G.; Anderson, F. C. A Phenomenological Model for Estimating Metabolic Energy Consumption in Muscle Contraction. J. Biomech. 2004, 37 (1), 81–88. 10.1016/s0021-9290(03)00239-2.

(124) Roh, J.; Cheung, V. C. K.; Bizzi, E. Modules in the Brain Stem and Spinal Cord Underlying Motor Behaviors. J. Neurophysiol. 2011, 106 (3), 1363–1378. 10.1152/jn.00842.2010.

(125) Cheung, V. C. K.; Piron, L.; Agostini, M.; Silvoni, S.; Turolla, A.; Bizzi, E. Stability of Muscle Synergies for Voluntary Actions after Cortical Stroke in Humans. Proc. Natl. Acad. Sci. 2009, 106 (46), 19563–19568. 10.1073/pnas.0910114106.

(126) Raja, B.; Neptune, R. R.; Kautz, S. A. Coordination of the Non-Paretic Leg During Hemiparetic Gait: Expected and Novel Compensatory Patterns. Clin. Biomech. Bristol Avon 2012, 27 (10), 1023–1030. 10.1016/j.clinbiomech.2012.08.005.

(127) Kautz, S. A.; Patten, C. Interlimb Influences on Paretic Leg Function in Poststroke Hemiparesis. J. Neurophysiol. 2005, 93 (5), 2460–2473. 10.1152/jn.00963.2004.

(128) Sánchez, N.; Acosta, A. M.; López-Rosado, R.; Dewald, J. P. A. Neural Constraints Affect the Ability to Generate Hip Abduction Torques When Combined With Hip Extension or Ankle Plantarflexion in Chronic Hemiparetic Stroke. Front. Neurol. 2018, 9. 10.3389/fneur.2018.00564.

(129) Pandian, S.; Arya, K. N.; Kumar, D. Does Motor Training of the Nonparetic Side Influences Balance and Function in Chronic Stroke? A Pilot RCT. Sci. World J. 2014, 2014 (1), 769726. 10.1155/2014/769726.

(130) Lim, H.; Madhavan, S. Non-Paretic Leg Movements Can Facilitate Cortical Drive to the Paretic Leg in Individuals Post Stroke with Severe Motor Impairment: Implications for Motor Priming. Eur. J. Neurosci. 2023, 58 (3), 2853–2867. 10.1111/ejn.16069.

(131) Kramer, S.; Johnson, L.; Bernhardt, J.; Cumming, T. Energy Expenditure and Cost During Walking After Stroke: A Systematic Review. Arch. Phys. Med. Rehabil. 2016, 97 (4), 619–632.e1. 10.1016/j.apmr.2015.11.007.

(132) Hoxie, R. E.; Rubenstein, L. Z.; Hoenig, H.; Gallagher, B. R. The Older Pedestrian. J. Am. Geriatr. Soc. 1994, 42 (4), 444–450. 10.1111/j.1532-5415.1994.tb07496.x.

(133) Perry, J.; Garrett, M.; Gronley, J. K.; Mulroy, S. J. Classification of Walking Handicap in the Stroke Population. Stroke 1995, 26 (6), 982–989. 10.1161/01.str.26.6.982.

(134) Hug, F.; Turpin, N. A.; Couturier, A.; Dorel, S. Consistency of Muscle Synergies during Pedaling across Different Mechanical Constraints. J. Neurophysiol. 2011, 106 (1), 91–103. 10.1152/jn.01096.2010.

(135) Pataky, T. C.; Goulermas, J. Y. Pedobarographic Statistical Parametric Mapping (pSPM): A Pixel-Level Approach to Foot Pressure Image Analysis. J. Biomech. 2008, 41 (10), 2136–2143. 10.1016/j.jbiomech.2008.04.034.

(136) Duckworth, T.; Betts, R. P.; Franks, C. I.; Burke, J. The Measurement of Pressures under the Foot. Foot Ankle 1982, 3 (3), 130–141. 10.1177/107110078200300303.

(137) Li, G.; Ao, D.; Vega, M. M.; Zandiyeh, P.; Chang, S.-H.; Penny, A. N.; Lewis, V. O.; Fregly, B. J. Changes in Walking Function and Neural Control Following Pelvic Cancer Surgery with Reconstruction. Front. Bioeng. Biotechnol. 2024, 12. 10.3389/fbioe.2024.1389031.

(138) Wirth, M. A.; Kovesi, P. MATLAB as an Introductory Programming Language. Comput. Appl. Eng. Educ. 2006, 14 (1), 20–30. 10.1002/cae.20064.

(139) Vicéns, J. L.; Zamora, B.; Ojados, D. Improvement of the Reflective Learning in Engineering Education Using MATLAB for Problems Solving. Comput. Appl. Eng. Educ. 2016, 24 (5), 755–764. 10.1002/cae.21748.

(140) Anderson, F. C.; Pandy, M. G. Static and Dynamic Optimization Solutions for Gait Are Practically Equivalent. J. Biomech. 2001, 34 (2), 153–161. 10.1016/s0021-9290(00)00155-x.

(141) Suwarganda, E. K.; Diamond, L. E.; Lloyd, D. G.; Besier, T. F.; Zhang, J.; Killen, B. A.; Savage, T. N.; Saxby, D. J. Minimal Medical Imaging Can Accurately Reconstruct Geometric Bone Models for Musculoskeletal Models. PLOS ONE 2019, 14 (2), e0205628. 10.1371/journal.pone.0205628.

(142) Killen, B. A.; Brito da Luz, S.; Lloyd, D. G.; Carleton, A. D.; Zhang, J.; Besier, T. F.; Saxby, D. J. Automated Creation and Tuning of Personalised Muscle Paths for OpenSim Musculoskeletal Models of the Knee Joint. Biomech. Model. Mechanobiol. 2021, 20 (2), 521–533. 10.1007/s10237-020-01398-1.

(143) Kainz, H.; Wesseling, M.; Jonkers, I. Generic Scaled versus Subject-Specific Models for the Calculation of Musculoskeletal Loading in Cerebral Palsy Gait: Effect of Personalized Musculoskeletal Geometry Outweighs the Effect of Personalized Neural Control. Clin. Biomech. 2021, 87, 105402. 10.1016/j.clinbiomech.2021.105402.

(144) Michaud, F.; Frey-Law, L. A.; Lugrís, U.; Cuadrado, L.; Figueroa-Rodríguez, J.; Cuadrado, J. Applying a Muscle Fatigue Model When Optimizing Load-Sharing between Muscles for Short-Duration High-Intensity Exercise: A Preliminary Study. Front. Physiol. 2023, 14. 10.3389/fphys.2023.1167748.

(145) Ma, L.; Zhang, W.; Hu, B.; Chablat, D.; Bennis, F.; Guillaume, F. Determination of Subject-Specific Muscle Fatigue Rates under Static Fatiguing Operations. Ergonomics 2013, 56 (12), 1889–1900. 10.1080/00140139.2013.851283.

