## Supplementary material for "The Neuromusculoskeletal Modeling Pipeline: MATLAB-based Model Personalization and Treatment Optimization Functionality for OpenSim": JointMomentErrors%MaxID.pdf

**Lower Body Joint Moment Matching Errors (% maximum ID moment) following Muscle-tendon Model Personalization**

| Joint Coordinate | Hip Flexion | Hip Adduction | Hip Rotation | Knee Angle | Ankle Angle | Subtalar Angle |
|---|---|---|---|---|---|---|
| Right (% max ID) | 3.9 | 6.1 | 36.4 | 4.8 | 1.2 | 5.6 |
| Left (% max ID) | 9.1 | 6.1 | 30.3 | 6.6 | 2.3 | 5.7 |
