## Supplementary material for "The Neuromusculoskeletal Modeling Pipeline: MATLAB-based Model Personalization and Treatment Optimization Functionality for OpenSim": Table S1. Cost Terms.pdf

**Table S1. Summary of available cost function terms for Tracking, Verification, and Design Optimization.**

**Tracking Optimization**

| *Cost Term* | *Component Type* | *Max Allowable Error* | *Error Center* |
|---|---|---|---|
| generalized_coordinate_tracking | coordinate | Y | N |
| generalized_speed_tracking | coordinate | Y | N |
| joint_acceleration_minimization | coordinate | Y | N |
| marker_position_tracking | marker | Y | N |
| inverse_dynamics_load_tracking | load | Y | N |
| inverse_dynamics_load_minimization | load | Y | N |
| inverse_dynamics_slope_tracking | load | Y | N |
| external_force_tracking | force | Y | N |
| external_moment_tracking | moment | Y | N |
| muscle_activation_tracking | muscle | Y | N |
| controller_slope_minimization | controller | Y | N |

**Verification Optimization**

| *Cost Term* | *Component Type* | *Max Allowable Error* | *Error Center* |
|---|---|---|---|
| generalized_coordinate_tracking | coordinate | Y | N |
| generalized_speed_tracking | coordinate | Y | N |
| joint_acceleration_minimization | coordinate | Y | N |
| marker_position_tracking | marker | Y | N |
| controller_tracking | controller | Y | N |
| controller_slope_minimization | controller | Y | N |
| controller_frequency_minimization | controller | Y | N |

**Design Optimization**

| *Cost Term* | *Component Type* | *Max Allowable Error* | *Error Center* |
|---|---|---|---|
| generalized_coordinate_tracking | coordinate | Y | N |
| generalized_speed_tracking | coordinate | Y | N |
| joint_acceleration_minimization | coordinate | Y | N |
| joint_power_minimization | coordinate | Y | N |
| joint_energy_generation_goal | coordinate | Y | Y |
| joint_energy_absorption_goal | coordinate | Y | Y |
| marker_position_tracking | marker | Y | N |
| inverse_dynamics_load_tracking | load | Y | N |
| inverse_dynamics_slope_tracking | load | Y | N |
| external_force_tracking | force | Y | N |
| external_moment_tracking | moment | Y | N |
| muscle_activation_tracking | muscle | Y | N |
| muscle_activation_minimization | muscle | Y | N |
| controller_tracking | controller | Y | N |
| controller_slope_minimization | controller | Y | N |
| controller_frequency_minimization | controller | Y | N |
| controller_shape_tracking | controller | Y | N |
| angular_momentum_minimization | None | Y | N |
| synergy_vector_tracking | None | Y | Y |
| belt_speed_goal | None | Y | Y |
| relative_walking_speed_goal | None | Y | Y |
| relative_metabolic_cost_per_time | None | Y | Y |
| relative_metabolic_cost_per_distance | None | Y | Y |
| propulsive_impulse_goal | None | Y | Y |
| braking_impulse_goal | None | Y | Y |
| user_defined | Any | Y | Y |
