## Supplementary material for "The Neuromusculoskeletal Modeling Pipeline: MATLAB-based Model Personalization and Treatment Optimization Functionality for OpenSim": Table S2. Constraint Terms.pdf

**Table S2. Summary of available constraint terms for Tracking, Verification, and Design Optimization.**

**Tracking Optimization**

| *Constraint Term* | *Component Type* |
|---|---|
| state_position_periodicity | coordinate |
| state_velocity_periodicity | coordinate |
| kinetic_consistency | load |
| root_segment_residual_load | load |
| root_segment_residual_load_periodicity | load |
| external_force_periodicity | force |
| external_moment_periodicity | moment |
| synergy_weight_sum | synergy_group |
| synergy_weight_magnitude | synergy_group |

**Verification Optimization**

| *Constraint Term* | *Component Type* |
|---|---|
| state_position_periodicity | coordinate |
| state_velocity_periodicity | coordinate |
| kinetic_consistency | load |
| root_segment_residual_load | load |
| root_segment_residual_load_periodicity | load |
| external_force_periodicity | force |
| external_moment_periodicity | moment |

**Design Optimization**

| *Constraint Term* | *Component Type* |
|---|---|
| initial_state_position | coordinate |
| final_state_position | coordinate |
| final_state_velocity | coordinate |
| state_position_periodicity | coordinate |
| state_velocity_periodicity | coordinate |
| kinetic_consistency | load |
| root_segment_residual_load | load |
| root_segment_residual_load_periodicity | load |
| external_force_periodicity | force |
| external_moment_periodicity | moment |
| limit_muscle_activation | muscle |
| limit_normalized_fiber_length | muscle |
| synergy_weight_sum | synergy_group |
| synergy_weight_magnitude | synergy_group |
