## Supplementary figures and images for "The Neuromusculoskeletal Modeling Pipeline: MATLAB-based Model Personalization and Treatment Optimization Functionality for OpenSim"

### activations.png

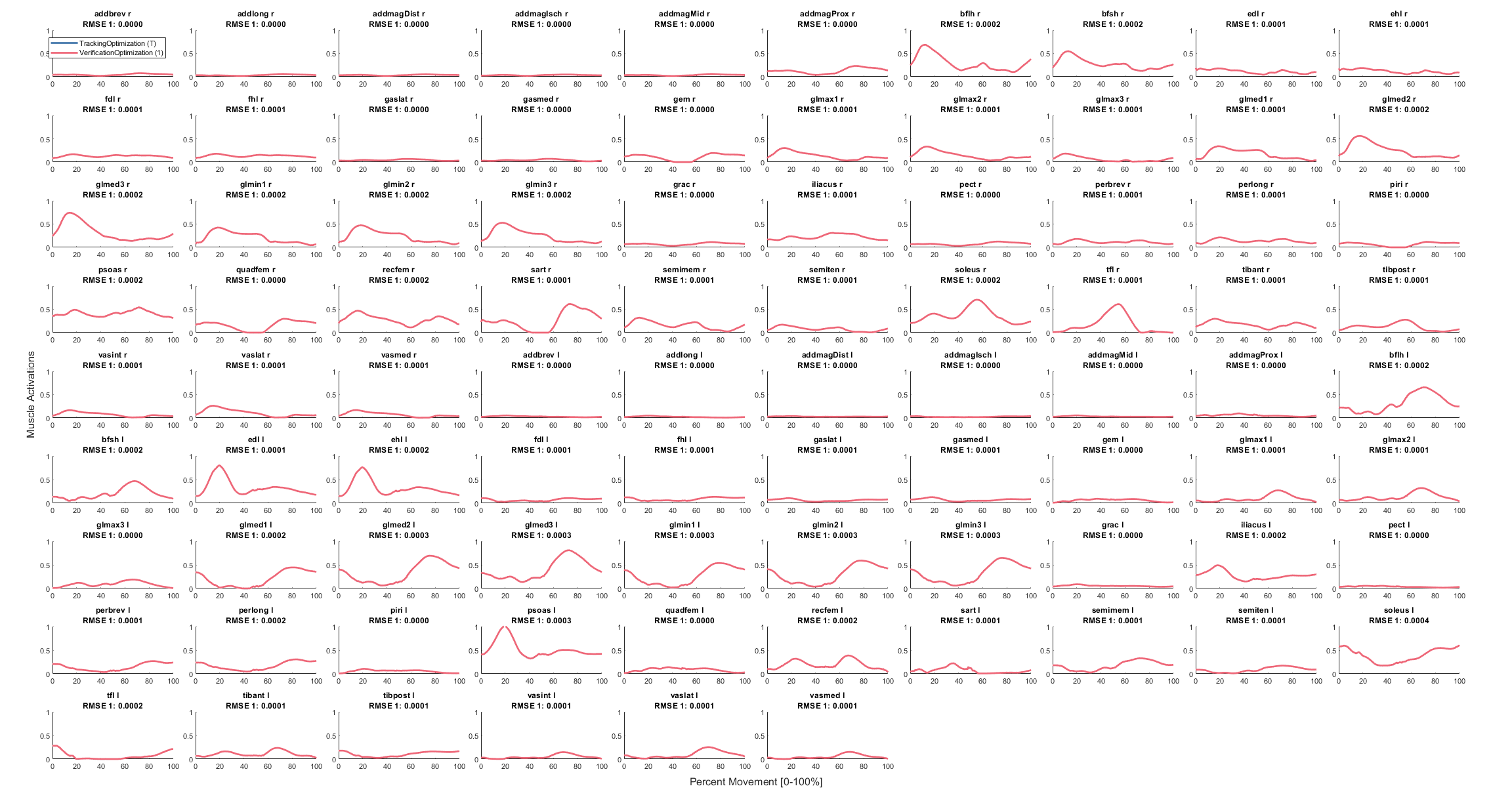

### activations.png

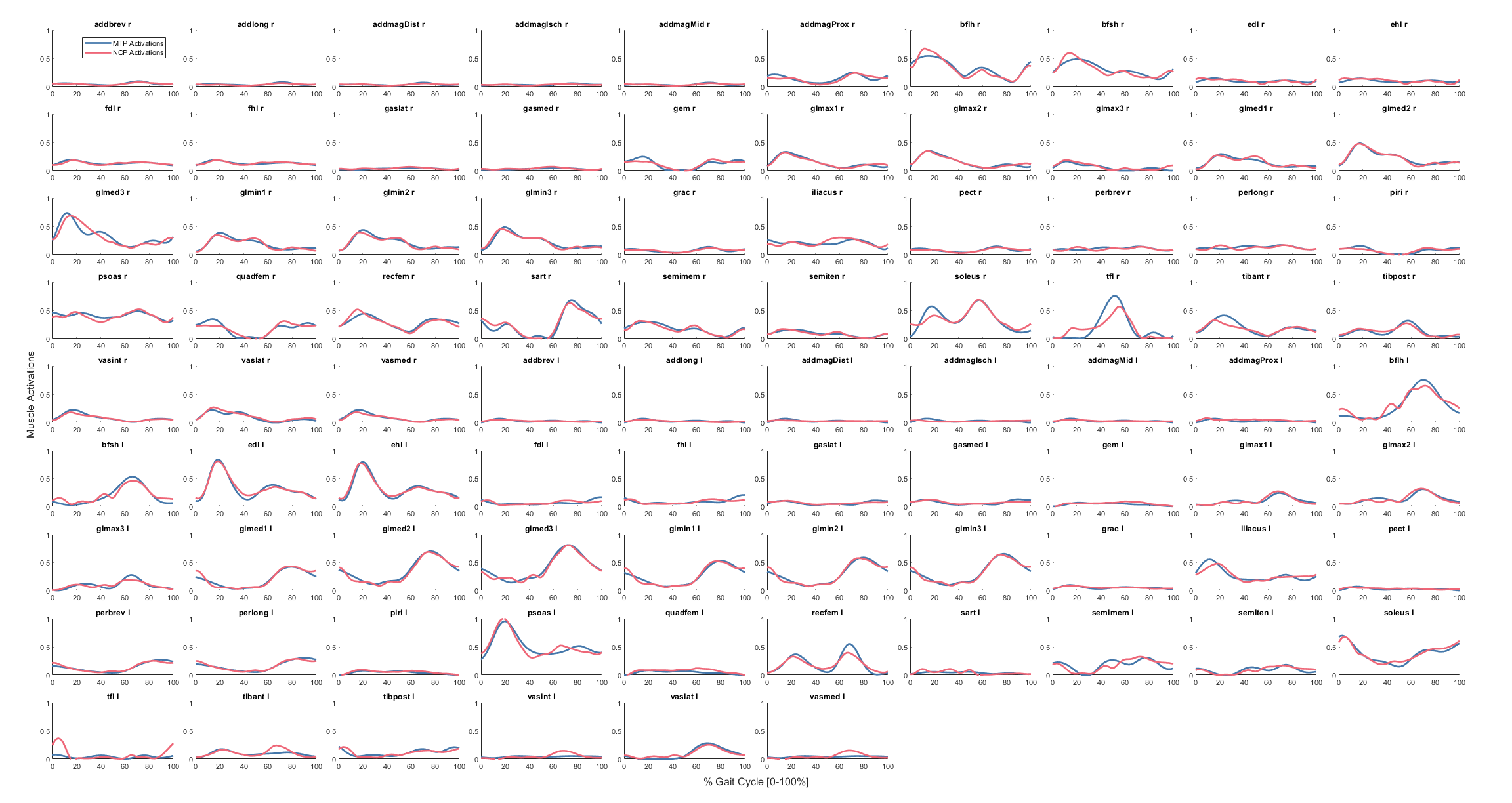

### activations.png

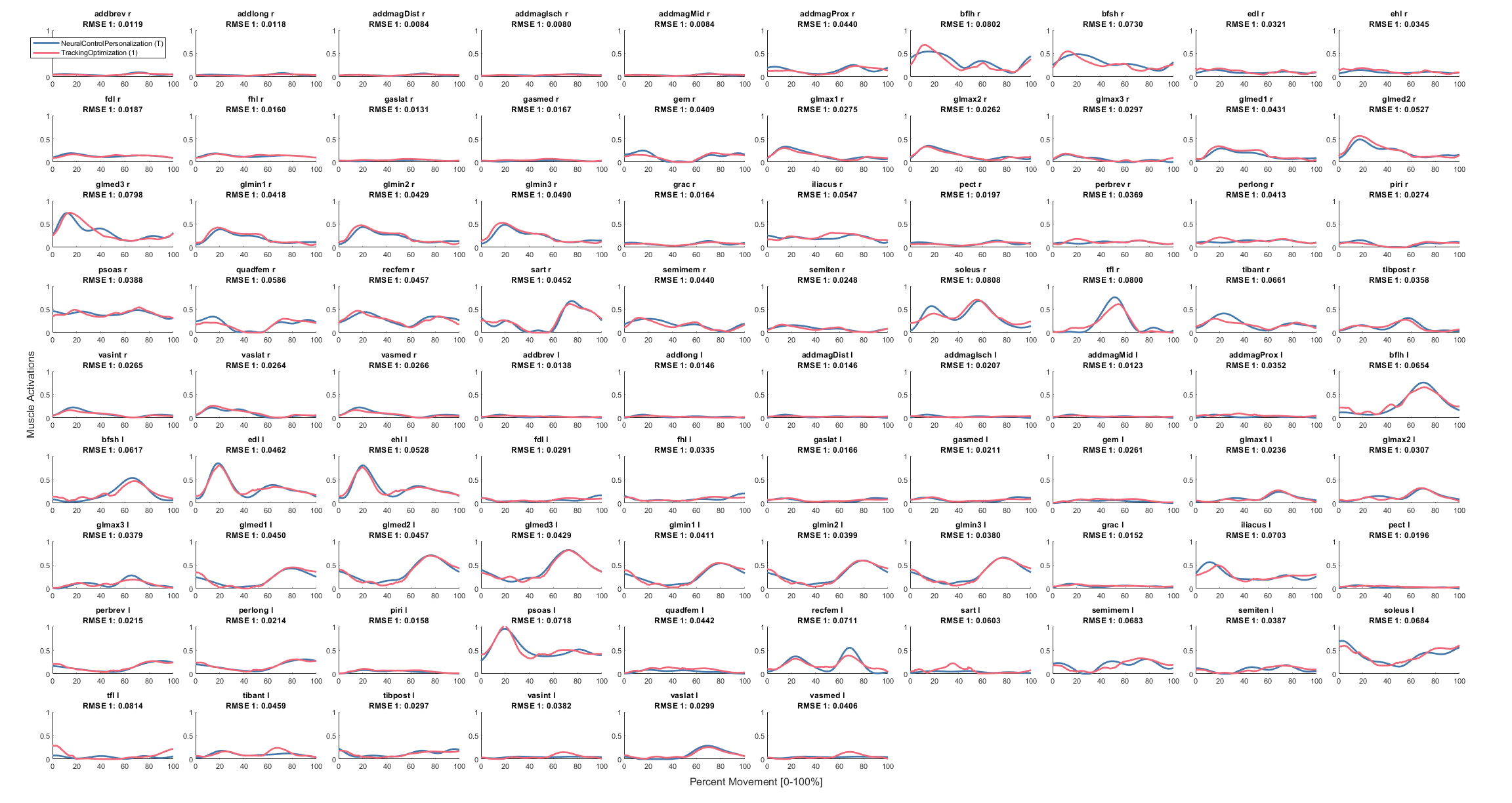

### foot1GroundReactions.png

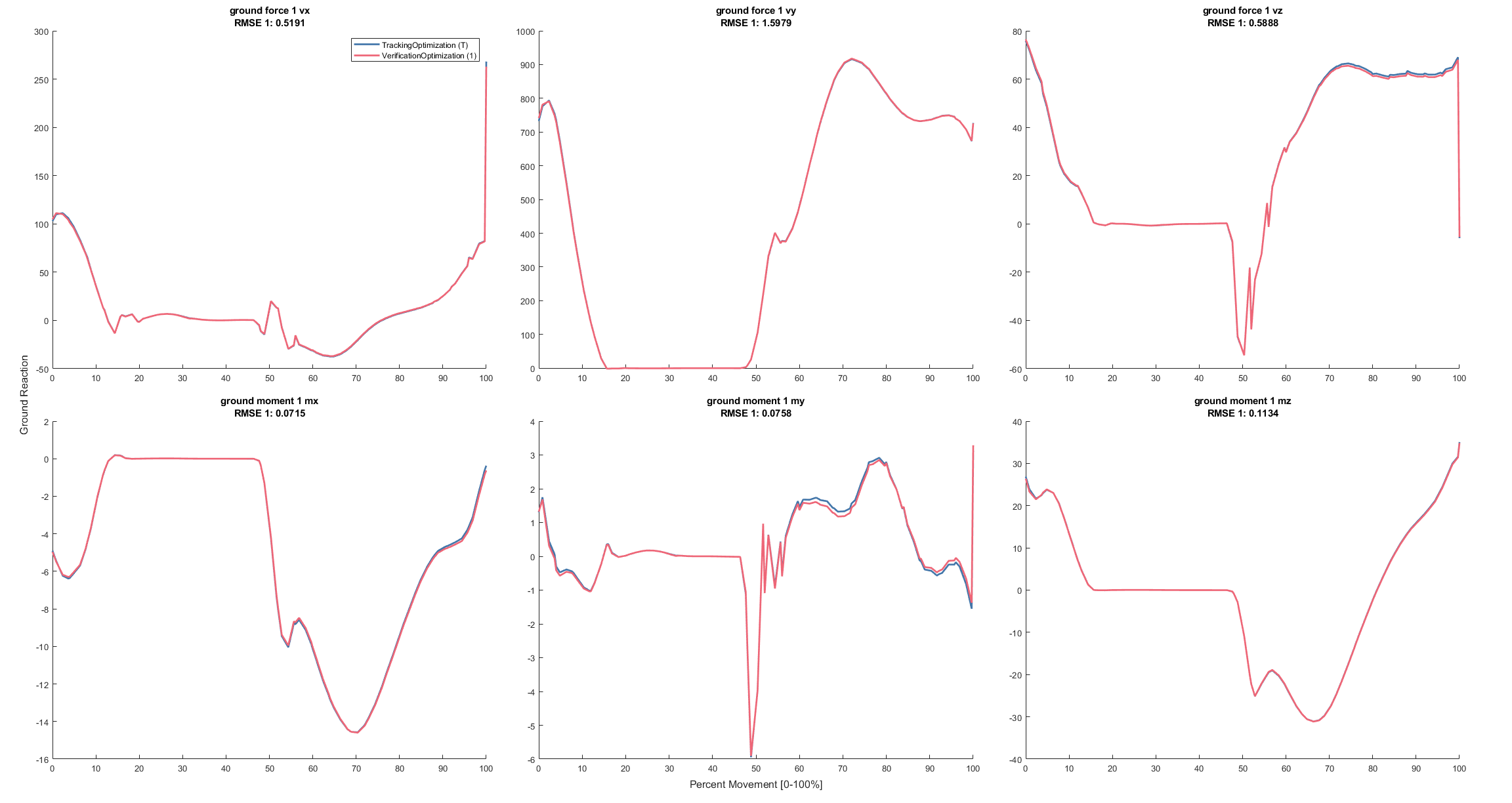

### foot1GroundReactions.png

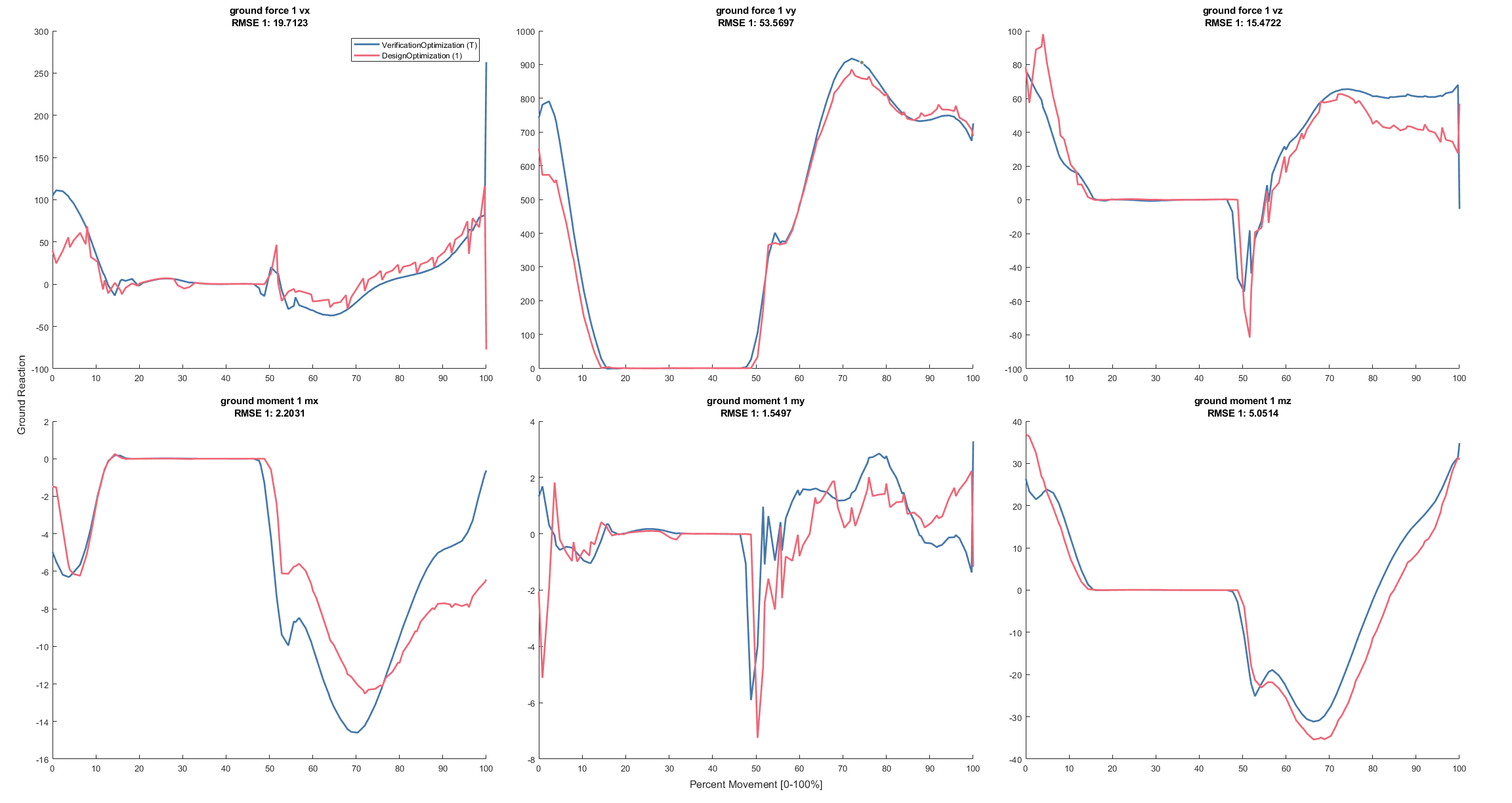

### foot1GroundReactions.png

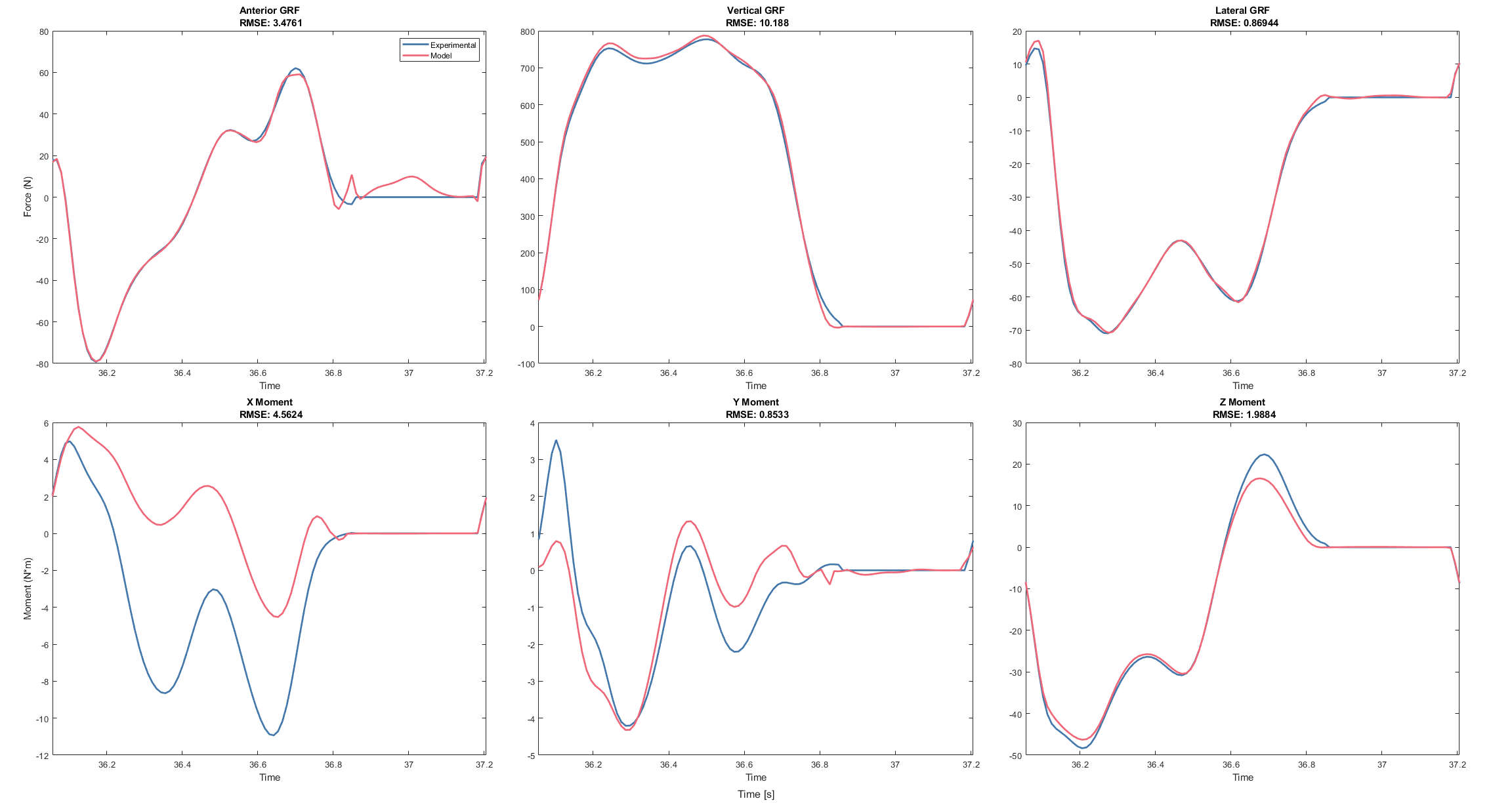

### foot1GroundReactions.png

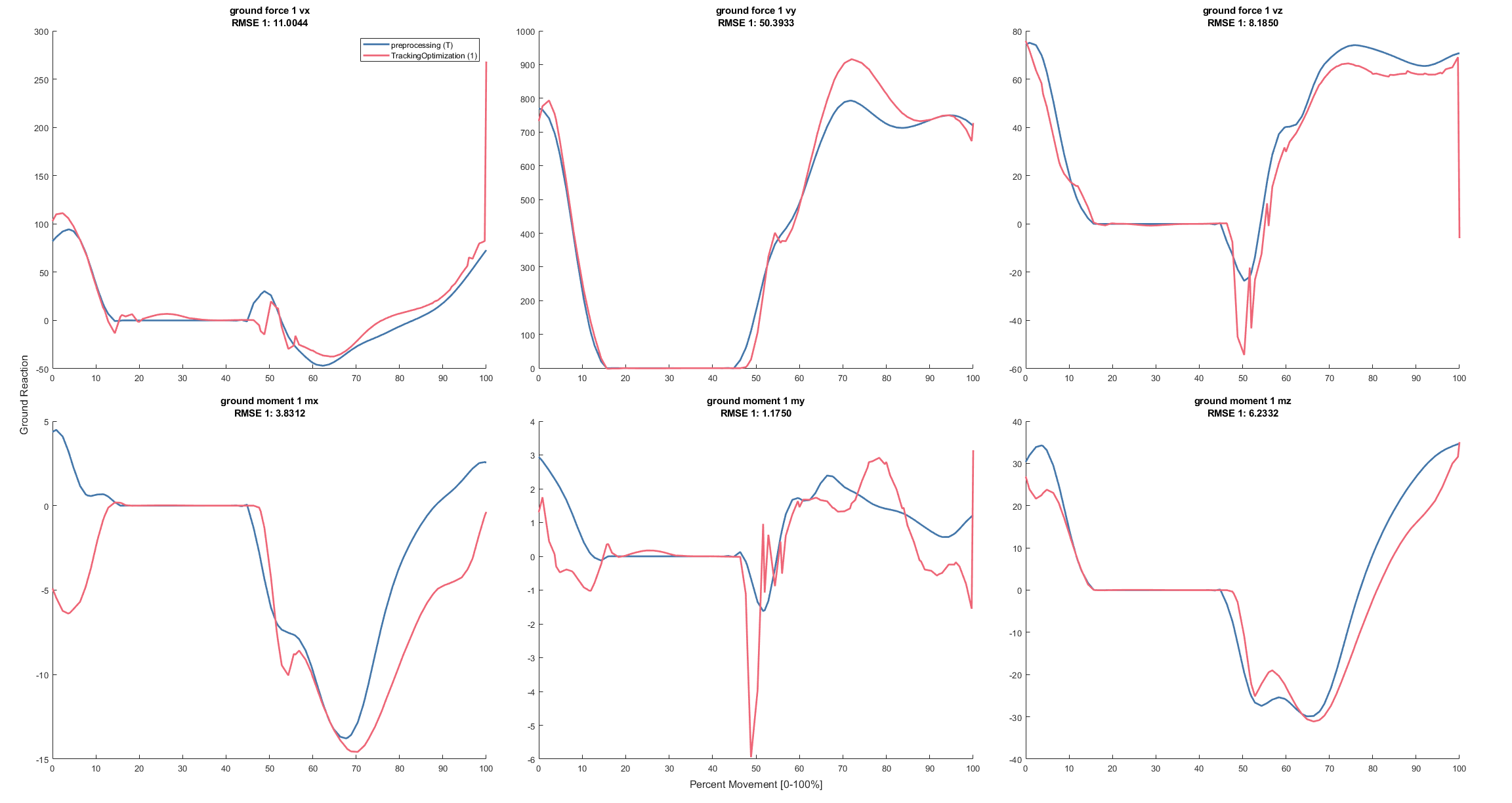

### foot1Kinematics.png

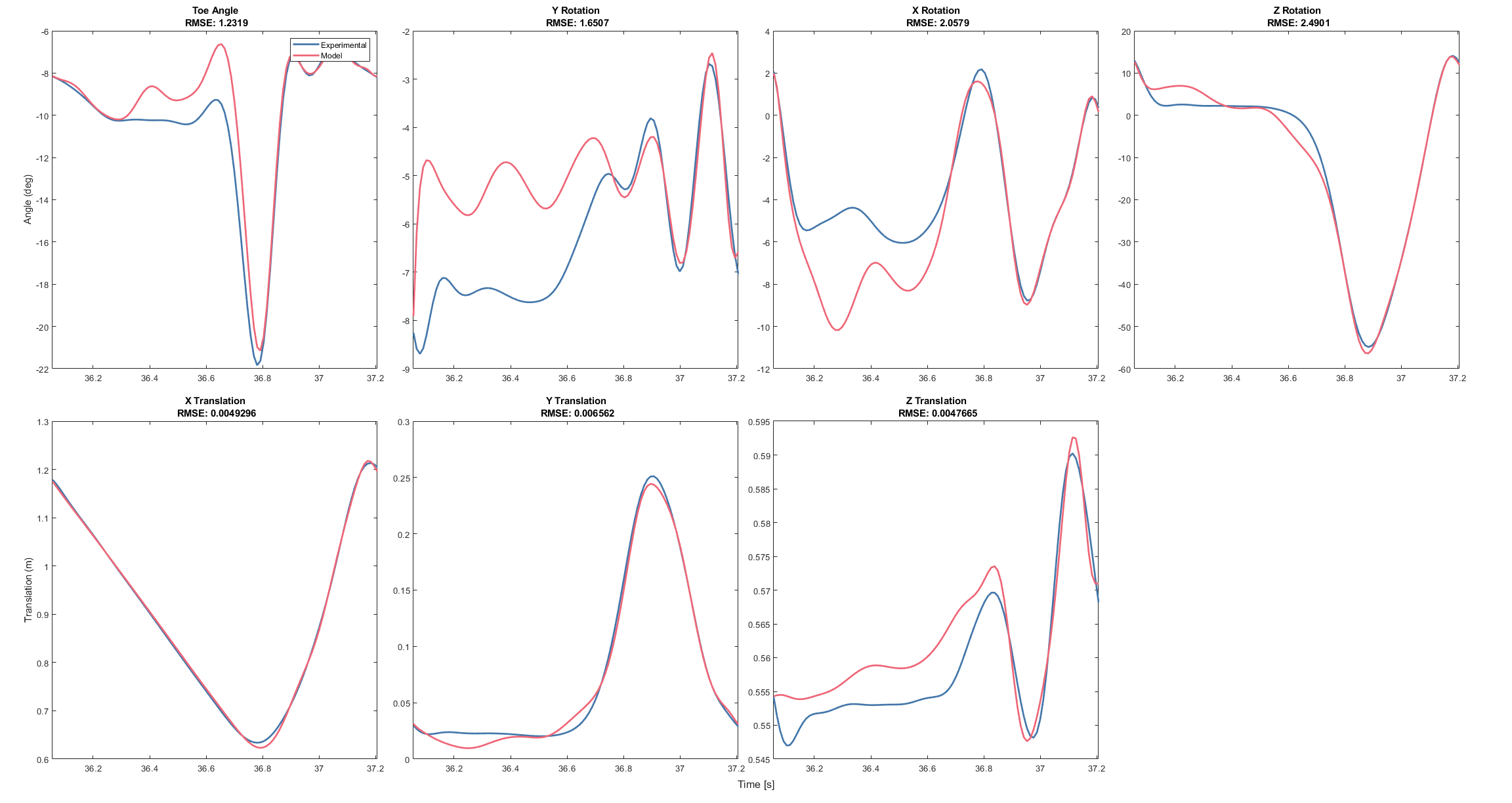

### foot2GroundReactions.png

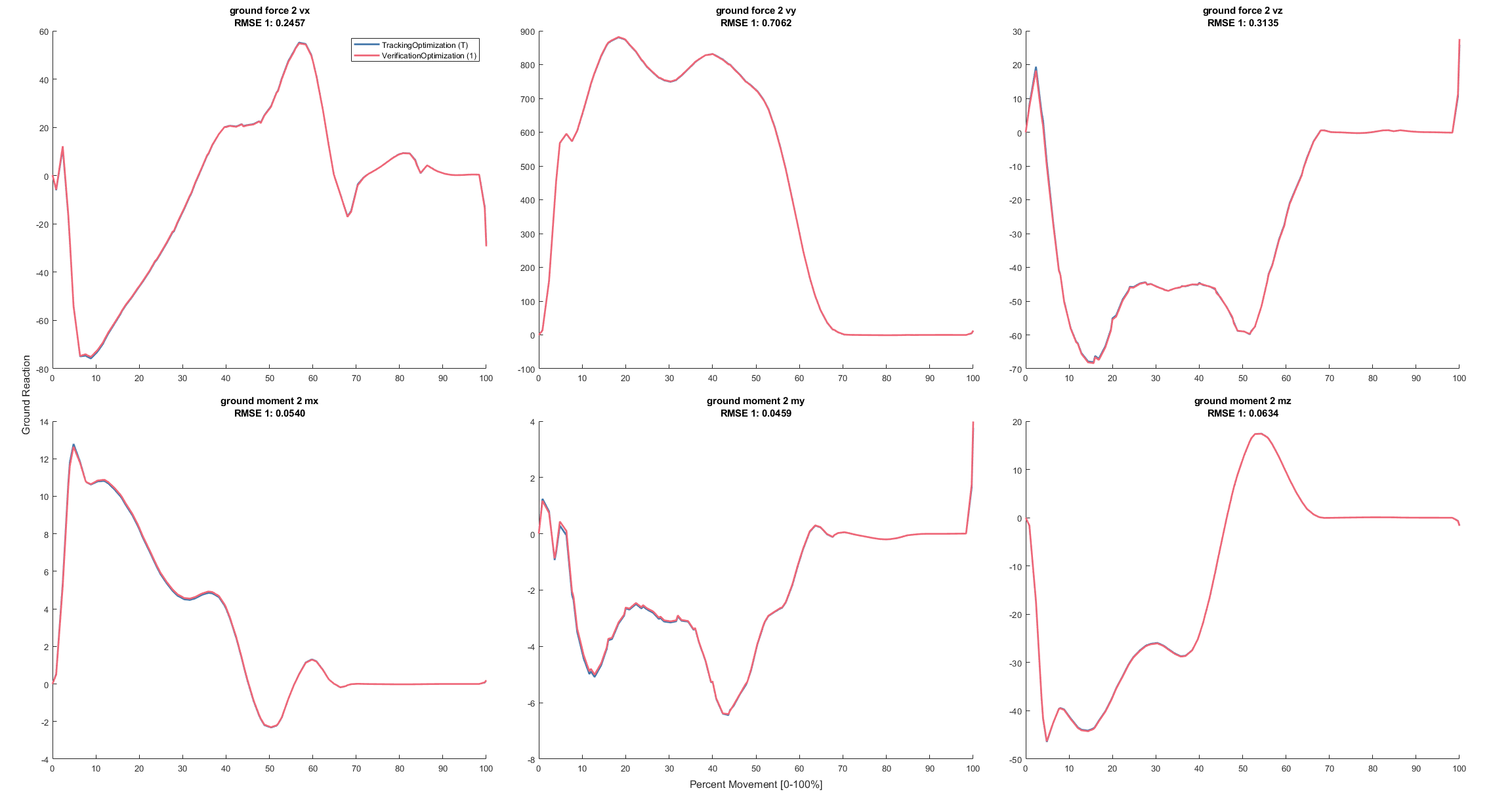

### foot2GroundReactions.png

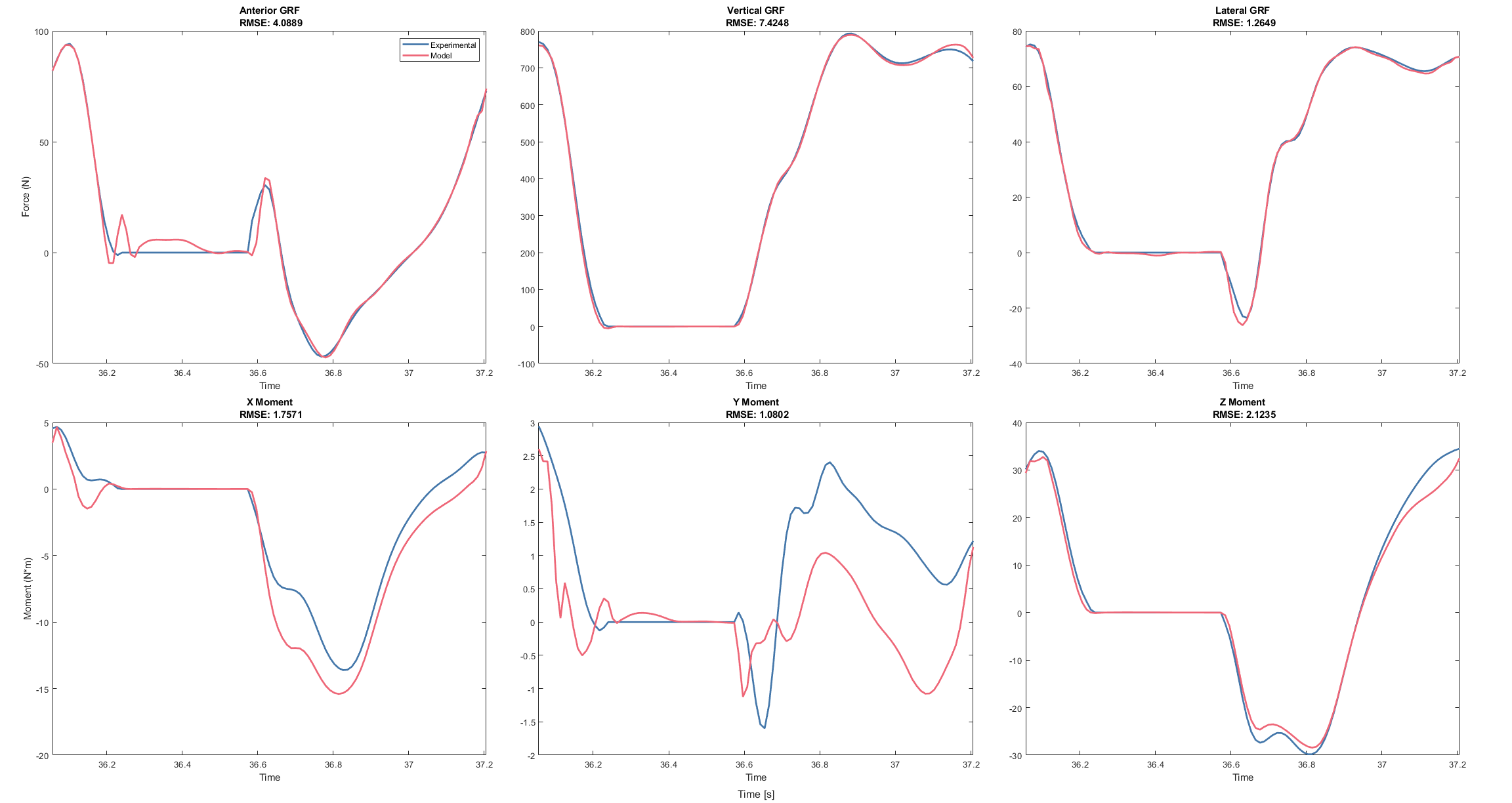

### foot2GroundReactions.png

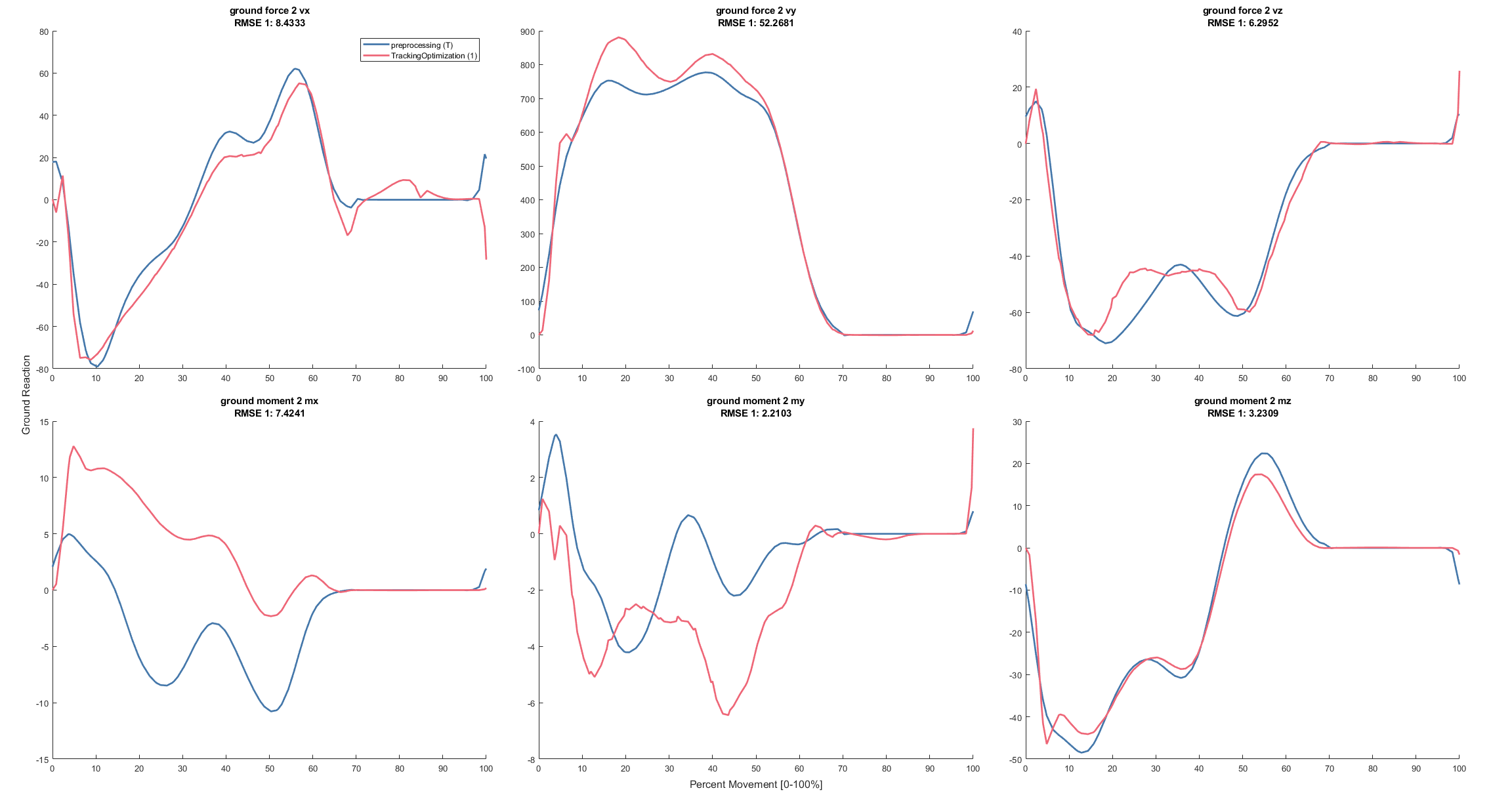

### foot2GroundReactionsa.png

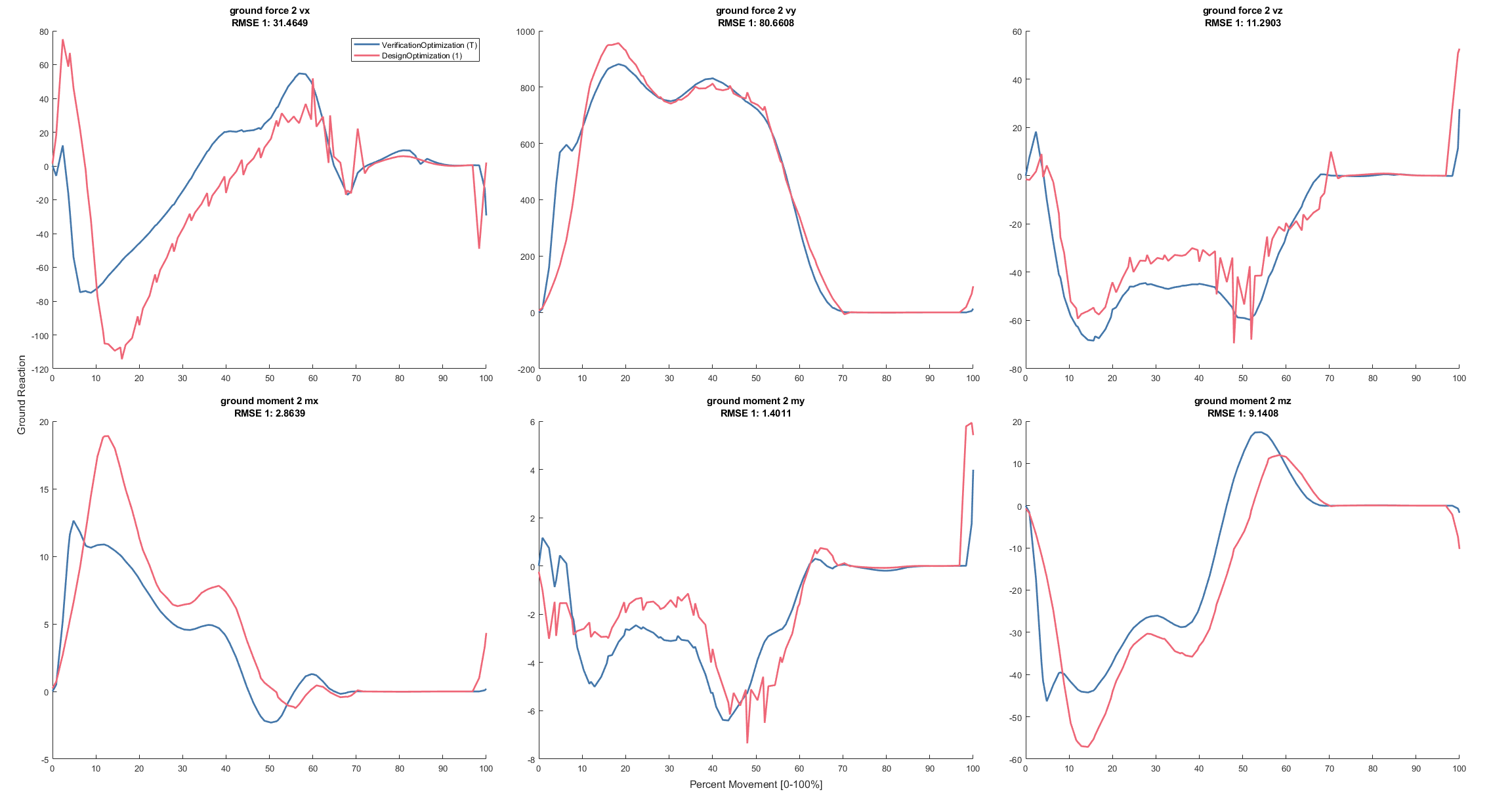

### foot2Kinematics.png

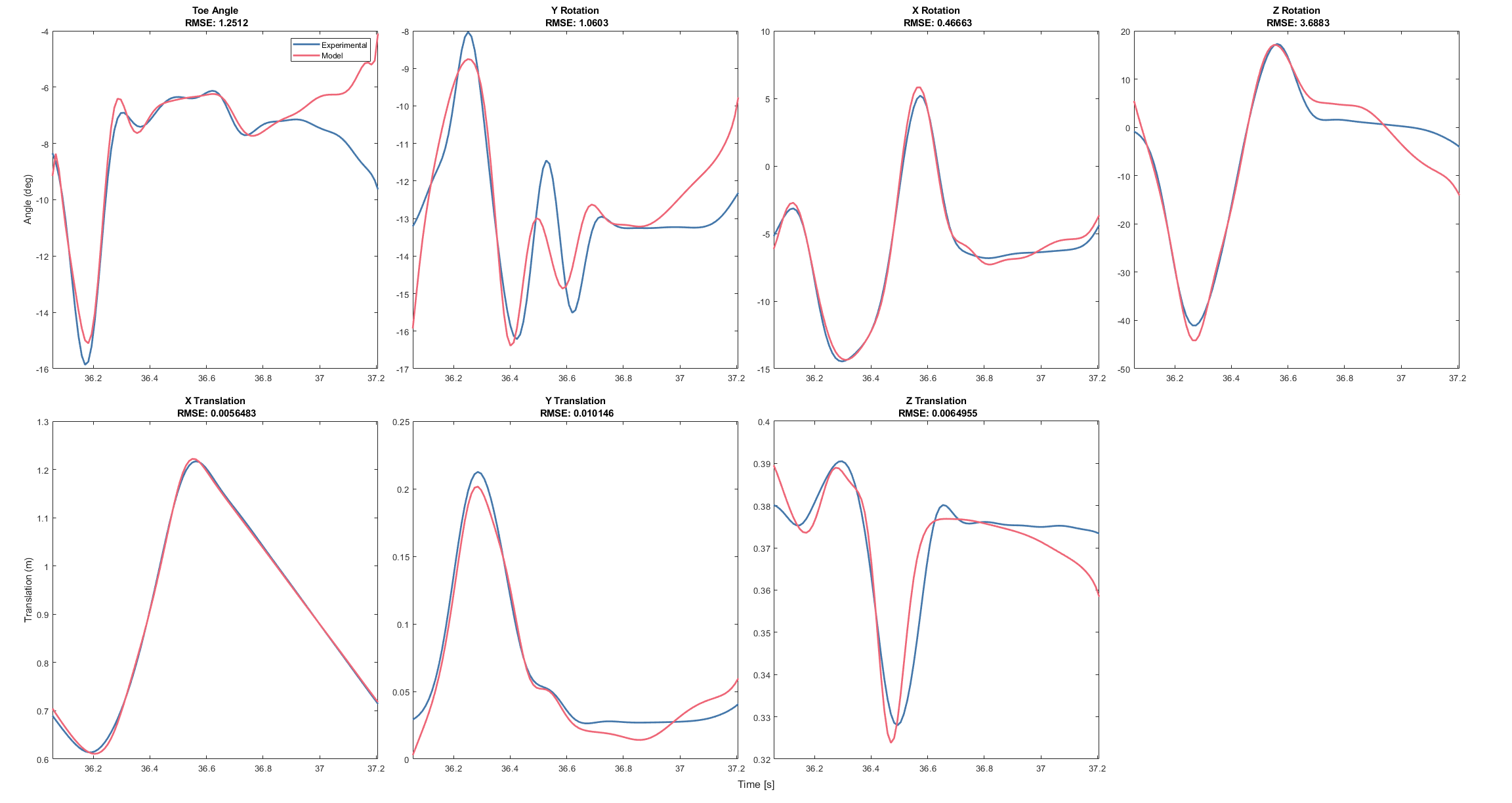

### jointAngles.png

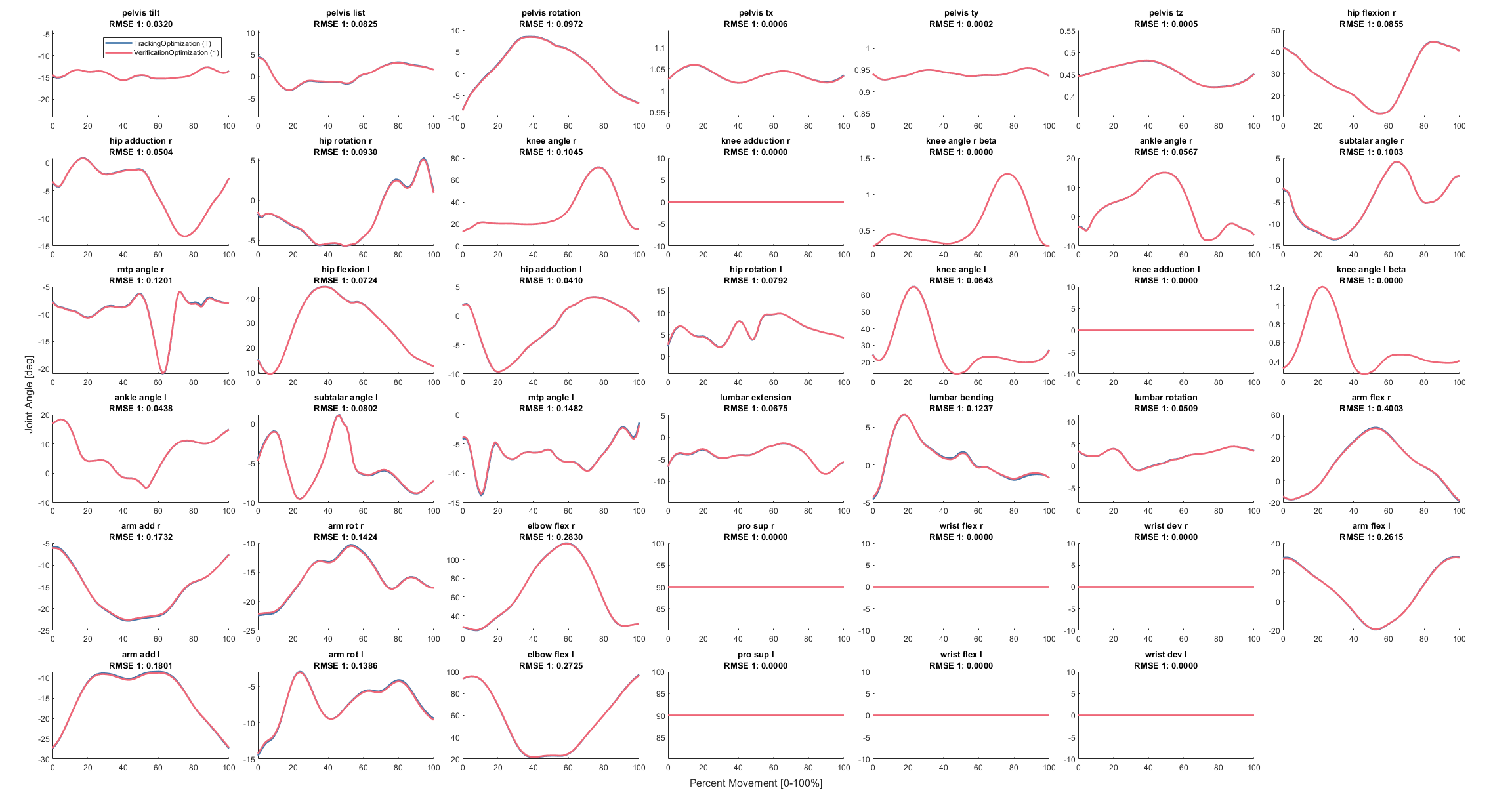

### jointAngles.png

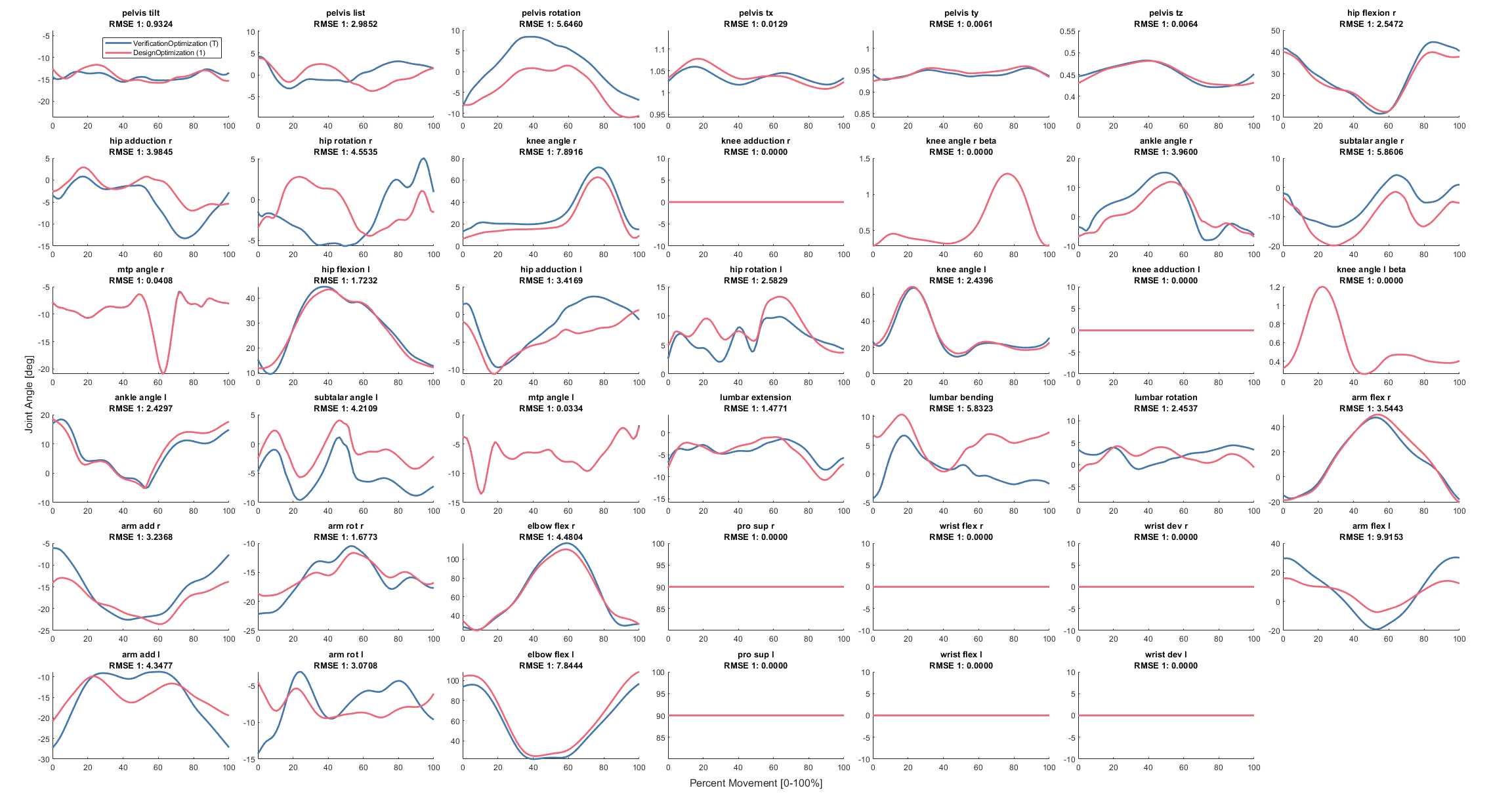

### jointAngles.png

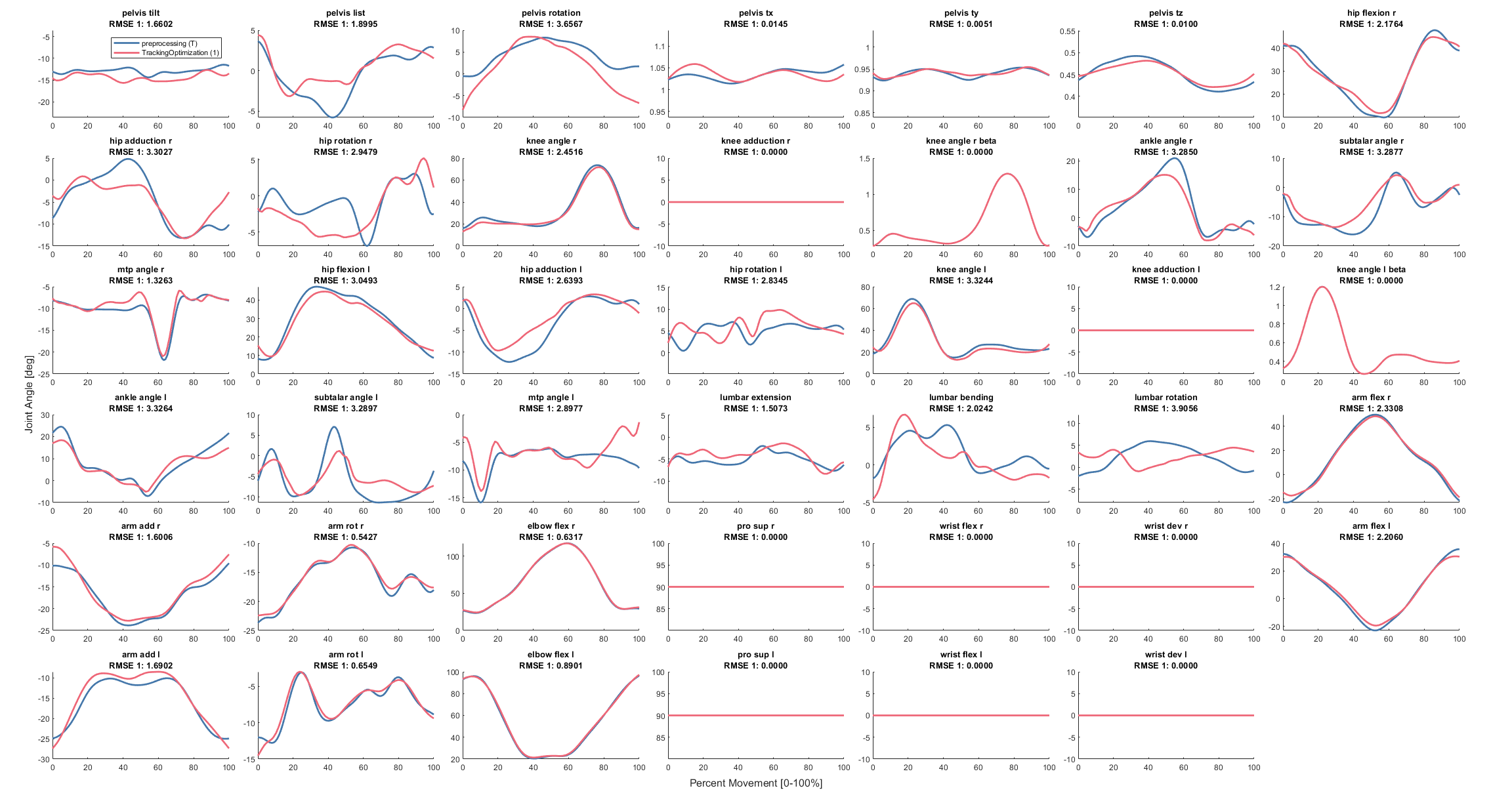

### jointLoads.png

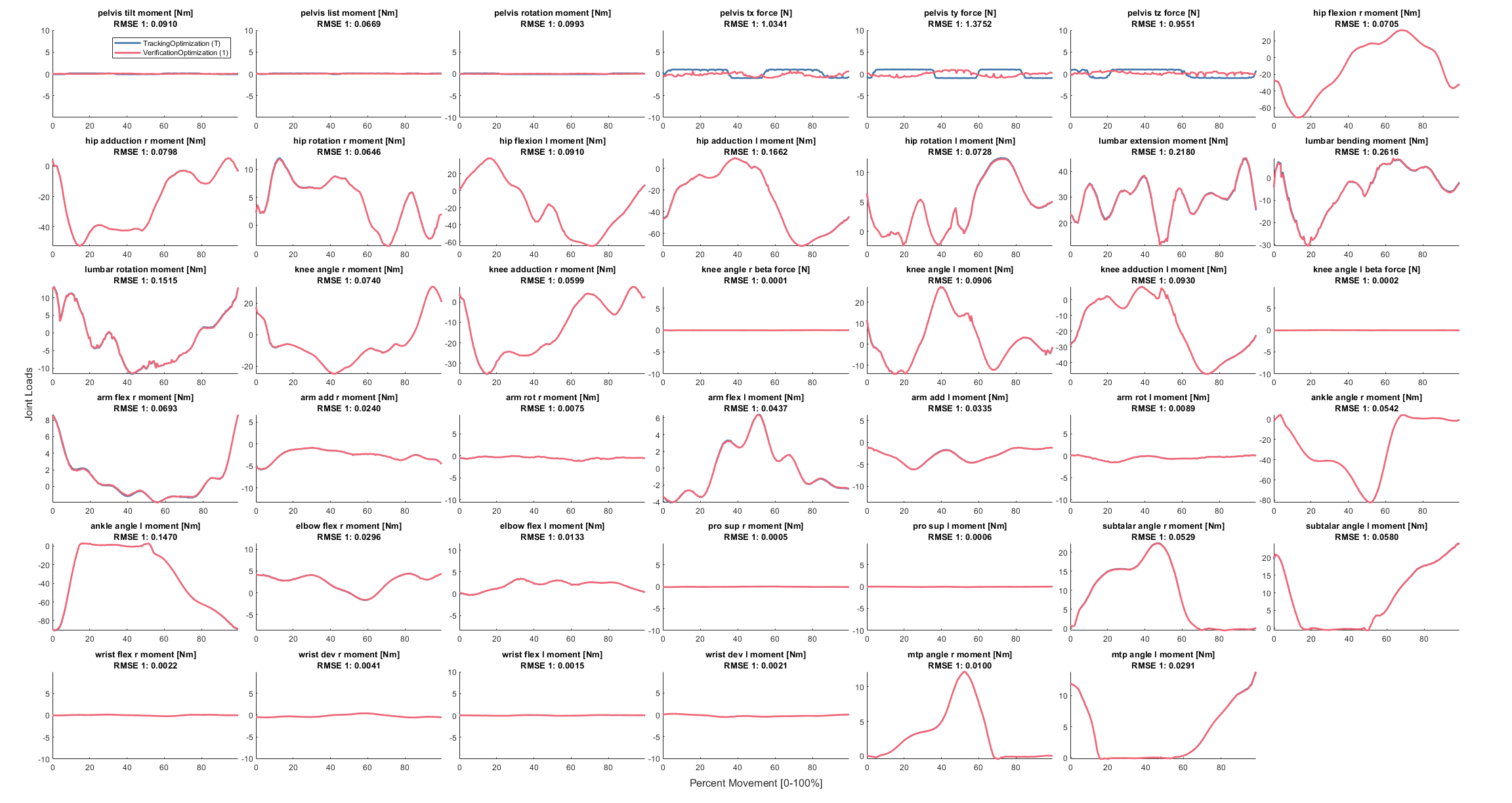

### jointLoads.png

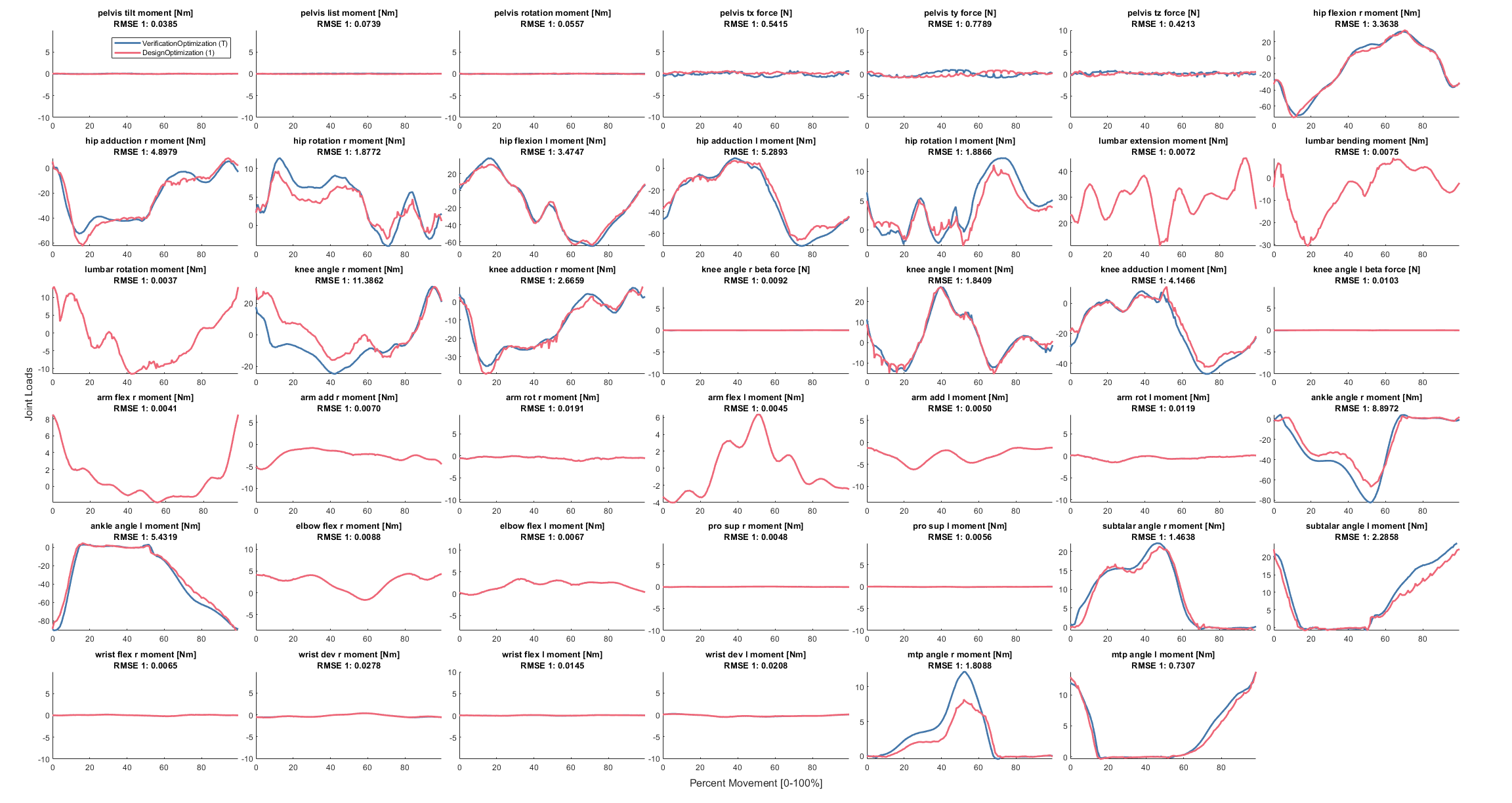

### jointLoads.png

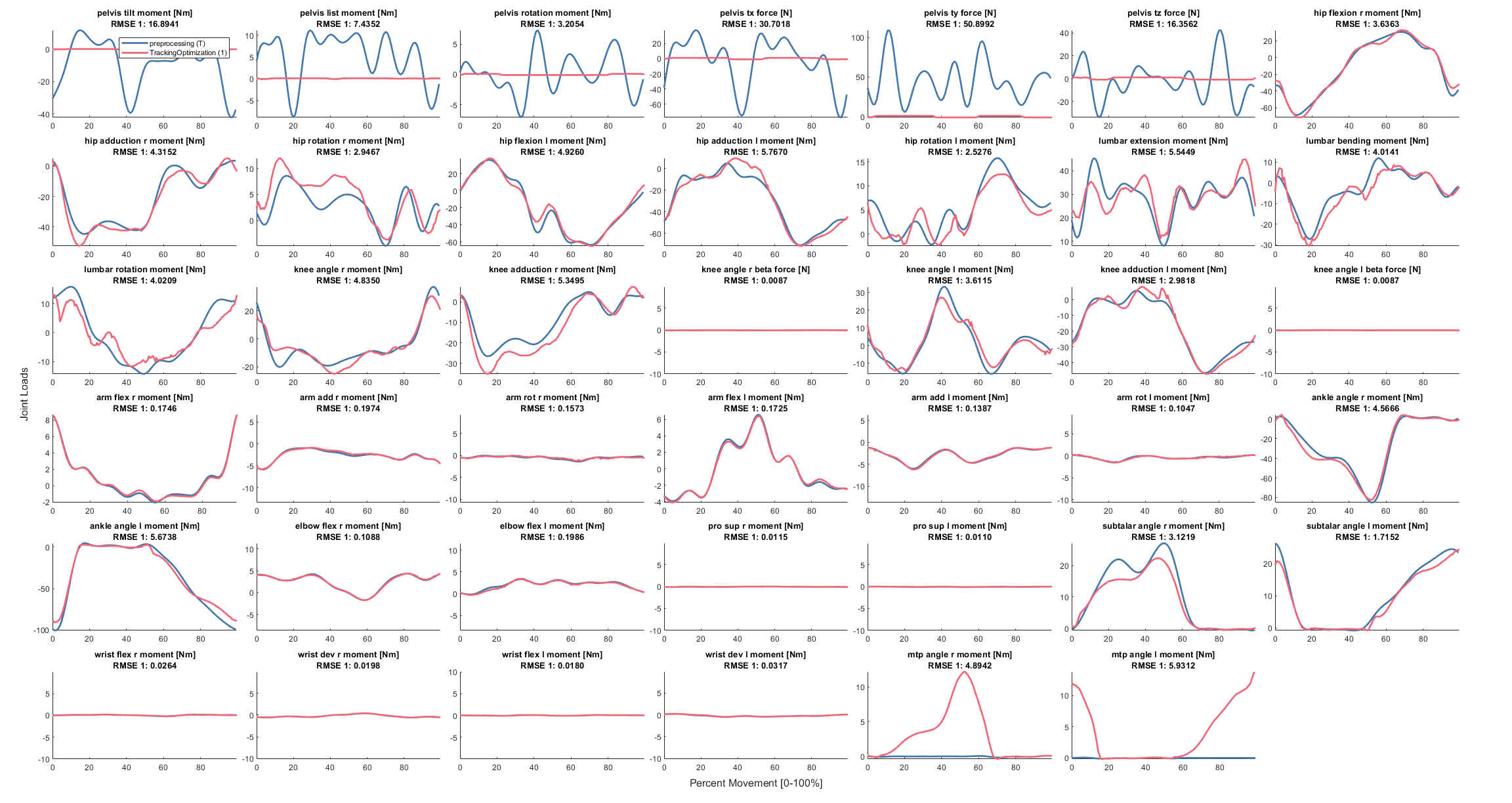

### jointMoments.png

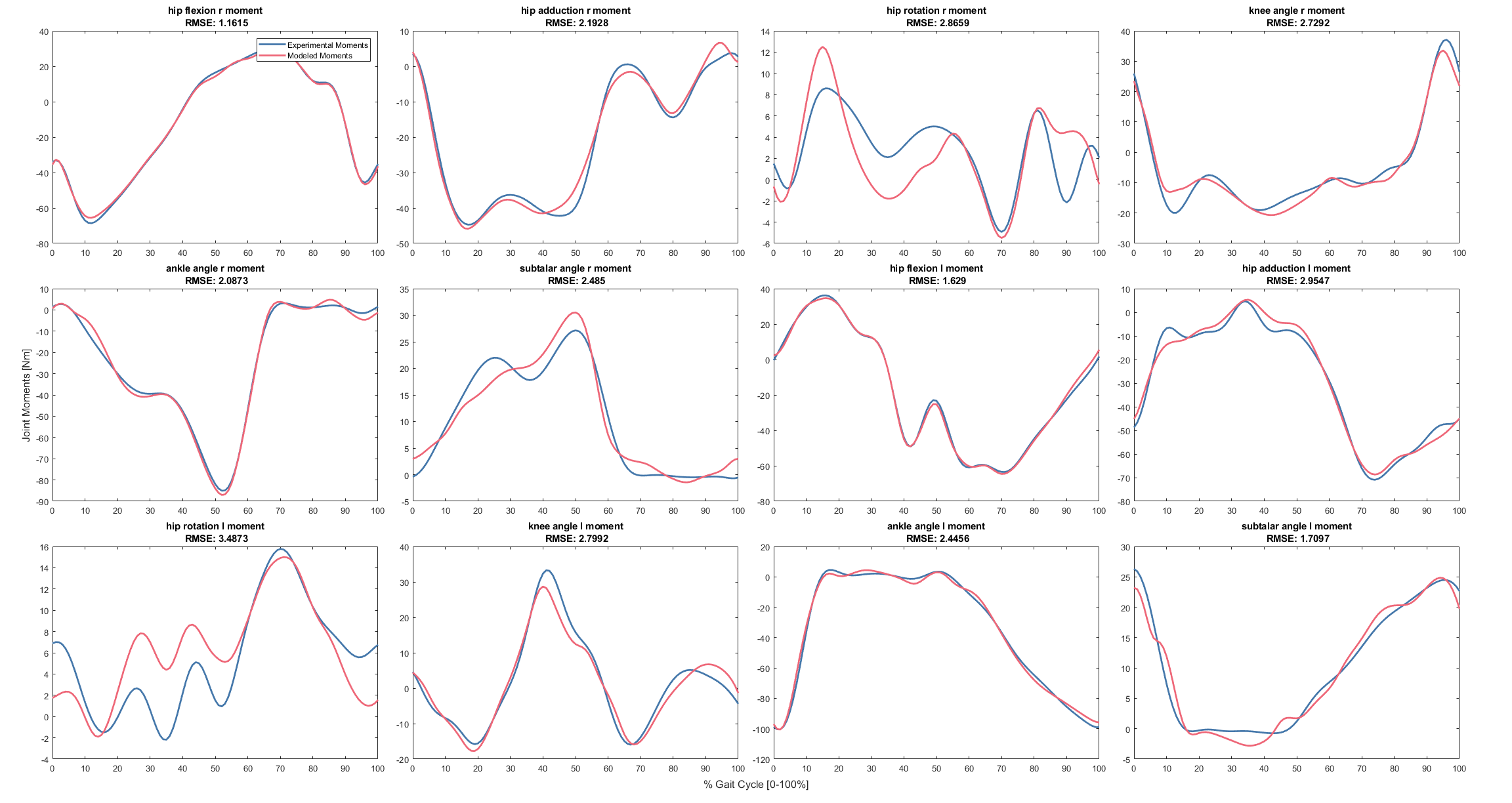

### jointVelocities.png

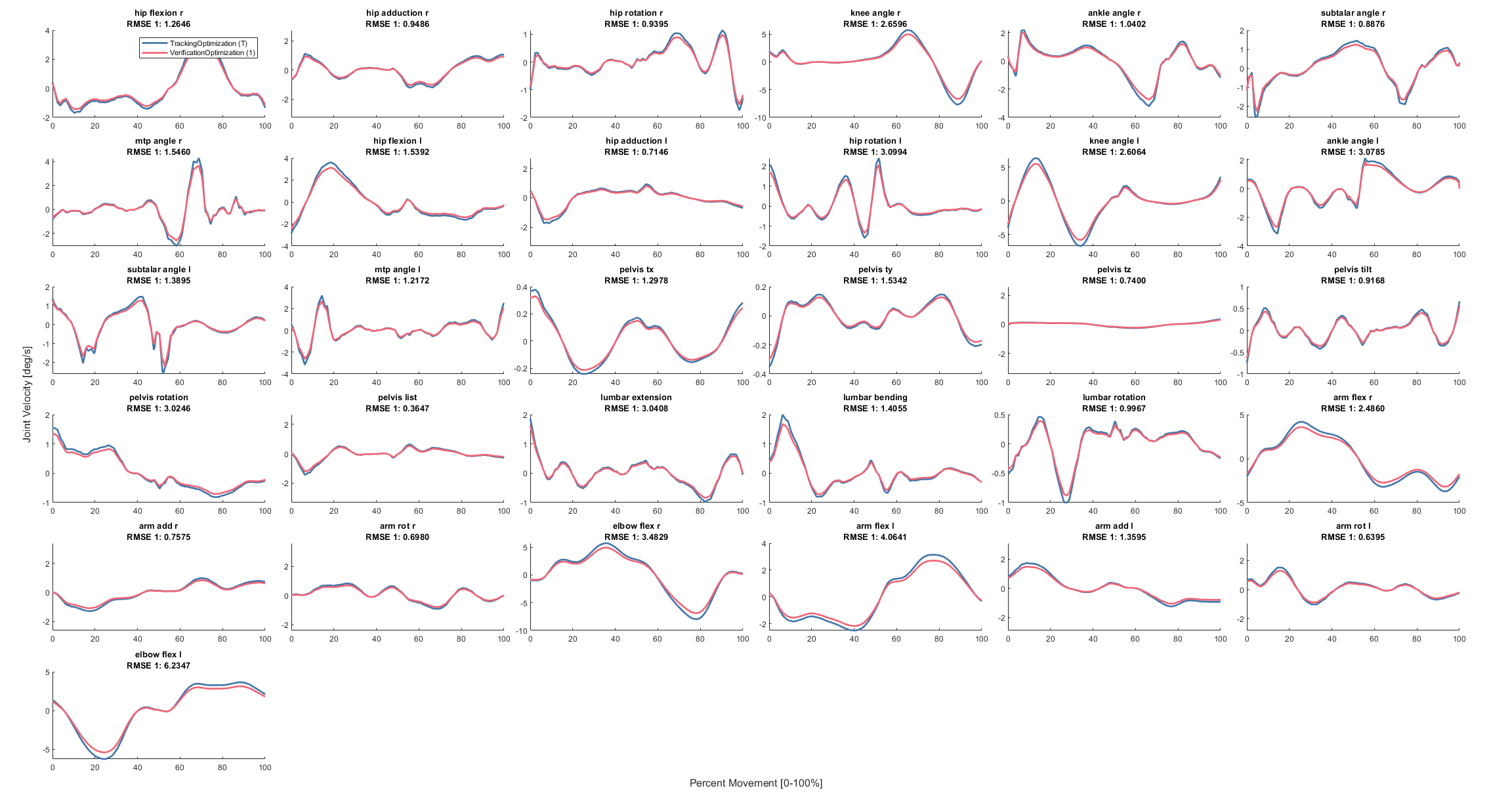

### jointVelocities.png

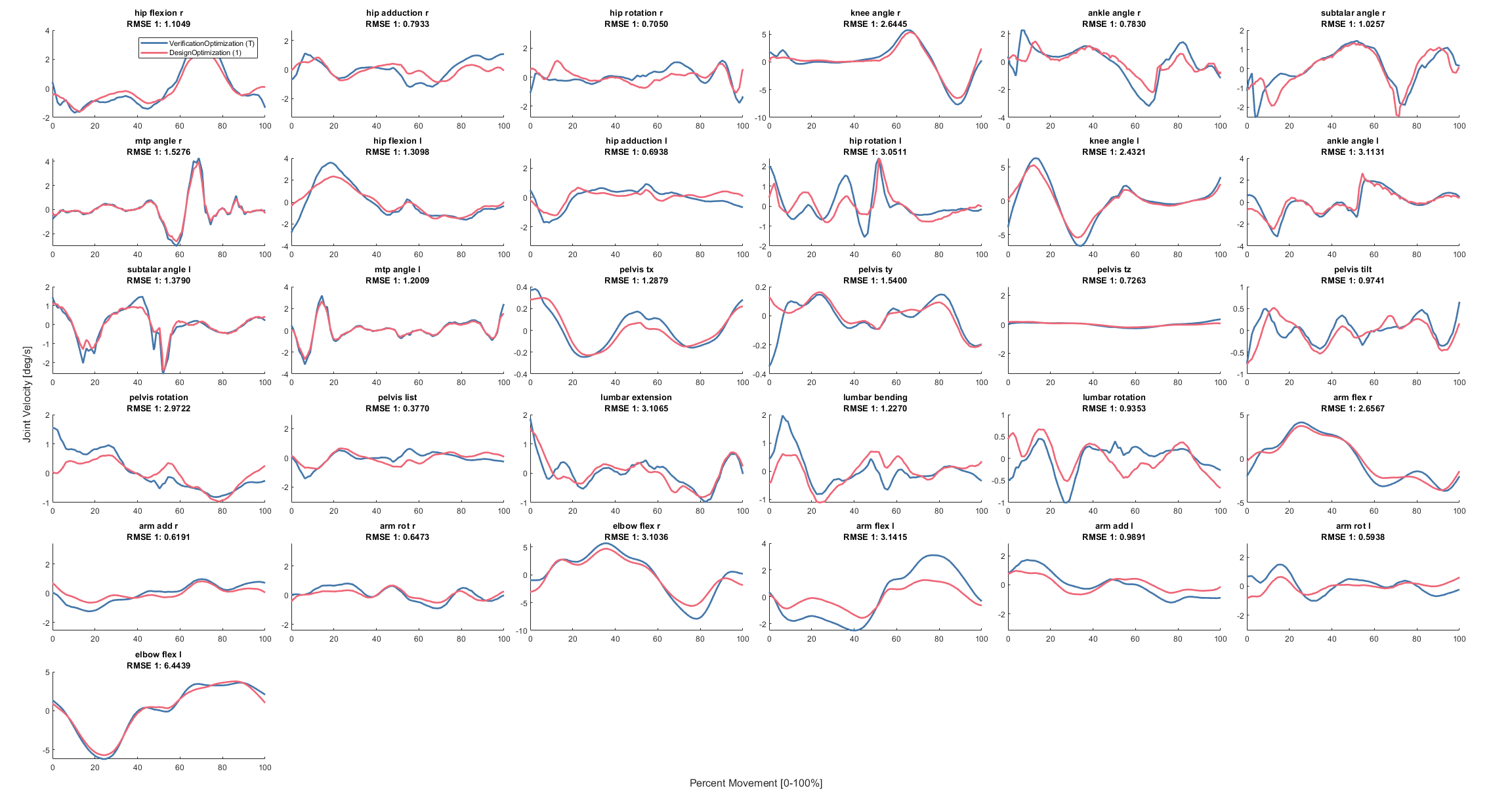

### jointVelocities.png

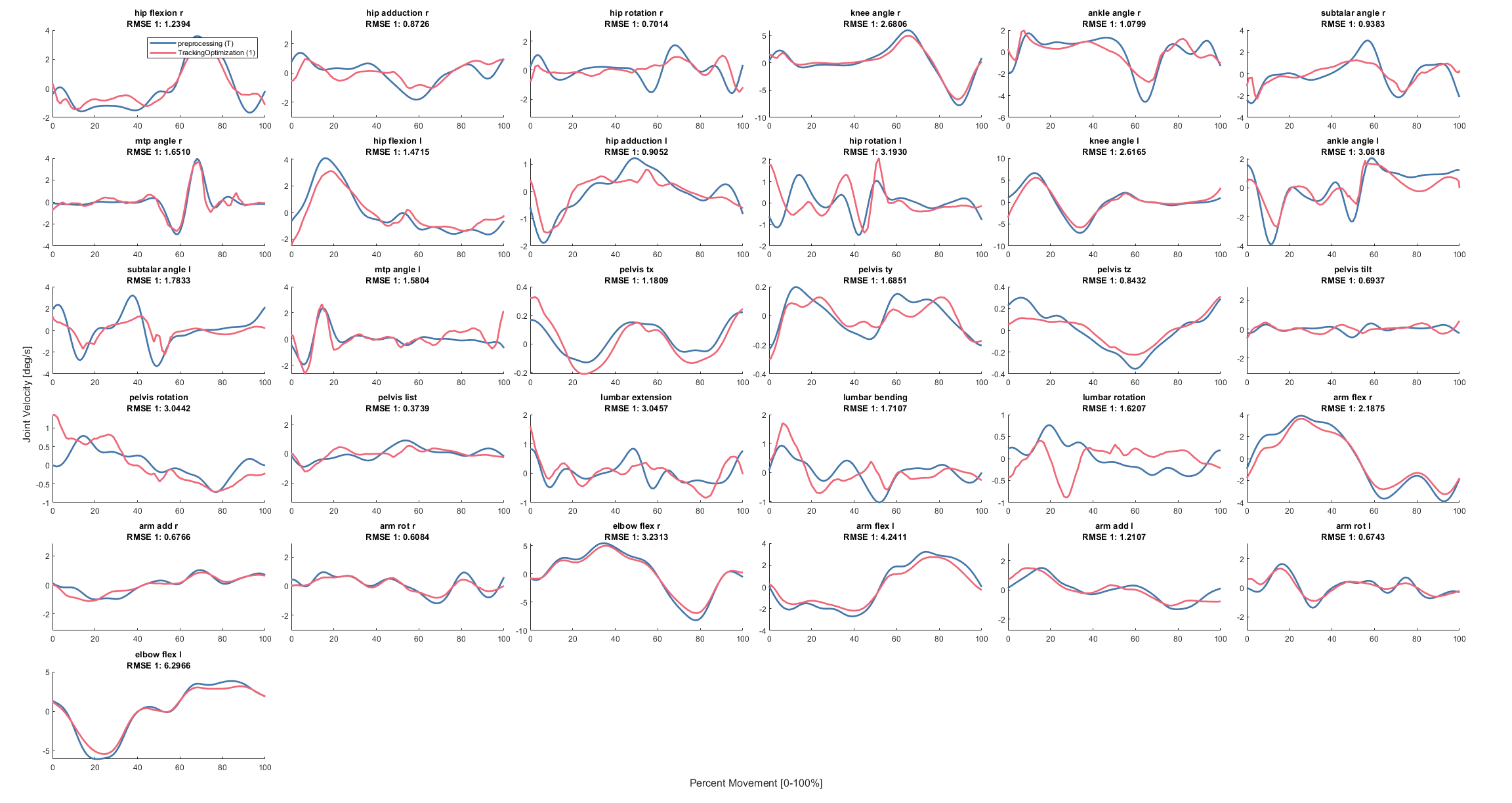

### leftActivation.png

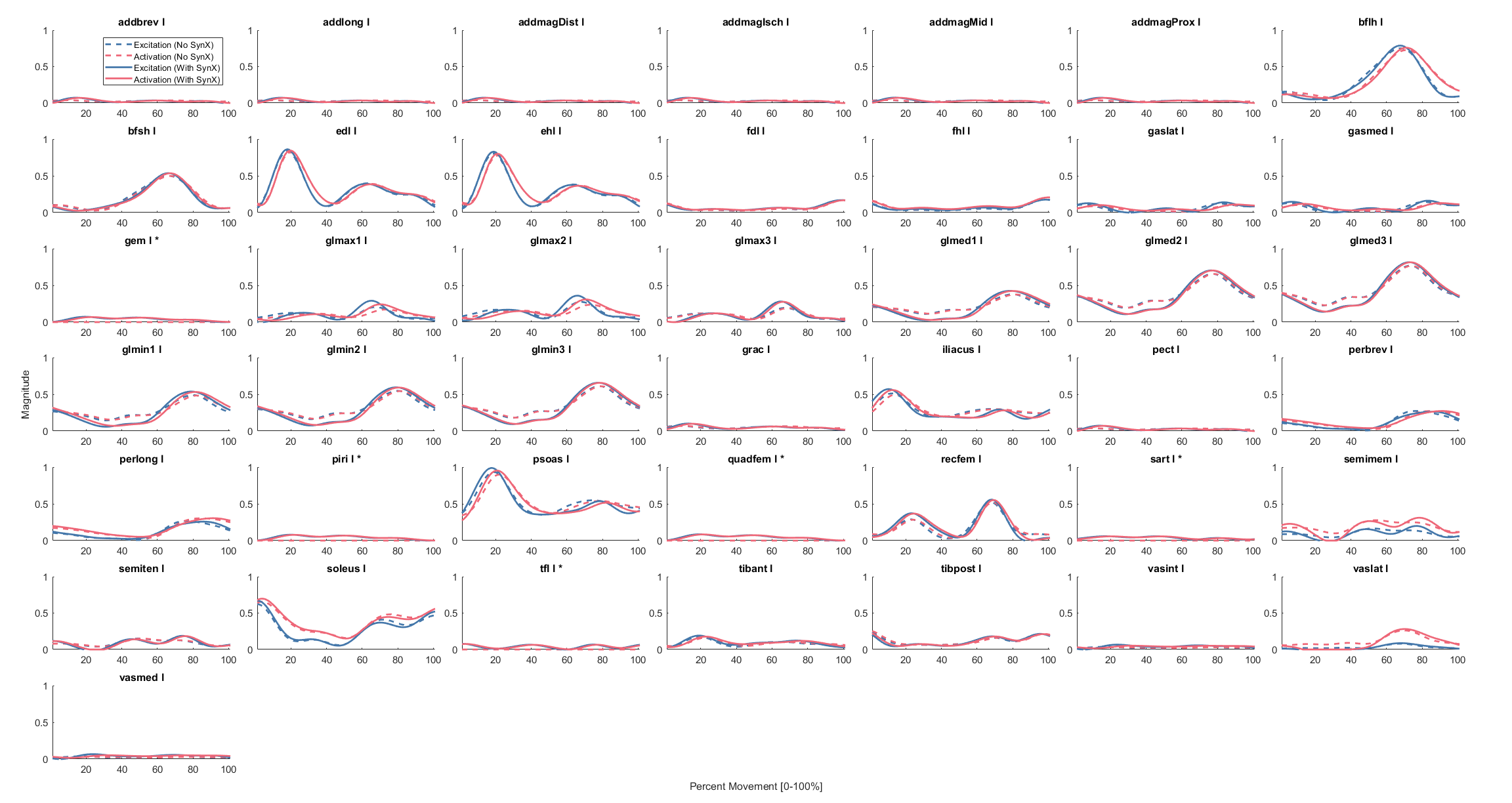

### leftFiberLength.png

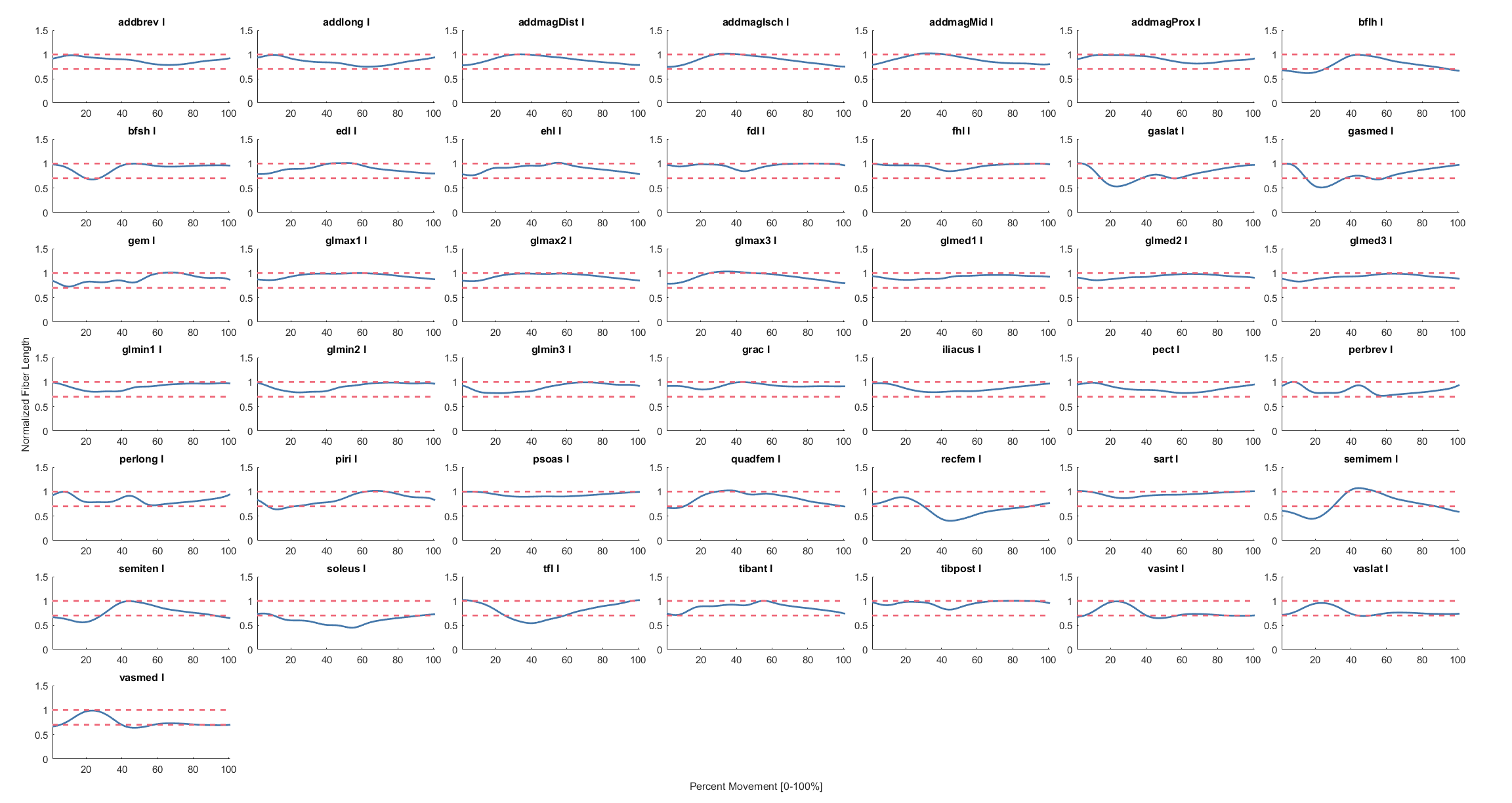

### leftMoment.png

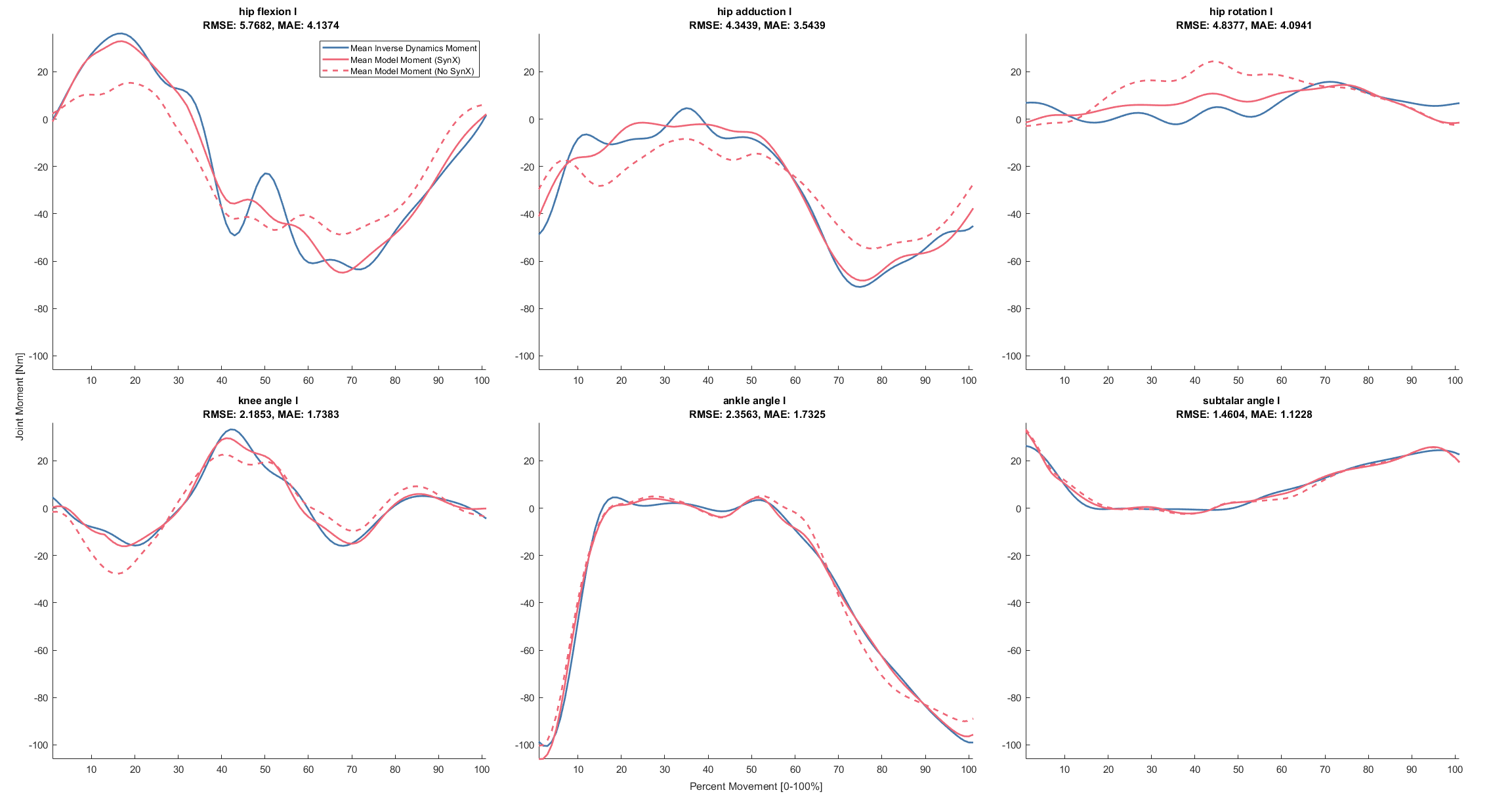

### leftParameters.png

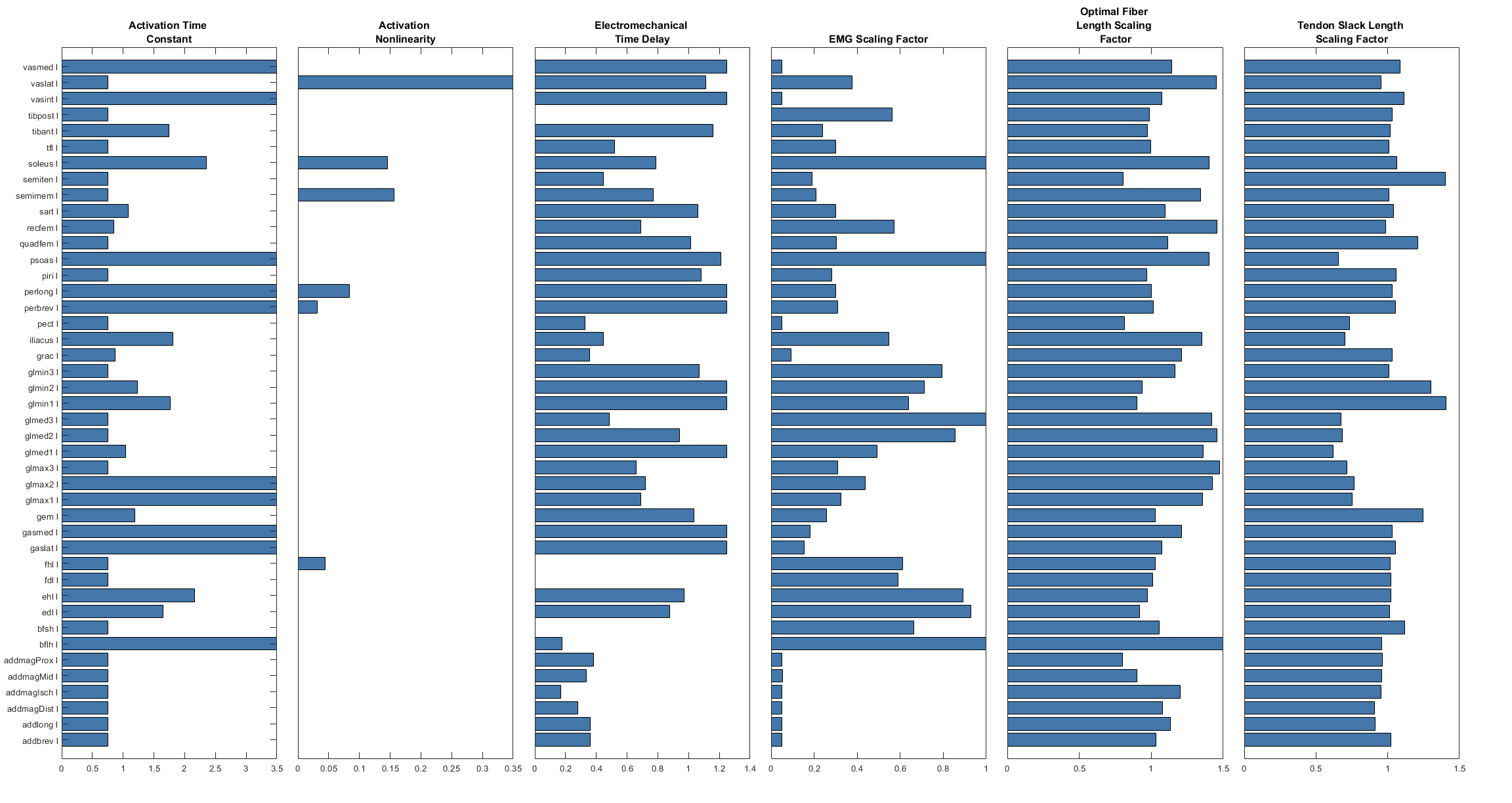

### leftPassiveForce.png

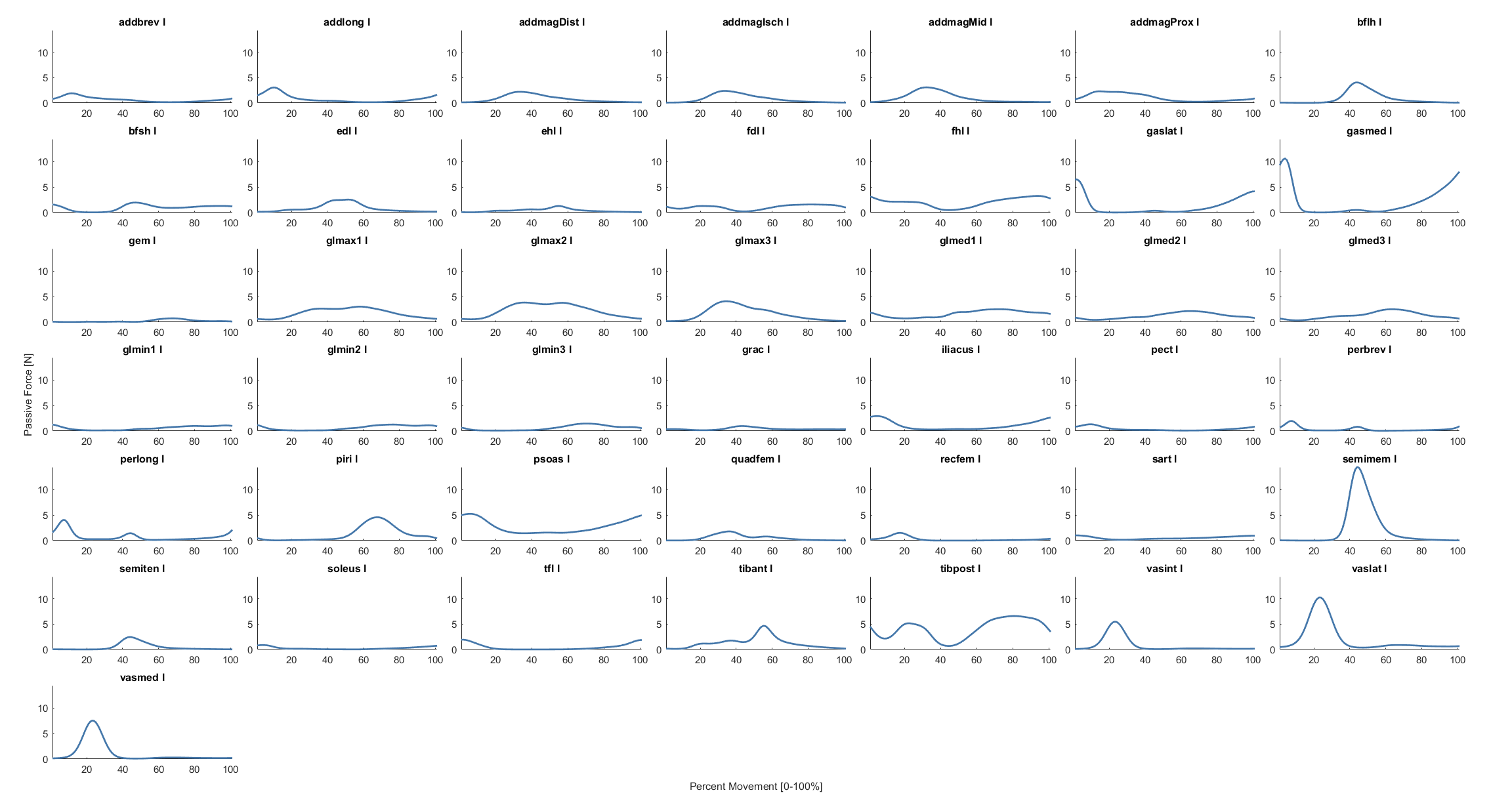

### leftPassiveMoment.png

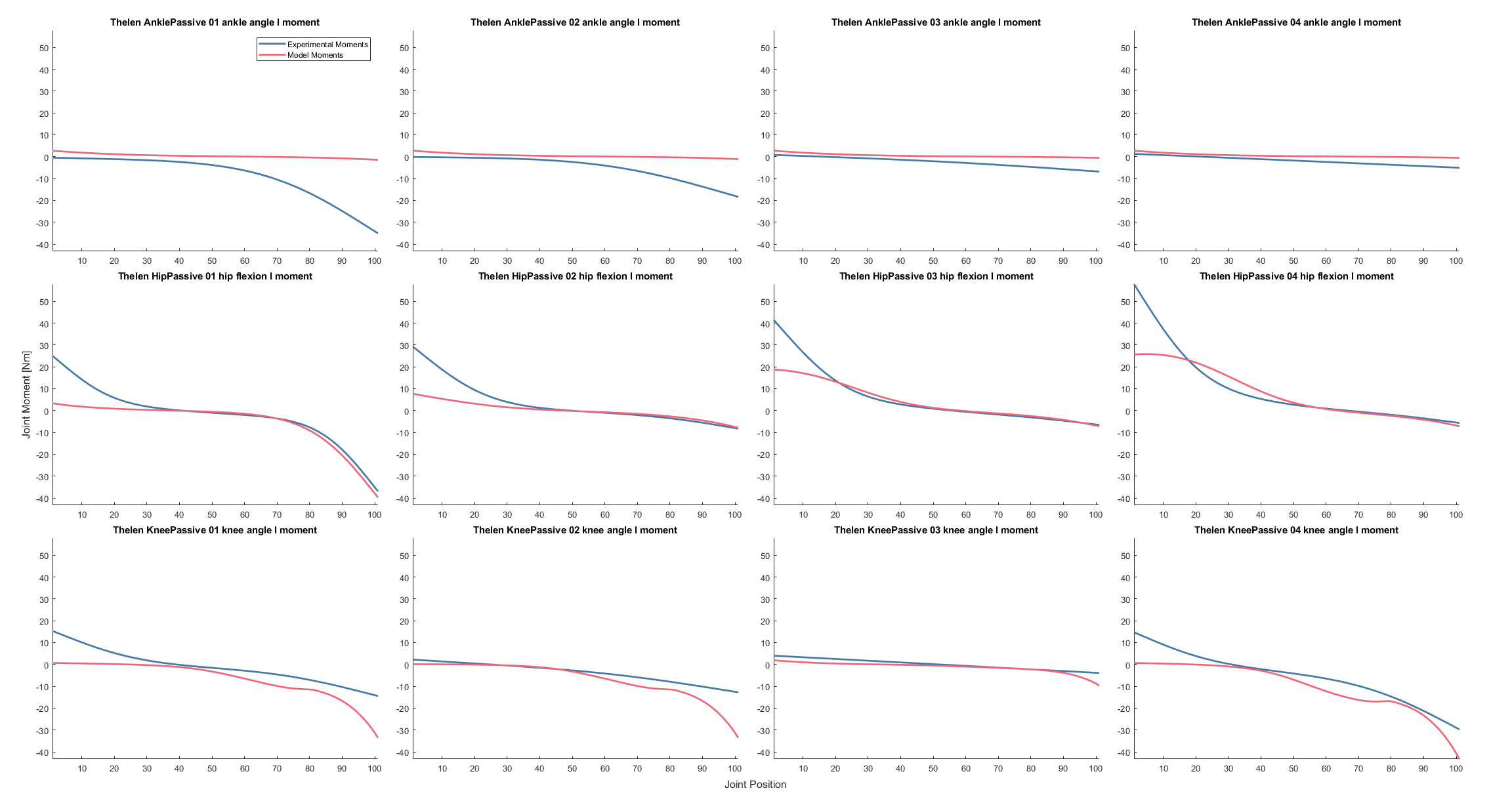

### markerErrors.png

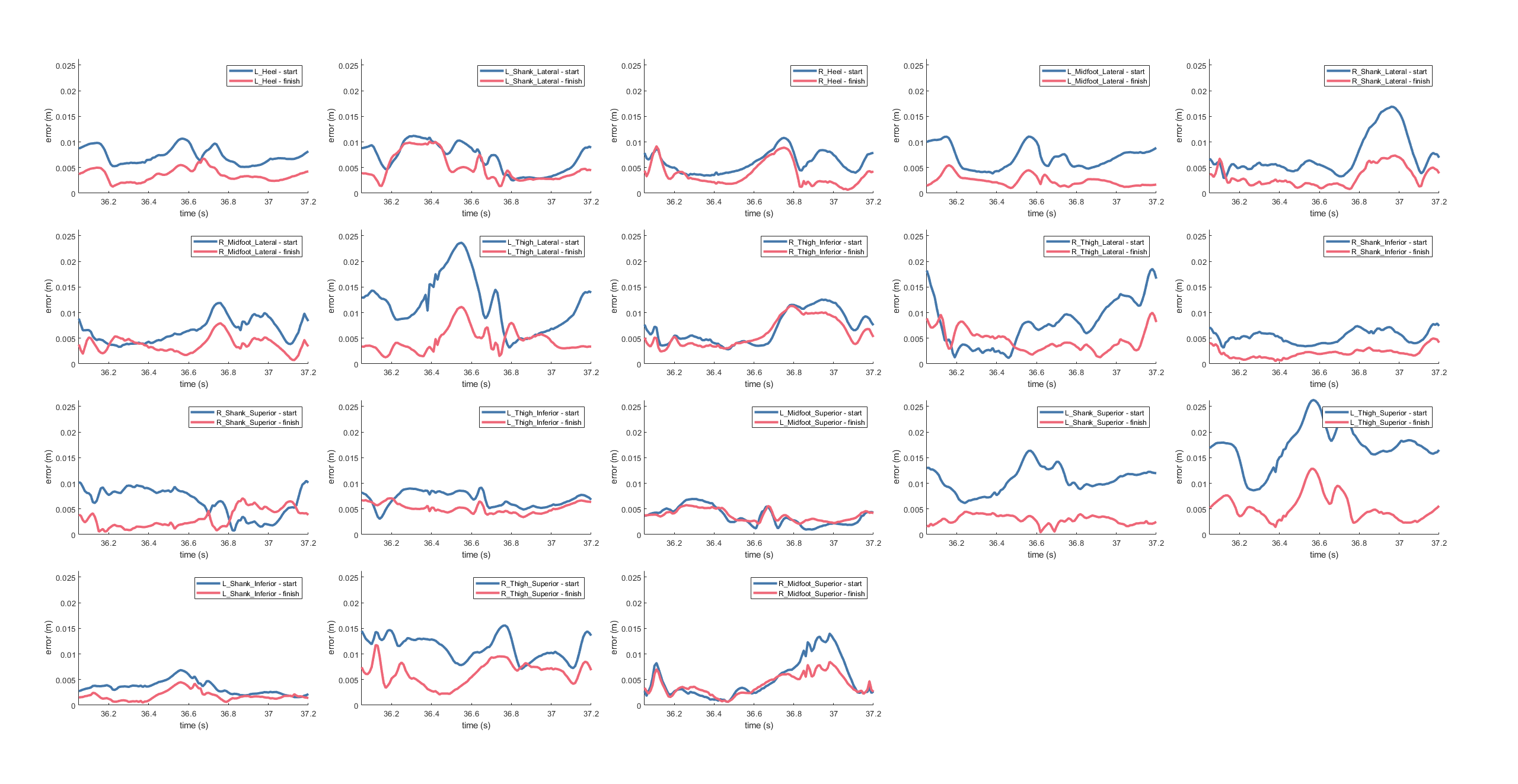

### muscleActivations.png

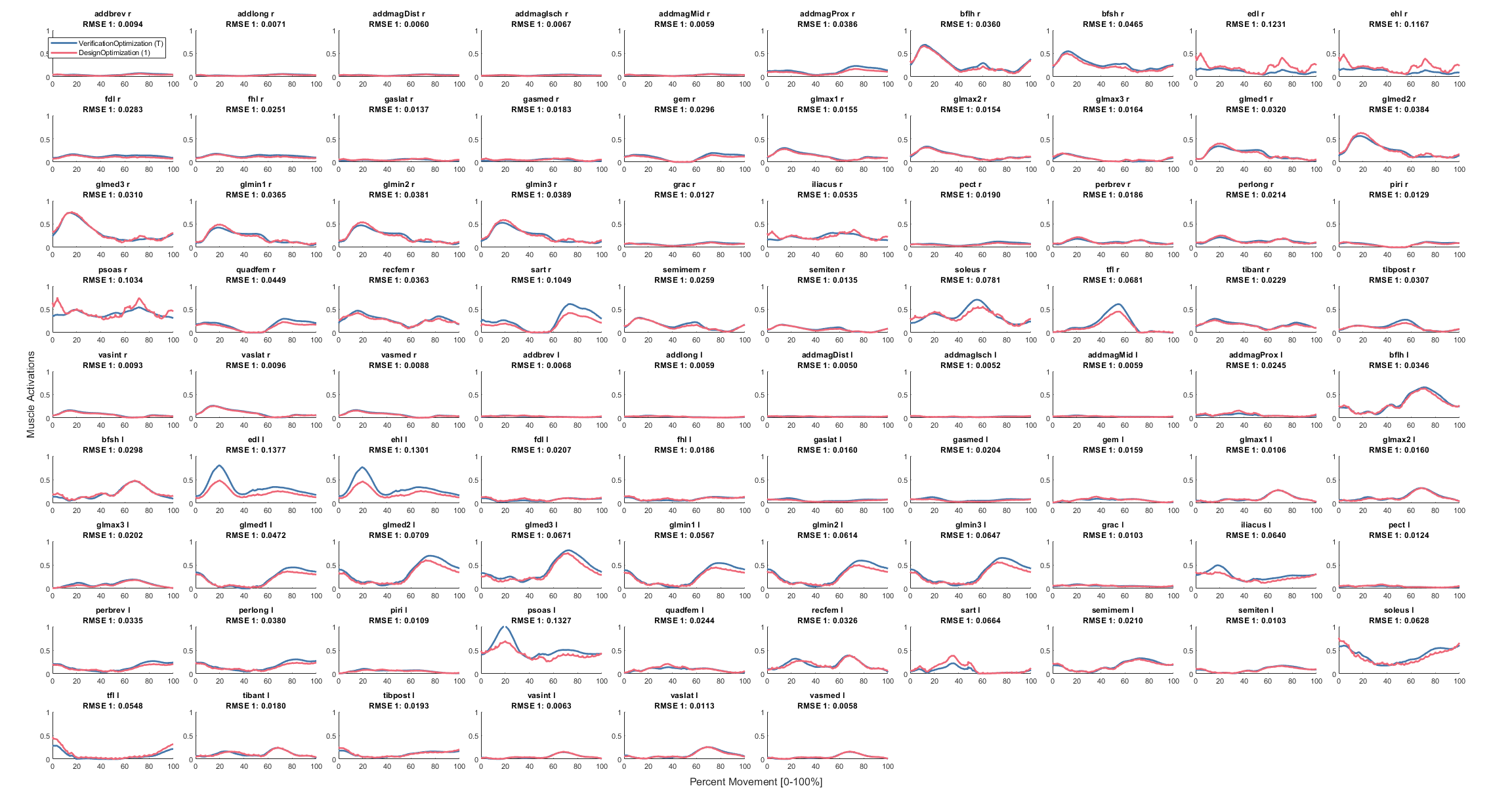

### rightActivation.png

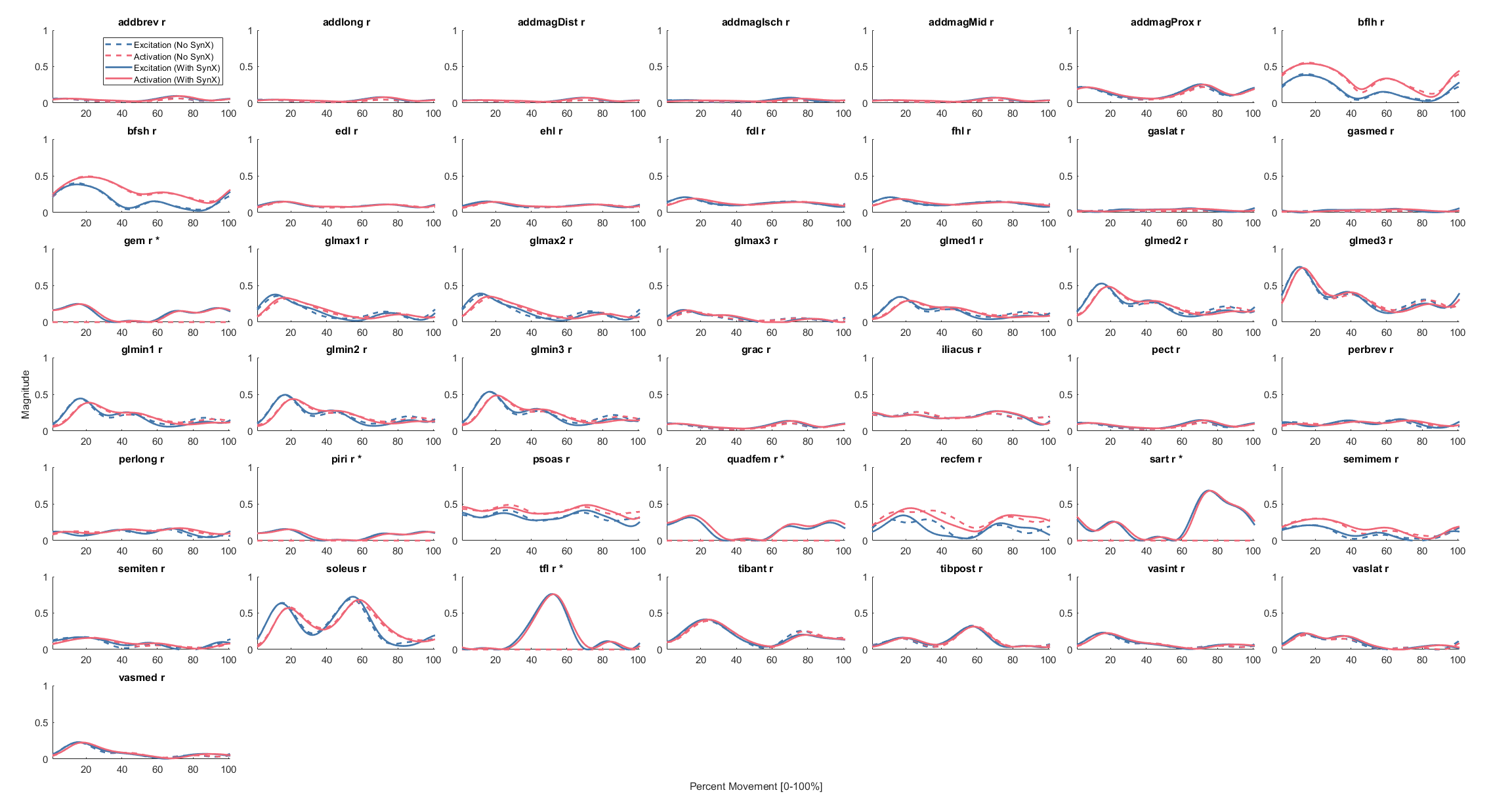

### rightFiberLength.png

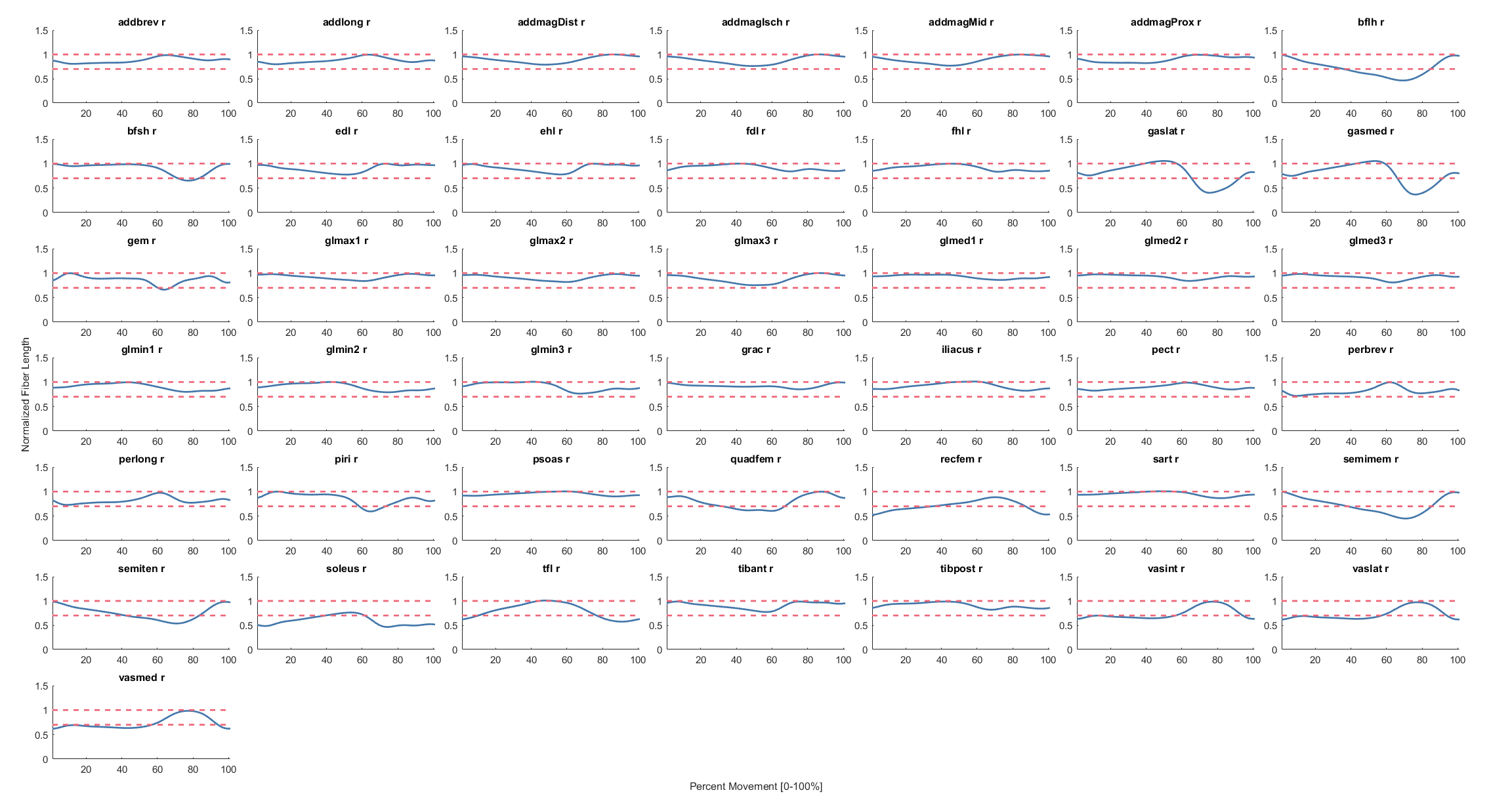

### rightMoment.png

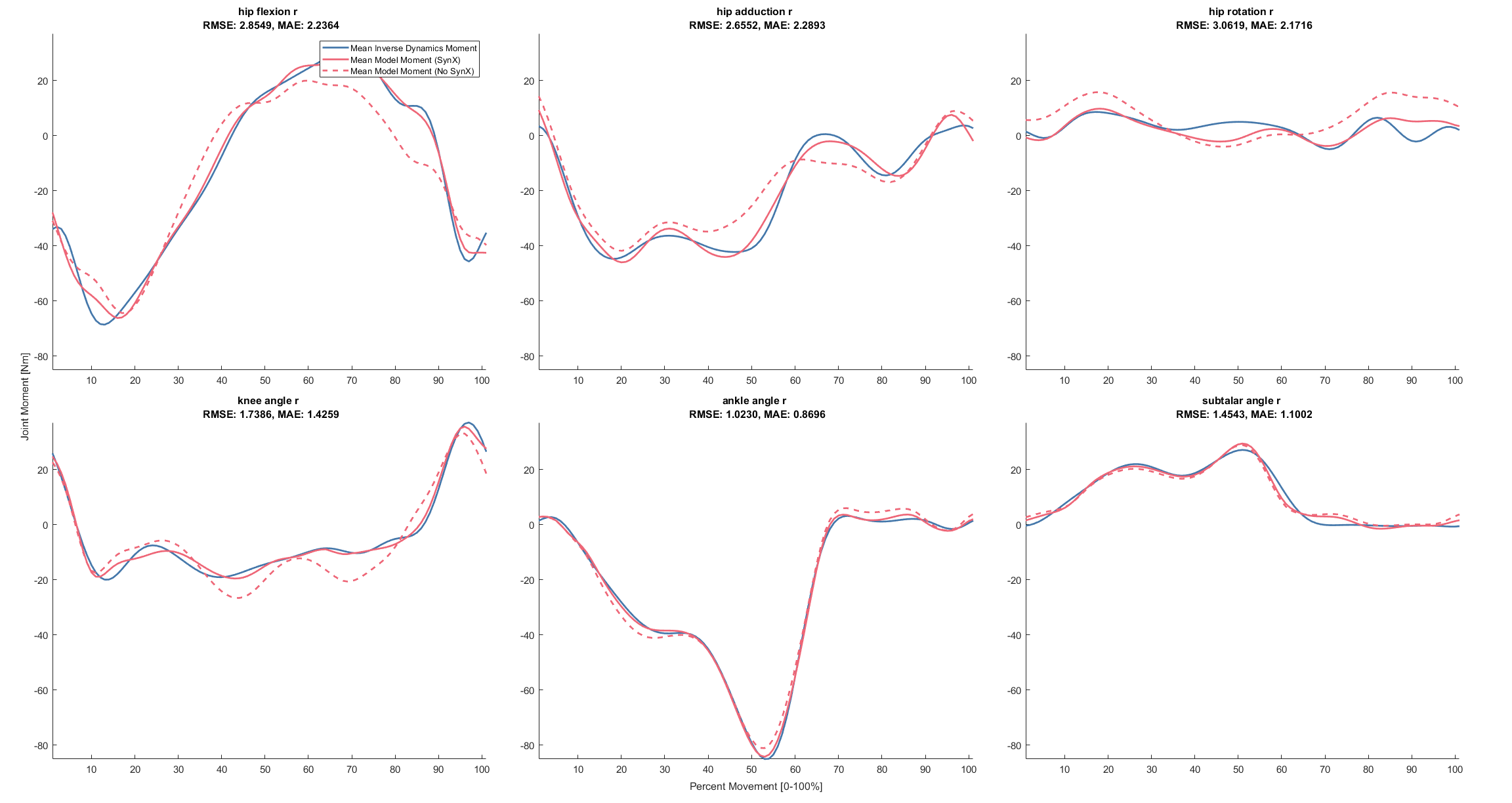

### rightParameters.png

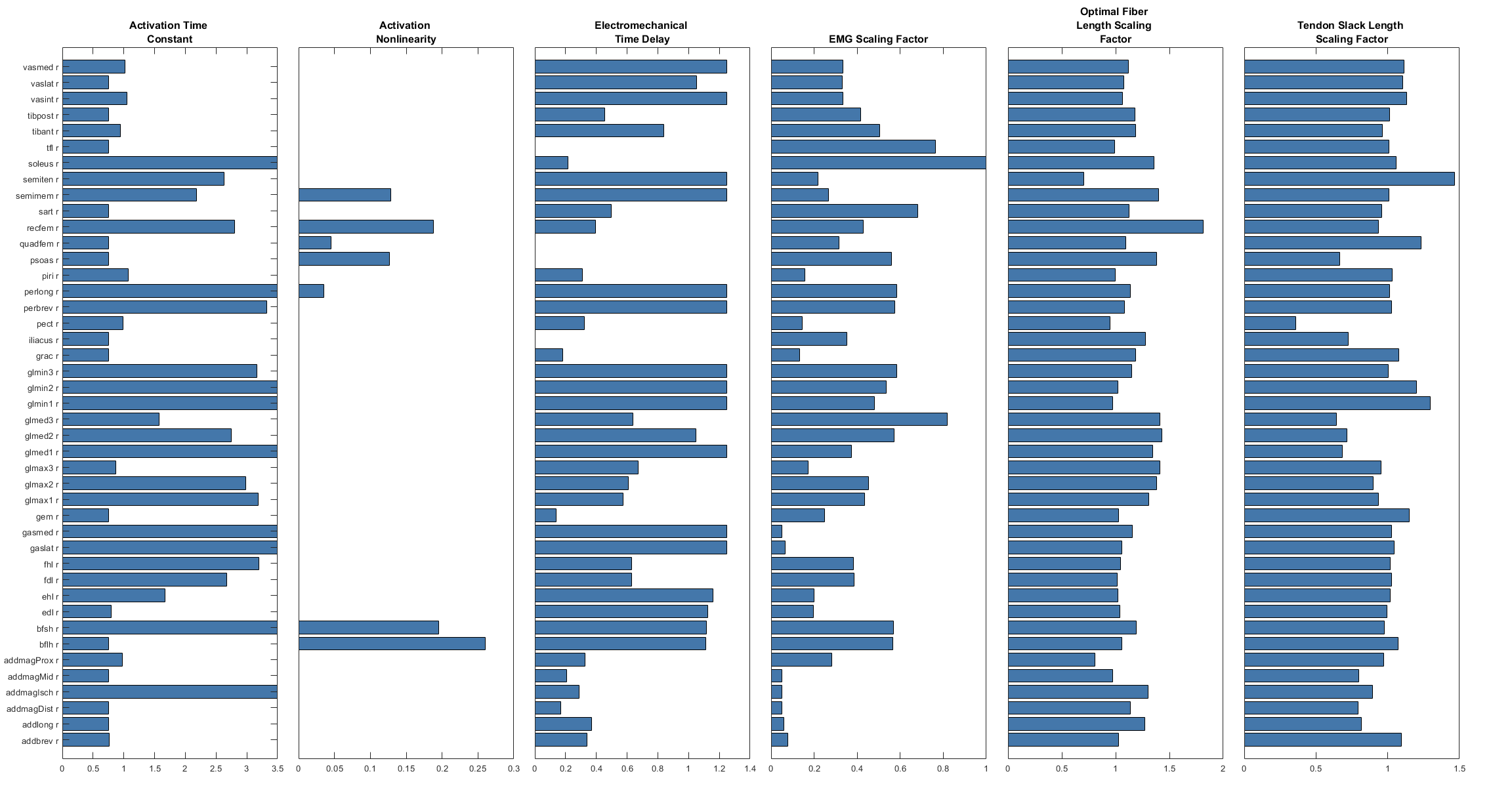

### rightPassiveForce.png

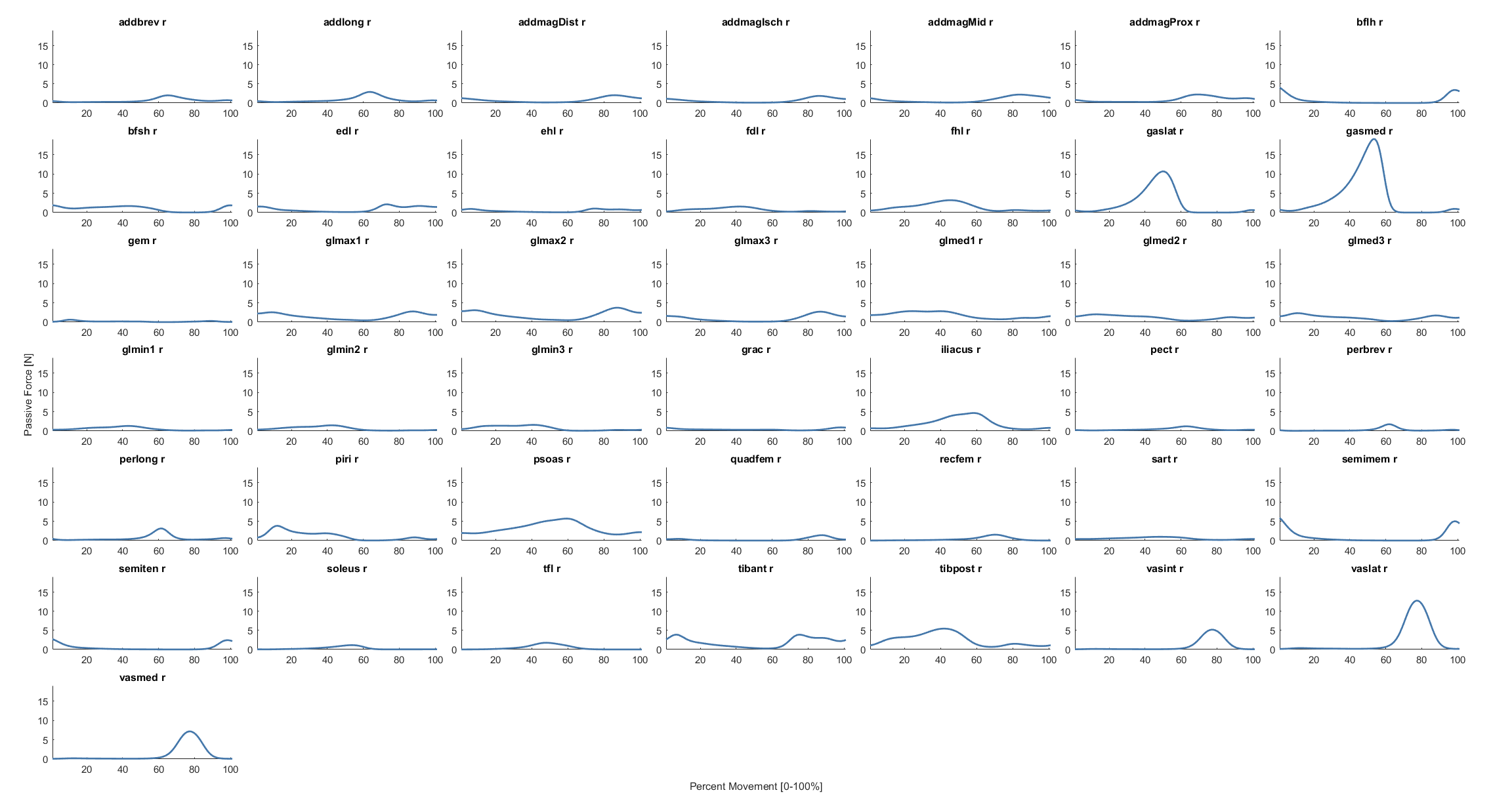

### rightPassiveMoment.png

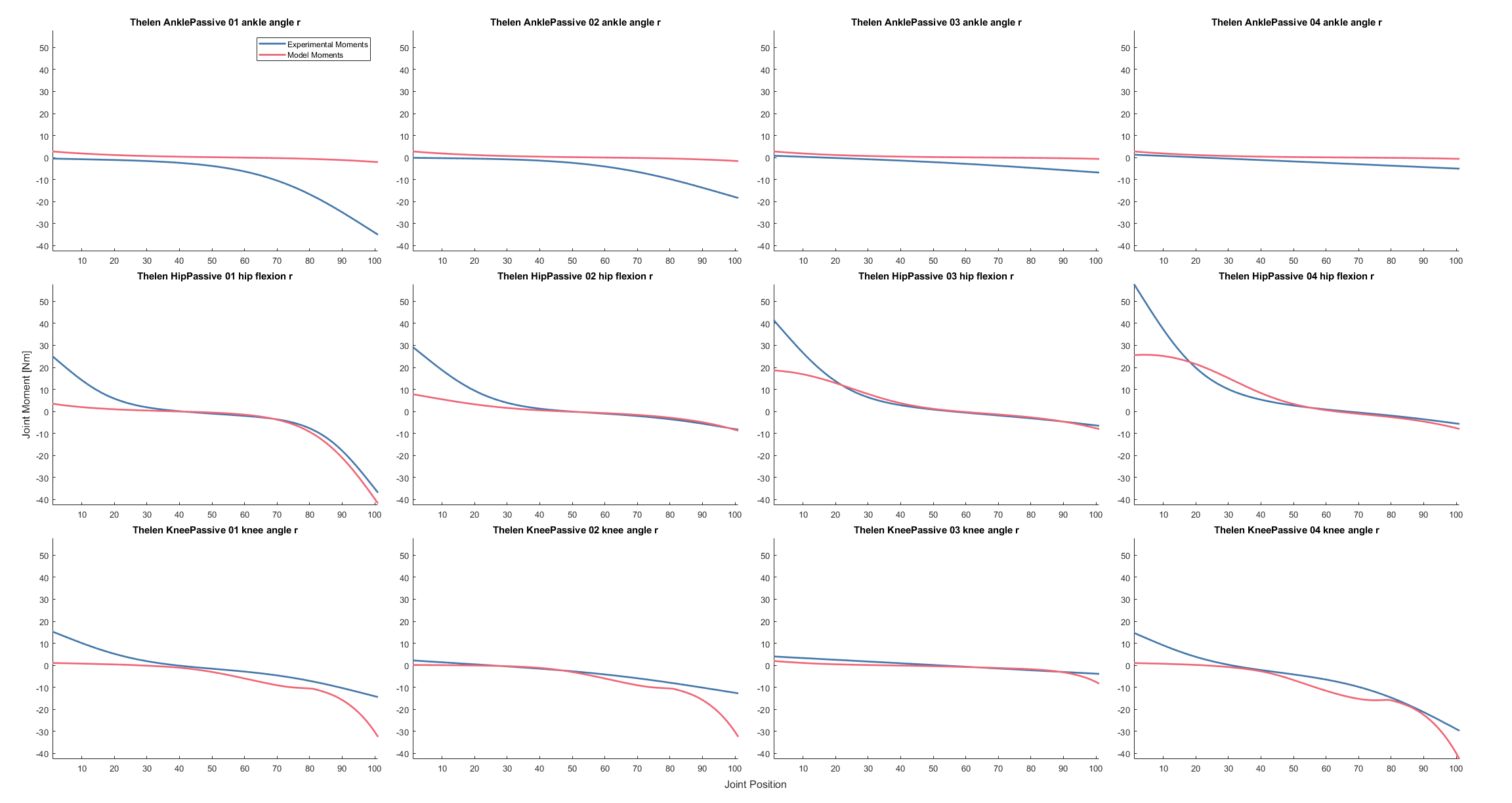

### stiffnessCoefficients.png

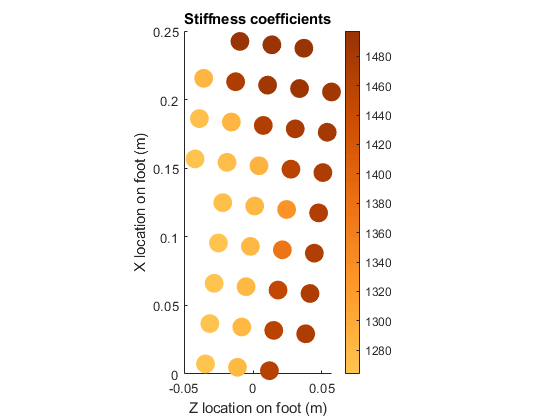

### synergyControls.png

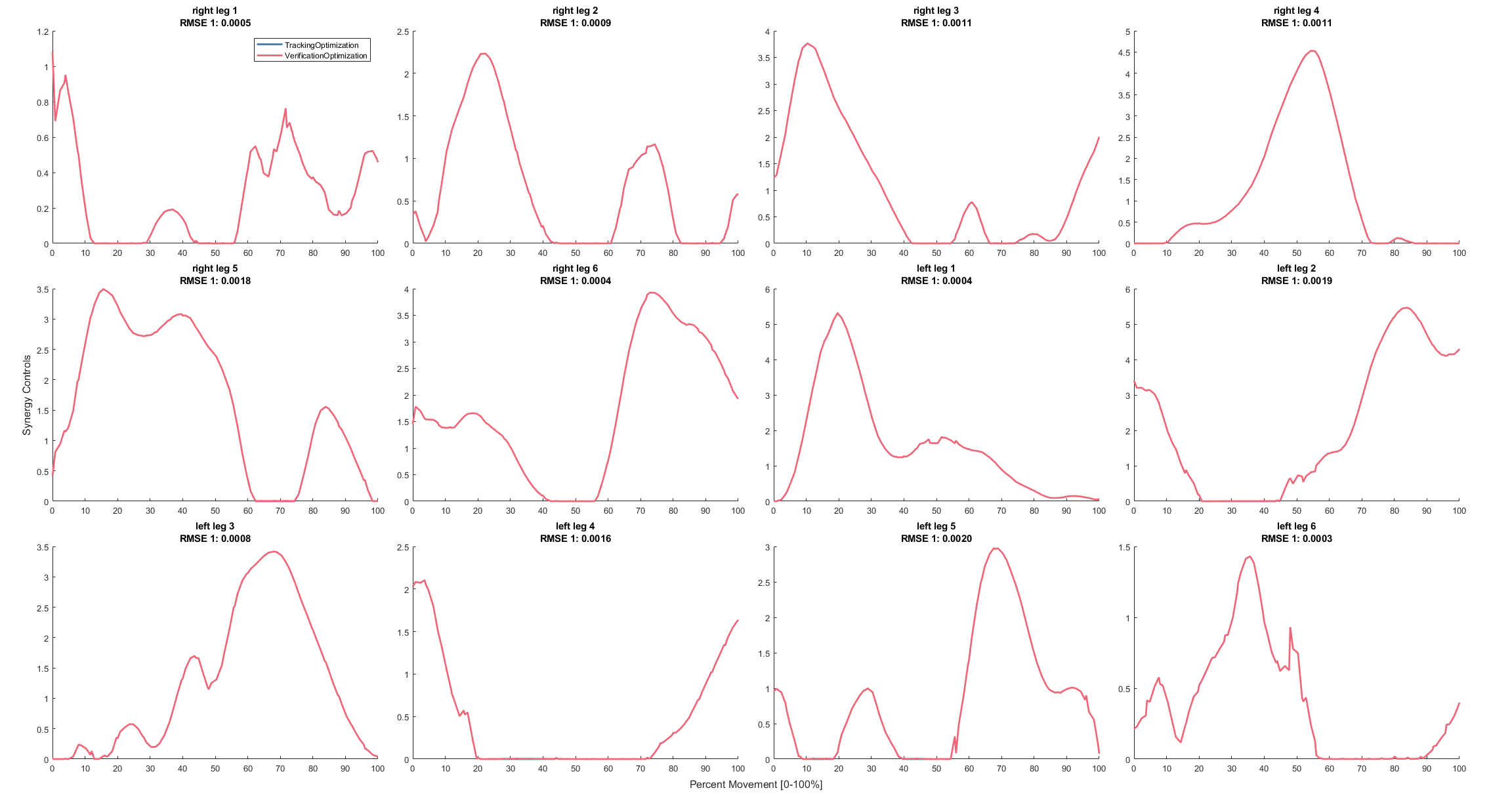

### synergyControls.png

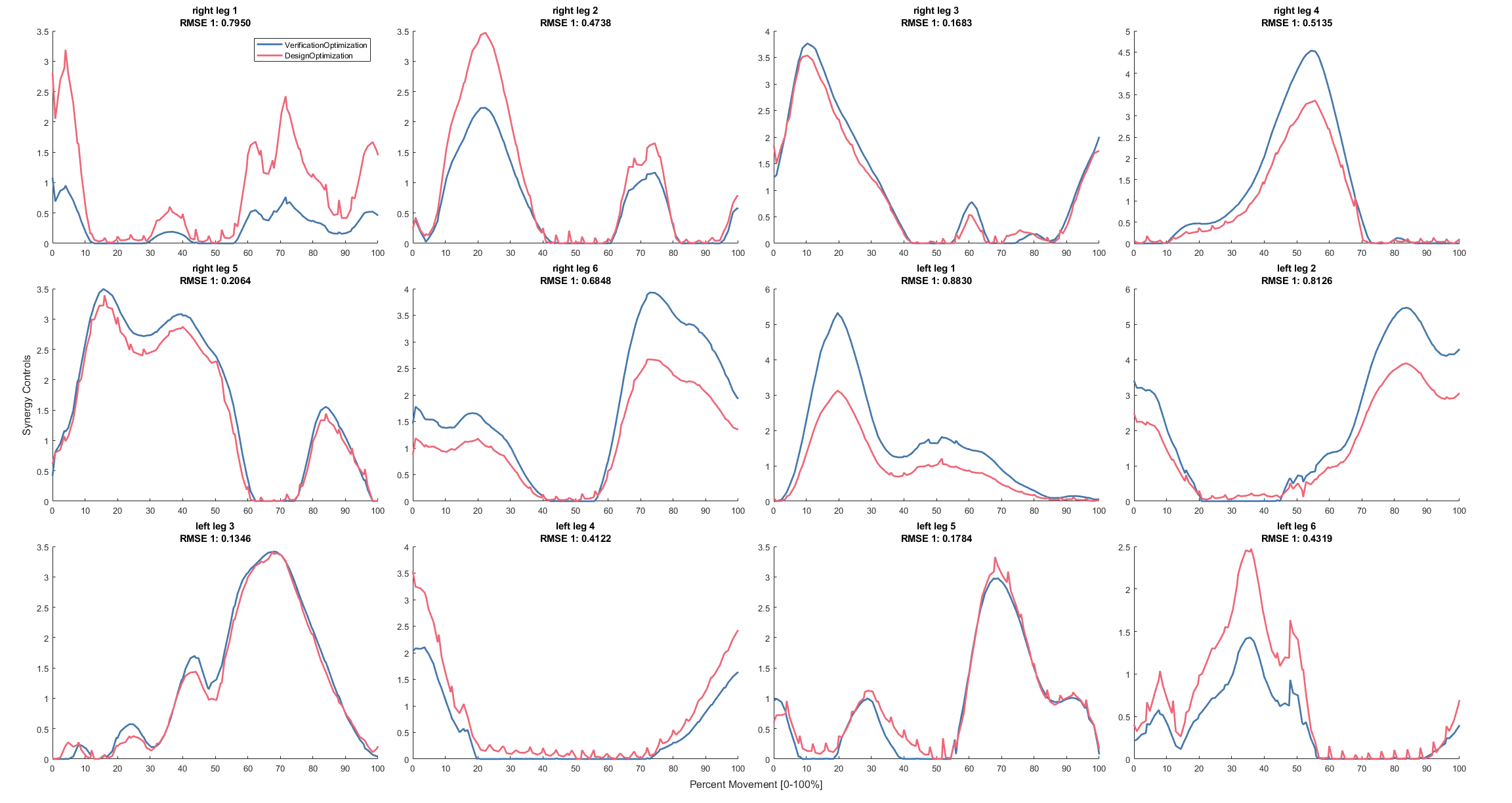

### synergyControls.png

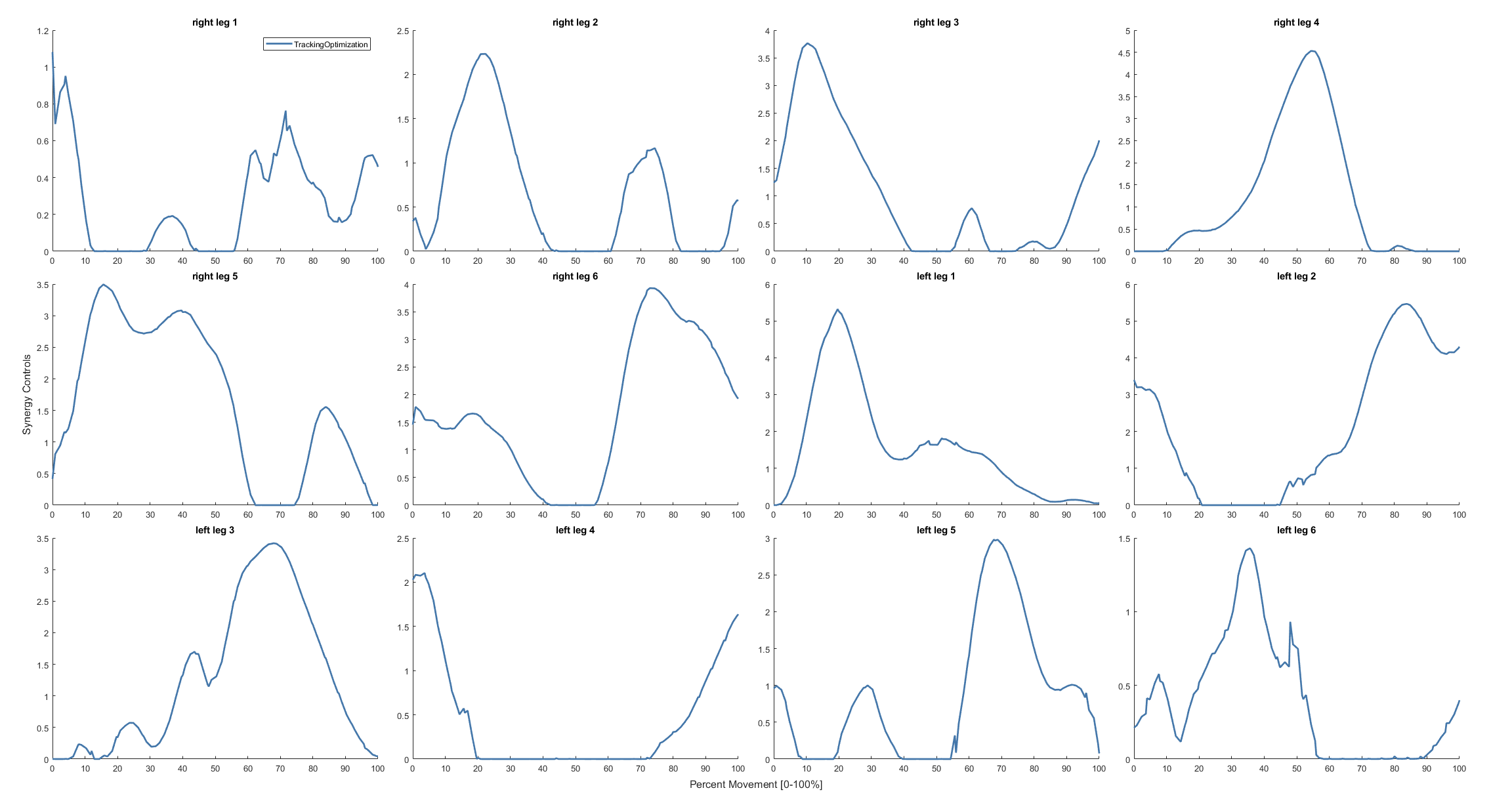

### torqueControls.png

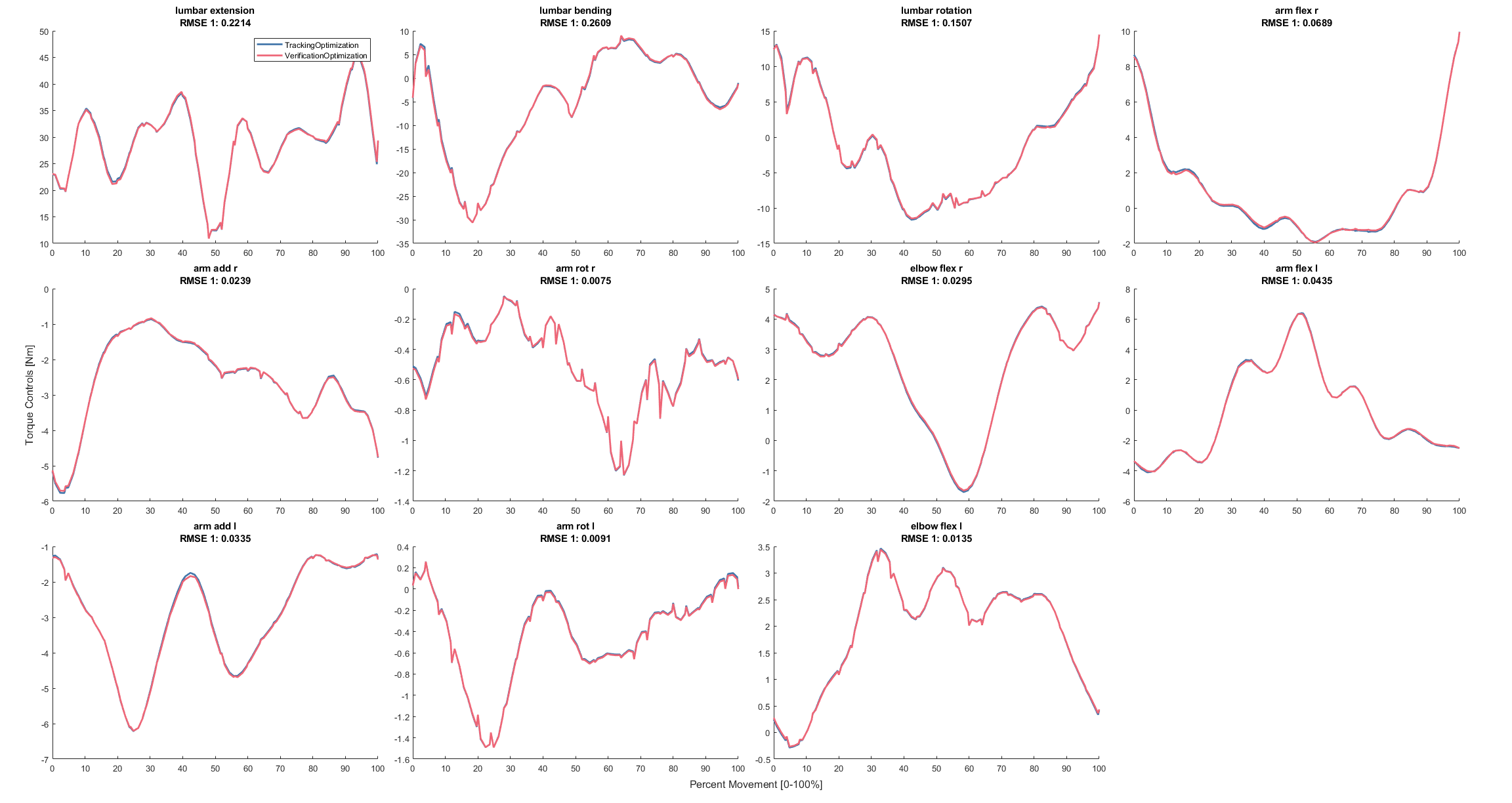

### torqueControls.png

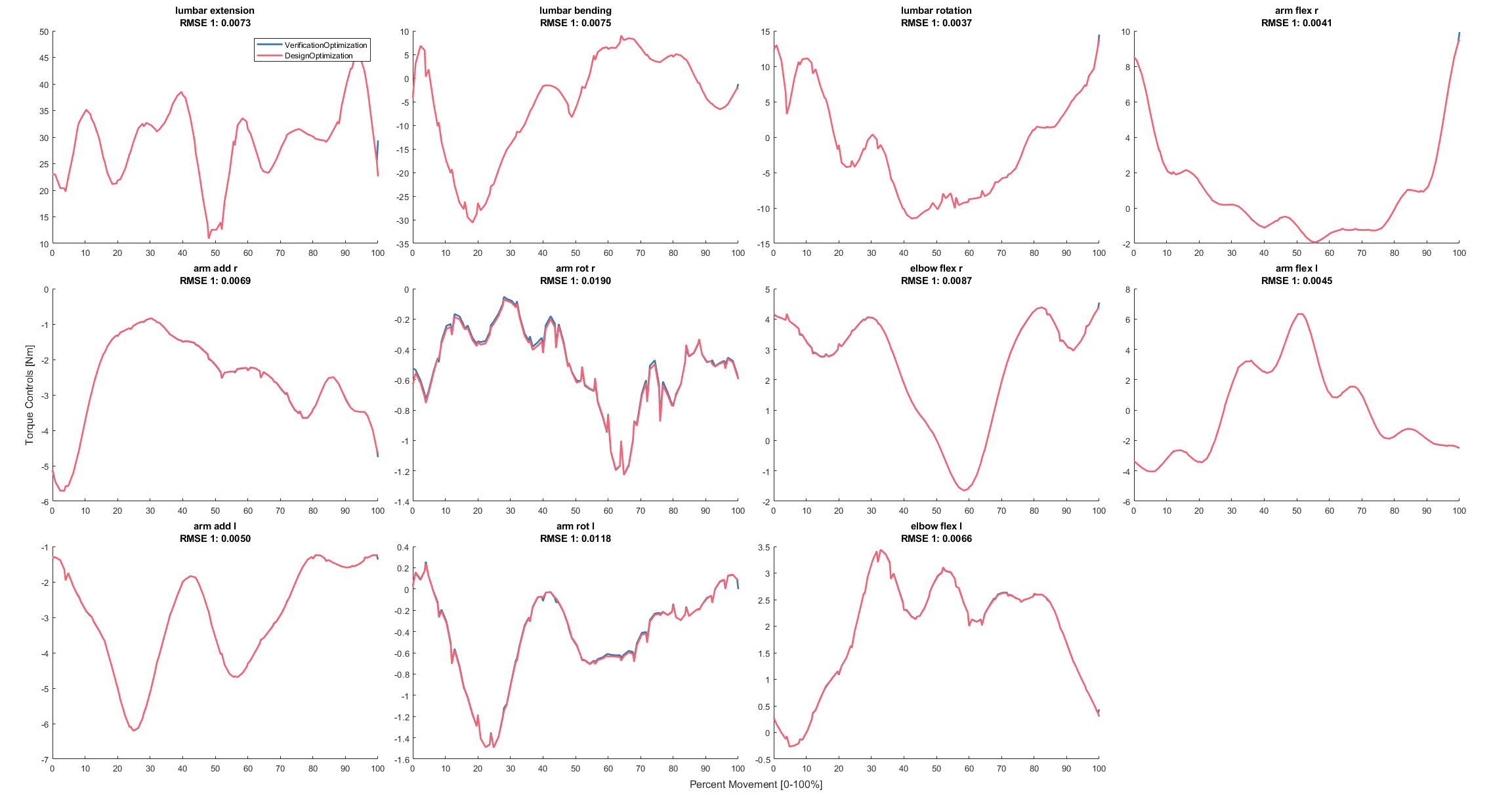

### torqueControls.png

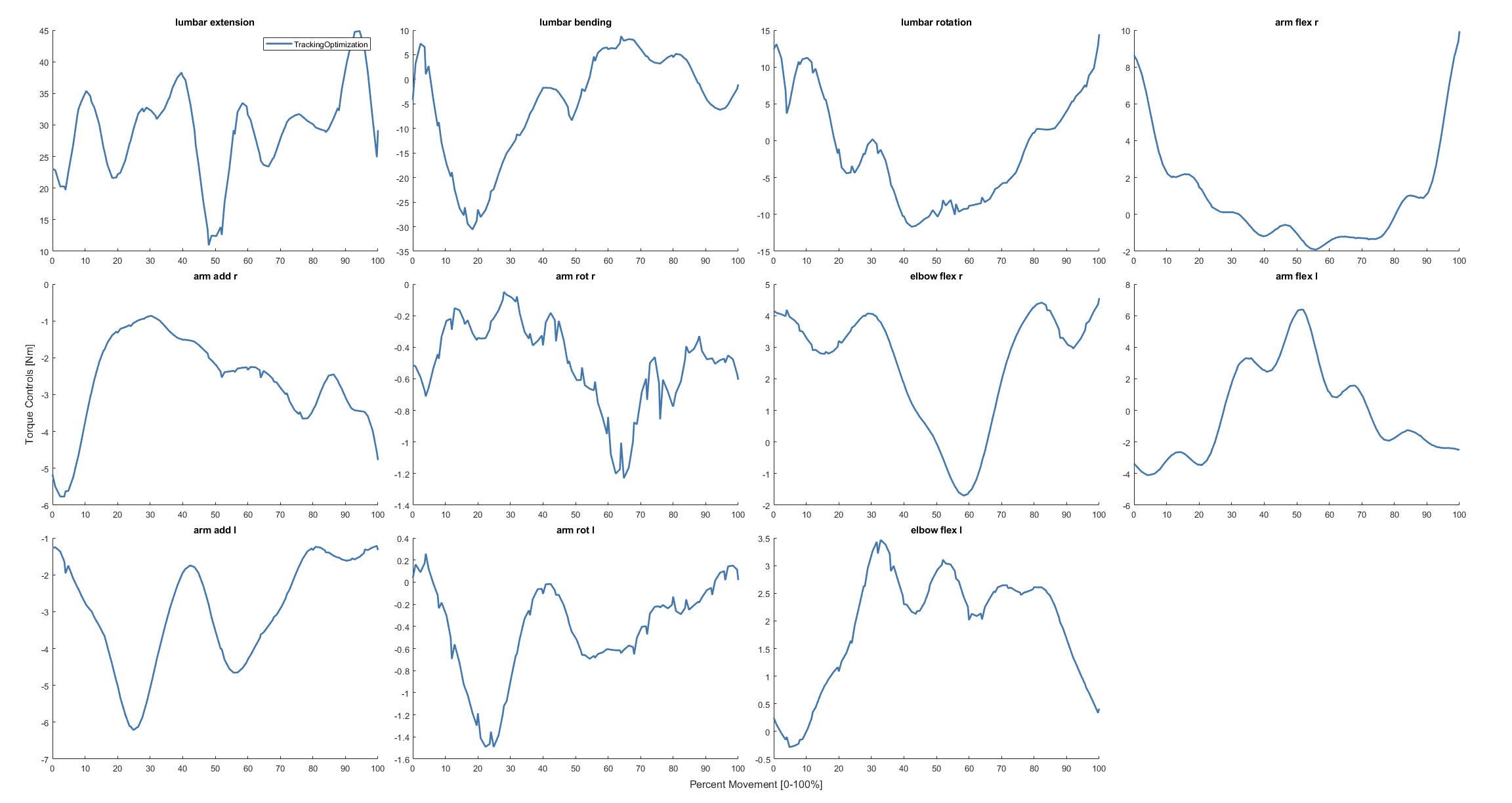

### VAF.png

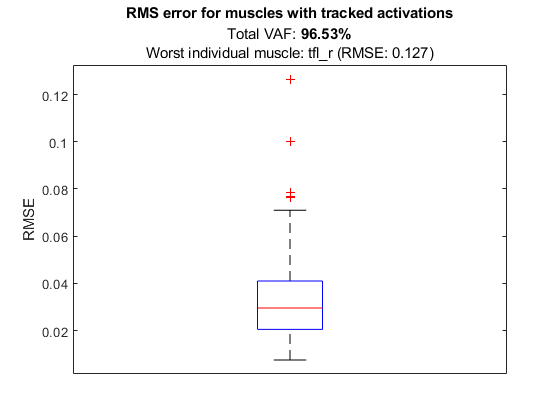
